# The Genomic Diversity of the *Eliurus* genus in northern Madagascar with a Putative New Species

**DOI:** 10.1101/2022.10.21.513246

**Authors:** Gabriele Maria Sgarlata, Emmanuel Rasolondraibe, Jordi Salmona, Barbara Le Pors, Tantely Ralantoharijaona, Ando Rakotonanahary, Fabien Jan, Sophie Manzi, Amaya Iribar-Pelozuelo, John Rigobert Zaonarivelo, Nicole Volasoa Andriaholinirina, Solofonirina Rasoloharijaona, Lounès Chikhi

**Author notes:** **<u>Corresponding authors:</u> Gabriele Maria Sgarlata** - Instituto Gulbenkian de Ciência, Rua da Quinta Grande, 6, 2780-156 Oeiras, Portugal - -, **Lounès Chikhi** - Instituto Gulbenkian de Ciência, Rua da Quinta Grande, 6, 2780-156 Oeiras, Portugal.

## Abstract

Madagascar exhibits extraordinarily high level of species richness and endemism, while being severely threatened by habitat loss and fragmentation (HL&F). In front of such threat to biodiversity, conservation effort can be directed, for instance, in the documentation of species that are still unknown to science, or in investigating how species respond to HL&F. The tufted-tail rats genus (*Eliurus* spp.) is the most speciose genus of endemic rodents in Madagascar, with 13 described species, which occupy two major habitat types: dry or humid forests. The large species diversity and association to specific habitat types make the *Eliurus* genus a suitable model for investigating species adaptation to new environments, as well as response to HL&F (dry *vs* humid). In the present study, we investigated *Eliurus* spp. genomic diversity across northern Madagascar, a region covered by both dry and humid fragmented forests. From the mitochondrial DNA (mtDNA) and nuclear genomic (RAD-seq) data of 124 *Eliurus* individuals sampled in poorly studied forests of northern Madagascar, we identified an undescribed *Eliurus* taxon (*Eliurus sp. nova*). We tested the hypothesis of a new *Eliurus* species using several approaches: i) DNA barcoding; ii) phylogenetic inferences; iii) species delimitation tests based on the Multi-Species Coalescent (MSC) model, iv) genealogical discordance index (*gdi*); v) the *ad-hoc* test of isolation-by-distance within *versus* between sister-taxa, vi) comparisons of %GC content patterns and vii) morphological analyses. All analyses support the recognition of the undescribed lineage as a distinct species. In addition, we show that *Eliurus myoxinus*, a species known from the dry forests of western Madagascar, is, surprisingly, found mostly in humid forests in northern Madagascar. In conclusion, we discuss the implications of such findings in the context of *Eliurus* species evolution and diversification, and use the distribution of northern *Eliurus* species as a proxy for reconstructing past changes in forest cover and vegetation type in northern Madagascar.

---

The endemic rodents of Madagascar (subfamily *Nesomyinae*) are a diverse taxonomic group of the family Muridae, whose morphological diversity has led to the recognition of nine genera (Musser and Carleton 2005) characterised by diverse life-history traits, such as scansorial, semi-fossorial or saltatorial behaviours. Their monophyly is now recognised, although debated over decades due to the lack of sufficiently powerful data and the morphological similarity with non-Malagasy rodents (Jansa et al. 1999; Michaux et al. 2001; Jansa and Weksler, 2004). Genetic data have established that Malagasy rodents came from Africa during a single colonization event (Steppan and Schenk 2017; Jansa and Carleton, in press), which has been dated between 15 and 30 Ma by Poux et al. (2005) and between 12.8 and 15.6 Ma by Steppan and Schenk (2017). Since then, the *Nesomyinae* have diversified across the various ecotypes of Madagascar.

Most *Nesomyinae* genera are composed of few species (either one or four), with the exception of the *Eliurus* genus which, with 13 described species, is by far the most speciose genus of *Nesomyinae* (Jansa et al. 2019). The *Eliurus* genus was originally characterised by Alphonse Milne-Edwards in 1885 (Milne Edwards, 1885), with the description of *E. myoxinus* in western Madagascar. During the following decades, new *Eliurus* species were mainly described from the humid forests of eastern Madagascar (Carleton 1994; Carleton and Goodman 1998; Carleton 2003). Recently, four *Eliurus* species from the west and north of Madagascar have been added to the genus (Carleton et al. 2001; Goodman et al. 2009; Jansa et al. 2019). In the last twenty-five years, field surveys, museum-specimen based research and genetic data (although limited to two mtDNA and four nuclear loci) have greatly contributed to a better understanding of endemic rodent biodiversity in Madagascar (Carleton 1994; Jansa et al. 1999; Goodman and Rasolonandrasana 1999; Carleton et al. 2001; Jansa and Carleton 2003; Carleton and Goodman 2007; Goodman et al. 2007, 2009; Goodman et al. 2013a; Tysdal and Jansa 2014; Jansa et al. 2019), leading to a seven-fold increase in *Eliurus* specimen collection and the description of five new species (see Fig.1 in Jansa et al. 2019).

**Figure 1.**
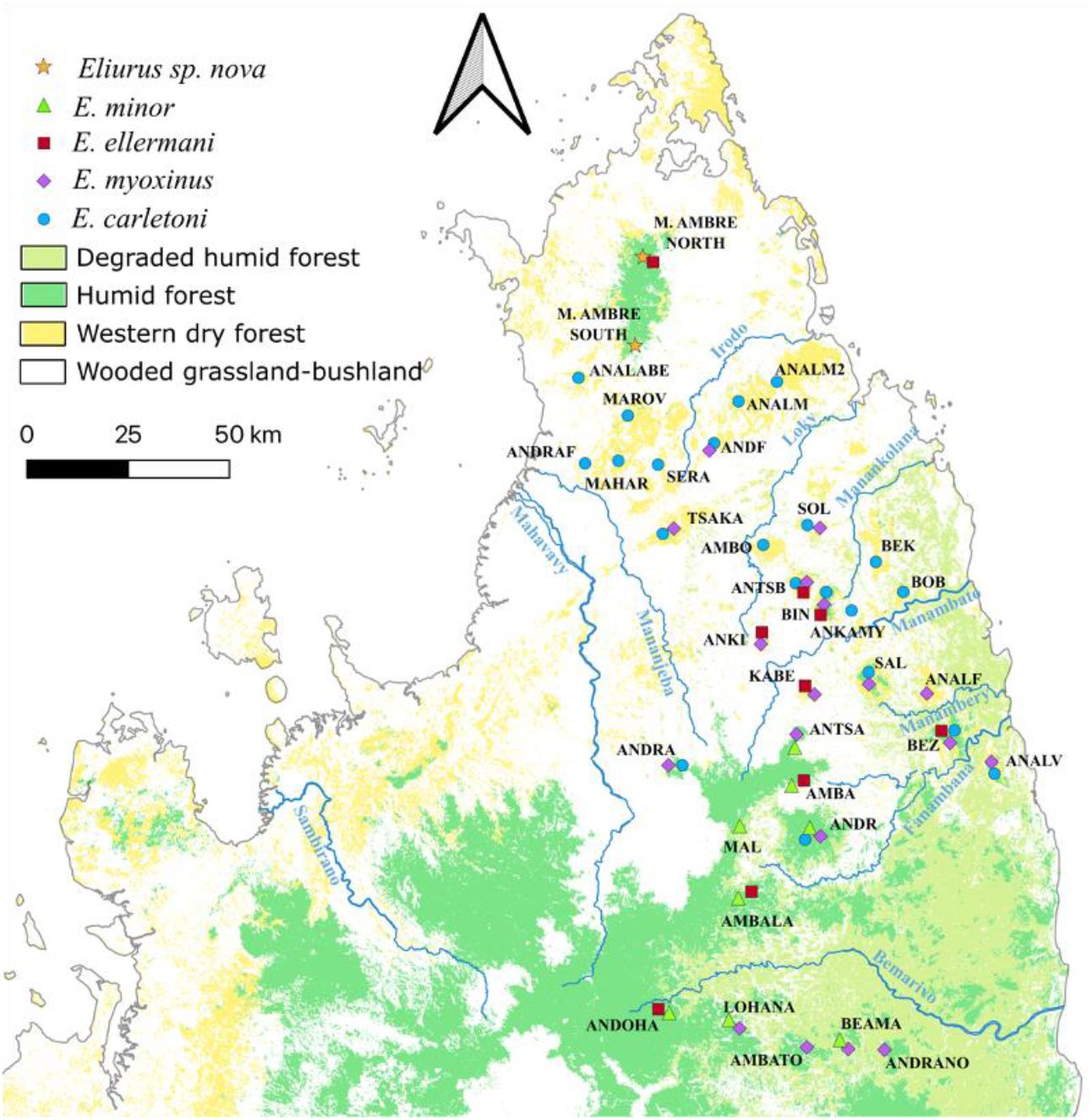
*Eliurus* individuals sampled in northern Madagascar. In this figure we present the sampling sites of the five *Eliurus* taxa identified in the present study: *E. myoxinus*; *E. carletoni*; *E. minor*; *E. ellermani* and *Eliurus* sp. nova. Detailed map for each species can be found in Supplementary Appendix S3: Figure S20.

While the use of genetic data may be problematic, as it has led to a likely inflation in the number of species on other Malagasy mammals, such as mouse lemurs and sportive lemurs (e.g., Tattersall 2007; Tattersall and Cuozzo 2019), morphological data may not always be sufficient to identify species. This is the case when phenotypic differences are small, difficult to quantify or the result of limited sampling, even when divergence between sister-species is not recent (Bickford et al. 2007; Struck et al. 2018; Chenuil et al. 2019). For instance, one of the diagnostic morphological characters that Carleton (1994) proposed for distinguishing *Eliurus ellermani* from *Eliurus tanala* was a ‘tail brush completely dark to the tip’. However, the recent study of Jansa et al. (2019), which integrated morphological and genetic data from a larger number of specimens, showed that individuals genetically identified as *Eliurus ellermani* had in most cases a white tail tuft, and rarely a dark tipped tail. One could also use information on geographic distributions for preliminary taxonomic identification, if species distributions are well known and not overlapping. However, most *Eliurus* species are i) characterised by large and scattered species distributions, and ii) often in co-occurrence or syntopy with one or more *Eliurus* species (Carleton 1994; Marquart 2014; Jansa et al. 2019).

In the present study, we sampled 124 *Eliurus* individuals from northern Madagascar, a transition region between the eastern humid and the western dry ecoregions. We then assessed their mitochondrial (mtDNA) and nuclear genomic diversity. With the use of a Restriction site-associated DNA sequencing (RAD-seq) approach, which generates a reduced but wide representation of genomic diversity, we provided the first multispecies genome-wide dataset for the *Eliurus* genus, aiming at validating and clarifying the *Eliurus* species tree topology. We performed several phylogenetic inferences and species delimitation tests, taking into account genomic and geographic variability within each taxon, as well as gene tree discordance. In addition, we quantified morphological diversity across *Eliurus* species and correlate such diversity with bioclimatic variables to test for support of Bergman’s rule. In conclusion, our analyses identify a new monophyletic and divergent lineage which we hypothesize to be a new *Eliurus* species.

## Materials and Methods

### Study Region

We surveyed 35 forest sites between 2010 and 2018 in northern Madagascar, a region considered to be a transition zone between Madagascar’s eastern-humid and western-dry ecoregions. The landscape of northern Madagascar is characterized by forest fragments of diverse habitat types (dry deciduous, transition, humid evergreen, mountainous) separated by a matrix of savannah, grassland and agricultural lands. This region is crossed by six main rivers, the Irodo, the Loky, the Manambato, the Manambery, the Fanambana and the Bemarivo rivers, which further divide northern Madagascar in inter-river systems: North of the Loky river, Loky-Manambato region, South Manambato region, COMATSA region, North Marojejy (Fig. 1). A detailed description of each inter-river system can be found in Supplementary Appendix S1.

### Study Species

The tuft-tailed rats (genus *Eliurus*) are nocturnal and easily recognizable by a noticeable tuft of long hairs over the distal portion of the tail. They are associated with forested habitats, but some species show tolerance to mild forest disturbance. In particular, trap captures data provided evidence on the scansorial, arboreal and terrestrial behaviour of this group of rodents, since individuals can be both collected on lianas, trees and on the ground (e.g., Marquart 2014). However, there are some distinctions that can be made among species. For instance, Webb (1954) observed that western *Eliurus* species are more arboreal than eastern species, rarely descending to the ground. In the present work, we focus on four *Eliurus* species that have been described in northern and north-eastern Madagascar: *Eliurus carletoni, Eliurus ellermani, Eliurus minor* and *Eliurus myoxinus* (Fig. 1). Details for each described species can be found in Supplementary Appendix S1.

### Sampling and Laboratory Procedures

We collected small ear biopsies from 124 individuals captured with Sherman traps (H.B. Sherman Traps®) (Table 1), according to Rakotondravony and Radespiel (2009) field procedures. We ensured DNA preservation in field conditions by storing the samples in Queen’s lysis buffer (Seutin et al. 1991) and at −20 °C once in the laboratory. We extracted total genomic DNA using a DNeasy® blood and tissue kit (Qiagen®). To increase the amount of extracted genomic DNA, tissues were incubated overnight at 56 °C in 300 μL of ATL digestion buffer (Qiagen®) including proteinase *K* (Cf = 1 mg/ mL), and 20 μL of dithiothreitol (1 M). The final DNA solution was then eluted in 60 μL of AE buffer. We quantified nucleic acids and checked for DNA purity using the Nanodrop spectrophotometer (ND-1000, Thermo Fisher) and quantified double stranded DNA (dsDNA) using a Qubit 2.0 Fluorometer (Invitrogen). For maximizing the amount of genomic data recoverable from high-throughput DNA sequencing, we selected samples with a dsDNA concentration superior to 4 ng/μL and a 260/280 absorbance ratio higher than 1.75.

**Table 1.**
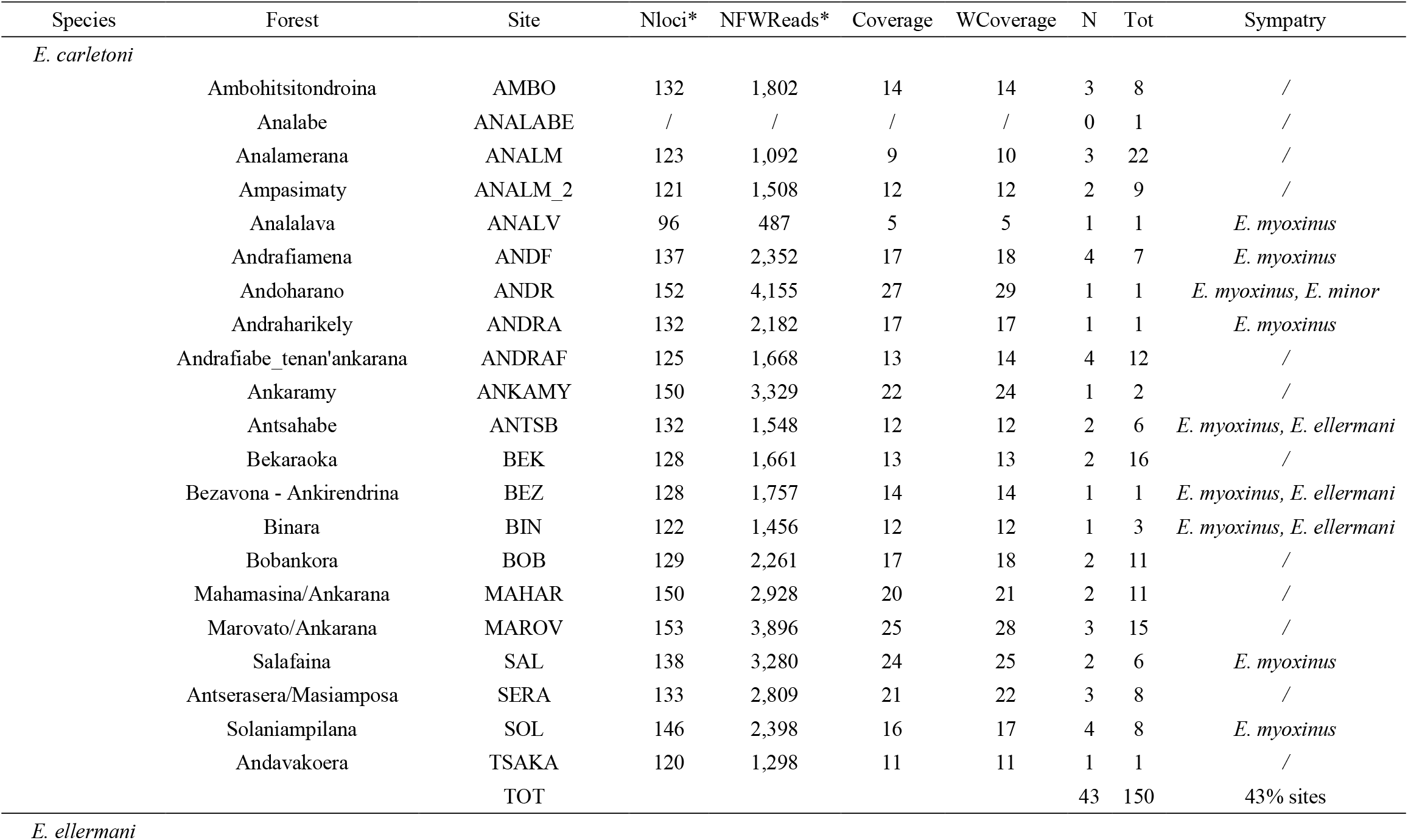
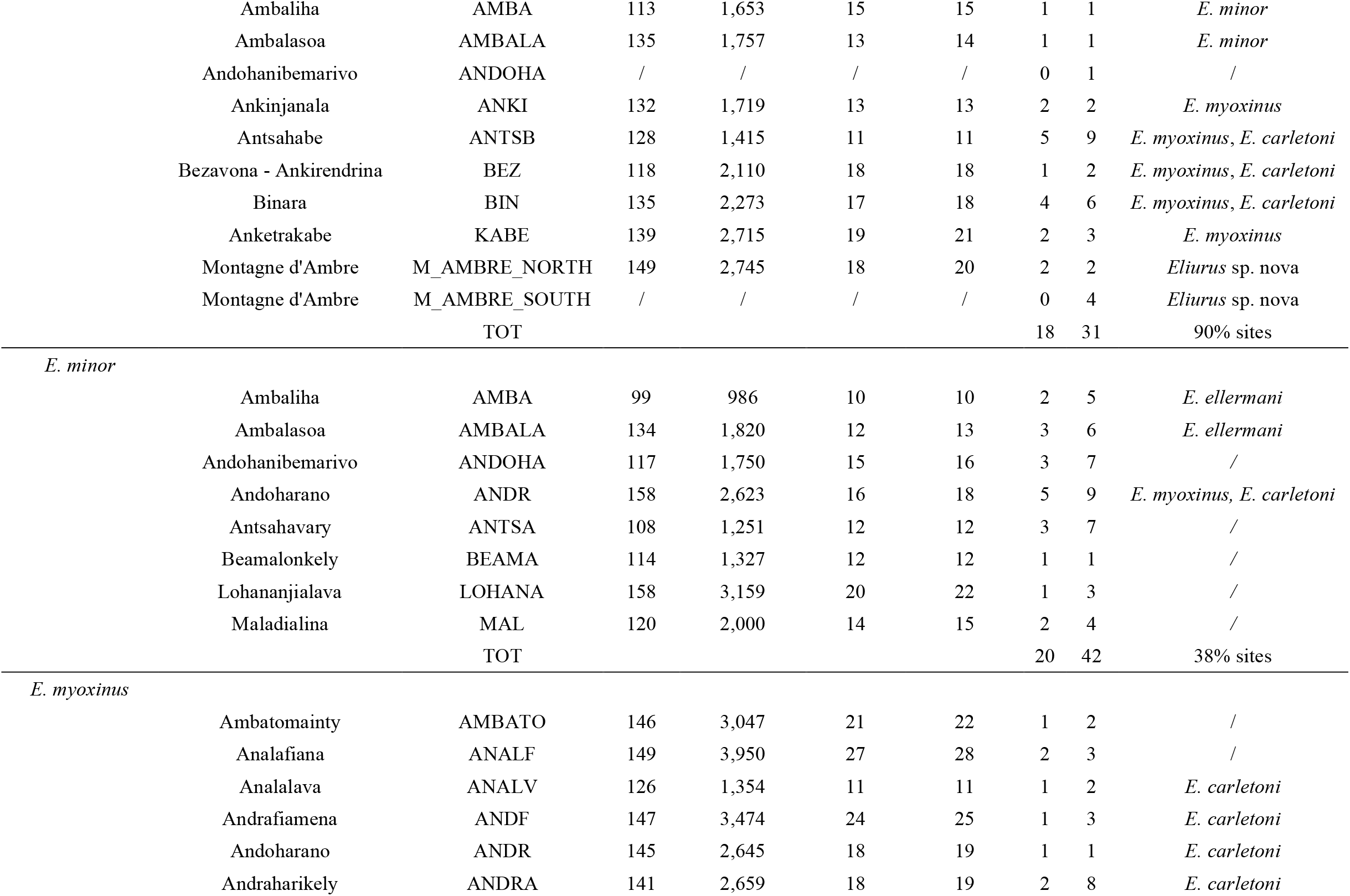
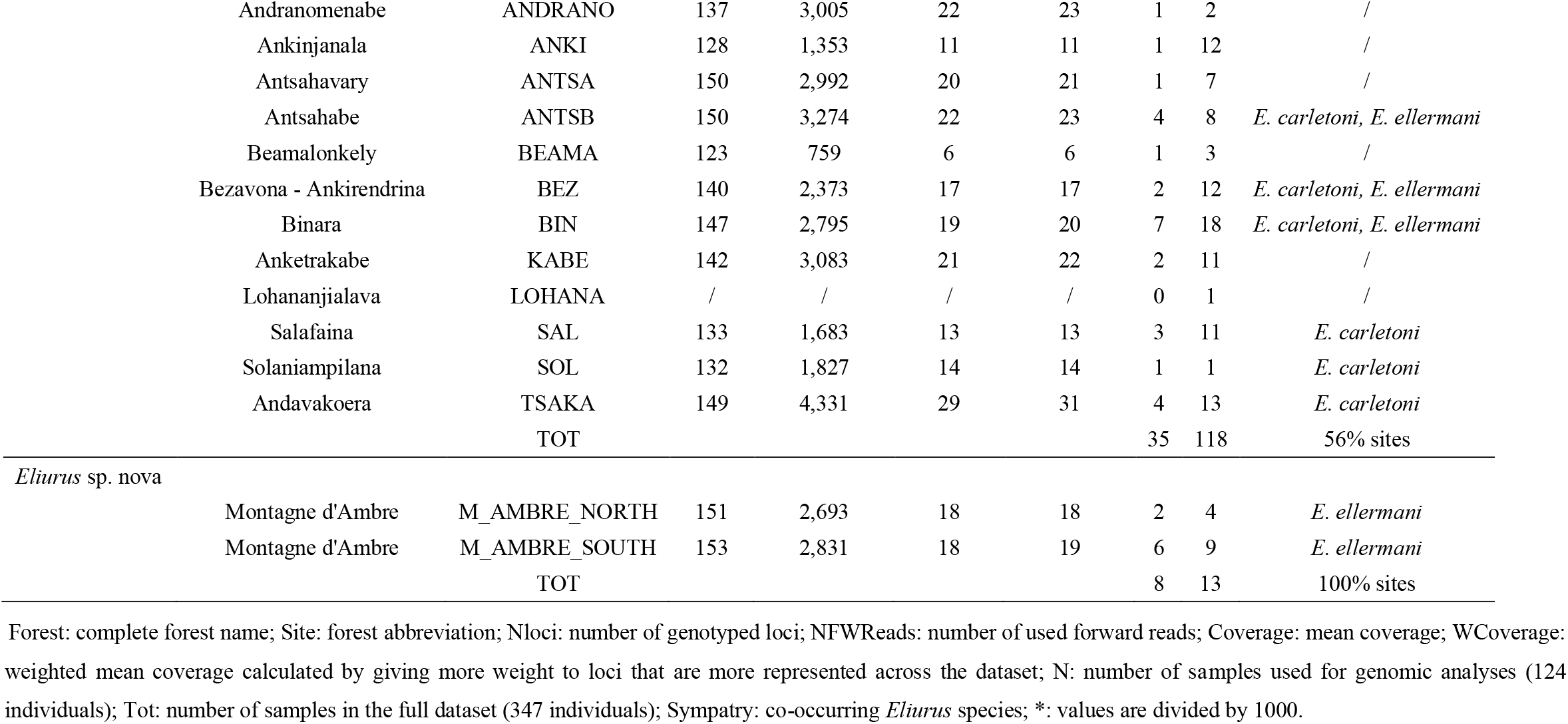
Localities and summary statistics of nuclear genomic data of the *Eliurus* individuals sampled across northern Madagascar.

We amplified and sequenced the mitochondrial (mtDNA) cytochrome b (*cytb*; 1140 bp) of 20 individuals, distributed in the four clades identified with preliminary analyses of the nuclear RAD-seq data (see Results). The mtDNA analyses aimed at clarifying species identity, based on previous mtDNA phylogenetic studies, since *Eliurus* species can be difficult to distinguish morphologically and since there are no published genomic data to compare our data with. mtDNA *cytb* was amplified using published primers: MVZ05 and UMMZ04 (Jansa et al. 1999). We carried out PCR amplification in a 10-μL reaction with final concentrations of 0.8 μl of 1.0 μM for each primer, 5 μl of 1.5 mM GoTaqFlexi DNA polymerase Mix (Promega no. M8305), 1 μl of genomic DNA and 2.4 μl of H_2_O. Thermocycling consisted of an initial denaturing temperature of 94 °C for 2 min, followed by 35 cycles of 1 min at 94 °C, 1 min at 51 °C (*cytb* annealing temperature) and 1 min at 72 °C, and a final extension of 10 min at 72 °C. We sequenced the PCR products in an ABI 3130 XL Genetic Analyzer (Applied Biosystems, Foster City, CA) and edited the sequences using GENEIOUS PRO v10.2.3 (http://www.geneious.com, Kearse et al. 2012).

### RAD-sequencing

Restriction site-Associated DNA sequencing (RAD-seq) consists in sequencing regions neighbouring restriction sites, to obtain homologous sequences across many individuals, spread across the genome, with high coverage and at a reasonable cost (Baird et al. 2008; Andrews et al. 2016). We prepared RAD-seq libraries using 100-200 ng of genomic DNA from each sample and one negative control made up of water following the protocol described in Etter et al. (2011). Briefly, samples were digested with SbfI (New England Biolabs) before P1 adapter ligation. After adapter ligation, we pooled 10 μL of 18 samples in four sub-libraries, before shearing them in a Covaris M220 for 45 sec to obtain fragments with an average size of 500 bp. Following end repair, we conducted size selection using AMPure XP beads (Agencourt). After 3’adenylation, we ligated the P2 adapter. The libraries were finally enriched with 10 cycles of PCR. Libraries quality was verified using Fragment Analyzer® (Advanced Analytical) and quantified using qPCR (QuantStudio6®, Applied Biosystems). We made equimolar pooling with the sub-libraries and sequenced them on an Illumina HiSeq-3000 (150-pb paired-end reads; 72 individuals/lane) at the GetPlage Sequencing platform facility (Toulouse, France). Raw reads were first checked for quality using FastQC (Andrews 2010) and MultiQC (Ewels et al. 2016), then demultiplexed, adapter-removed and quality-filtered in STACKS v2.2 (Rochette et al. 2019) using default parameters. PCR duplicates were removed using the *clone_filter* program in STACKS v2.2 under default settings. The last filtering step was carried out using the *kmer_filter* program in STACKS v2.2 with default settings. This step removes reads with putative sequencing errors (identifying kmers with a coverage lower than 15% of the median read coverage), decrease the number of over-abundant reads and normalize read-depth according to k-mer coverage.

### RAD-seq Data Processing and de novo Assembly

After filtering the raw short-read sequences, it is necessary to assemble the reads pairs into sets of reads (stacks) and merge overlapping stacks into putative loci, which will then be used for population genomic analyses. Since there is no reference-genome available for the *Eliurus* genus, we conducted a *de-novo* assembly of RAD-seq nuclear loci in STACKS v2.2 using *ustacks, cstacks, sstacks, tsv2bam, gstacks* and *populations* programs, which are described in details in Supplementary Appendix S1. The assembly of nuclear loci from RAD-seq data relies on several parameters (*m, M, n*). The choice of the best combination of these parameters can be challenging, since several factors can influence the amount and quality of genomic information recoverable from the dataset, such as the biology and natural history of the studied population(s) or the deployed RAD-seq technique (Paris et al. 2017). Therefore, we followed the protocol proposed by Paris et al. (2017) to optimize the three main parameters of the *de novo* assembly (*m, M, n*). The analysis for the optimization of *de novo* assembly parameters was performed on a subset of the whole dataset, following Rochette and Catchen (2017). In particular, Rochette and Catchen (2017) found that adding all samples does not increase significantly the number of widely shared *catalog* loci in the dataset and can add noise in the estimation of *m, M* and *n*. The individuals used for building the *catalog* were selected by considering i) those with highest coverage and ii) maximizing the geographic distribution of the selected individuals, as a proxy for maximizing the genetic diversity in the dataset. We thus performed the optimization analysis on twenty individuals: five *E. carletoni*, four *E. minor*, five *E. ellermani*, five *E. myoxinus* and one *Eliurus* sp. nova (see Supplementary Appendix S3: Fig. S2 and Supplementary Appendix S2: Table S1).

These were also the individuals used for building the *catalog*. The optimization procedure consisted in iterating a range of values (2 to 8) for the *m, M* and *n* parameters, separately, while keeping all other parameters fixed at default values (*m* = 3, *M* = 2 and *n* = 1). For each parameter combination we extracted the genomic information shared by either 40%, 60% or 80% of the individuals in the dataset. In order to identify the best parameter combination, we extracted and assessed several assembly metrics: i) individual coverage; ii) the number of assembled loci; iii) the number of polymorphic loci; iv) the number of SNPs; v) the variable; and vi) fixed sites identified across the whole population. According to Paris et al., (2017) the best parameter combination is identified considering: i) a stable set of loci that are highly replicated across 80% of the population, since it is unlikely to be derived from paralogous, repetitive sequences or sequencing errors; and ii) the highest amount of polymorphism, or the parameters ‘value at which the number of polymorphisms levels out. In other words, Paris et al. (2017) criteria provide a balance between obtaining true polymorphism and introducing sequencing error, or over-merging paralogous or repetitive loci. Once the informed choice of *m, M* and *n* parameters was done, we re-ran the STACKS v2.2 pipeline using the selected value and obtain the definitive *catalog* on which further genomic analyses were carried out.

### Mitochondrial DNA Analyses

We compared the mtDNA *cytb* of 20 sampled *Eliurus* individuals with published data on the other *Eliurus* species: *E. antsingy* (N= 6), *E. carletoni* (N=67), *E. ellermani* (N = 38), *E. grandidieri* (N = 3), *E. majori* (N = 16), *E. minor* (N = 11), *E. myoxinus* (N = 60), *E. tanala* (N = 14), *E. tsingimbato* (N = 11), *E. webbi* (N = 16) (Supplementary Appendix S2: Table S2, S3). Genetic data were not available for *E. petteri, E. danieli* and *E. penicillatus*. As outgroup we used mtDNA *cytb* of *Rattus norvegicus* (EU349782.1). The complete dataset consisted of 263 mtDNA *cytb* sequences of ∼1064 bp. We aligned and visually checked the sequences using the Clustal Omega method (Sievers et al. 2011). We performed format conversion of DNA alignments using ALTER (Glez-Peña et al. 2010). We used two statistical methods for phylogenetic analysis: a Bayesian MCMC approach implemented in MRBAYES v3.2.1 (Ronquist et al. 2012) and a maximum-likelihood (ML) approach implemented in RAXML v8.2.X (Stamatakis 2014). Information on the parameters used in the mtDNA phylogenetic analyses are described in Supplementary Appendix S1. We measured pairwise mtDNA distance within and between species using the *dist.dna* function in the *adegenet* R package (Jombart 2008; Jombart and Ahmed 2011). We used the ‘raw’ evolutionary model to compute genetic distance, that is the proportion of sites that differ between each pair of mtDNA sequences.

### RAD-seq Datasets Building

Genomic analyses were performed on several datasets, which varied in number and geographical distribution of the individuals, and list of loci to include in the analyses (*white list*). We decided to replicate several of the analyses on multiple datasets in order to gain confidence on the inferred pattern. Details on each of the dataset are presented in Supplementary Appendix S3: Table S4.

### Phylogenomic Analysis

We employed a maximum-likelihood approach (RAXML v8.2.11; Stamatakis 2014) to infer evolutionary relationships among the five *Eliurus* taxa. We performed concatenated analyses on datasets composed of fixed sites, among which invariant sites were removed. The concatenated analysis assumes that all the sites included in the multialignment share the same species tree topology (Felsenstein 1981). We set the unpartitioned General Time-Reversible Categorization (GTRCAT) molecular model with joint branch length optimization, the ‘-f a’ option to conduct 1000 rapid Bootstrap analysis, and the ‘autoMRE_IGN’ option to carry out bootstrap convergence test, which determines whether sufficient bootstrap replicates have been run to get stable support values in the inferred phylogenetic tree. We also carried out a partitioned analysis on the *N15.pop5.p4.min15* dataset, including all sites (variant + fixed), setting each *catalog* locus as a separate partition. We applied an ascertainment bias correction for binary SNP data in the RAXML analysis based on Paul O. Lewis formula (Lewis 2001). The partitioned analysis estimates the model parameters of the most likely evolutionary model for each partition while fixing the global set of branch lengths (genealogy). As for the concatenated analysis, we set GTRCAT molecular model for each partition. In this analysis we also used the argument ‘-M’, to estimate a separate set of branch lengths for every partition. For each maximum-likelihood analysis, the best likelihood tree with bootstrap supported value was visualized using *ggtree* and *treeio* R packages (Yu et al. 2017; Yu 2020; Wang et al. 2020).

### Genomic Clustering analyses

Clustering methods were used to identify genetic groups within the *Eliurus* dataset. We used the widely-used maximum likelihood model-based approach implemented in ADMIXTURE v1.3 (Alexander et al. 2009). This program uses the same likelihood model as STRUCTURE software (Pritchard et al. 2000), but instead of sampling the posterior probability of a model given the data, it maximises the likelihood of the model. In the model, it is assumed that individuals are sampled from an admixed population with contributions from K *a priori* clusters. Then, individuals are assigned to K-clusters by maximizing the likelihood of the model. For each dataset, we ran the program for K values ranging from 1 to 8, and repeated the analysis for 30 independent runs. ADMIXTURE analysis was carried out using the default optimization method and the default termination criterion, which stops the analysis when the log-likelihood increases by less than 10^−4^ between iterations. K values were compared using the cross-validation method by setting --cv = 20 and by comparing likelihoods across runs. We used CLUMPAK (Kopelman et al. 2015) to permute the 30 independent runs so that clusters align across runs. We selected the best K number of clusters considering the lower average cross-validation, but we also explored admixture models for K - 1 and K + 1. The final assignment of individuals to the best K-value was obtained by taking the mode containing most runs (*Major.Cluster*).

In addition, we used a model-free approach, the Principal Components Analysis (PCA), through the function *glPca* implemented in the R package *adegenet 2.0.1* (Jombart and Ahmed 2011). PCA constructs low-dimensional projections of the data that explain the gross variation in marker genotypes, and thus it can be useful to assess whether most of the variation in the genomic dataset can be explained by species divergence. ADMIXTURE and PCA analyses were performed on the *N124.pop5.p4.r80.min100* dataset.

Admixture and PCA analyses were also performed on the GLs-based ANGSD dataset using the NGSadmix and PCAngsd programs (Skotte et al. 2013). NgsAdmix was run for 20 iterations per K value (1 to 12) and a stringent tolerance of 10^−6^ for convergence. We compared K values using the ΔK procedure (Evanno et al. 2005).

### Species Validation

We used three approaches to test the hypothesis that the newly identified taxon (*Eliurus* sp. nova) was a new species: i) guided-tree species delimitation analyses; ii) estimation of the gdi-index; iii) comparison of isolation-by-distance patterns within *versus* between taxa; and iv) comparison of GC-content across taxa. Species delimitation analysis with guided tree was performed in BPP (Yang and Rannala 2010; Flouri et al. 2018, 2020) using loci full-length FASTA file of the three individuals per species dataset (*N15.pop5.p4.min15* dataset). We used six combinations of priors for divergence time (τ) and population size (θ). The θ and τ priors consisted of an inverse gamma distribution with α = 2 (θ) or α = 3 (τ) and β equal to either 0.002 or 0.02 or 0.2. Six combinations of priors are tested to ensure that model selection would not be influenced by prior information. Four MCMC runs were carried out for each combination of parameters, and convergence was checked by comparing the outputs from the four runs. Each run was conducted with a burnin of 5000 steps, sample frequency of 100 steps (i.e., thinning), and collected 200,000 samples. Each analysis was repeated using both rjMCMC algorithms in BPP, algorithm0 and algorithm1. Prior combinations, two distinct MCMC algorithms, and four runs per analysis were conducted to ensure convergence of the results. The tree model with higher posterior probabilities across all runs was selected as the best tree topology.

The genealogical divergence index (*gdi*; Jackson et al. 2017; Leaché et al. 2019) was calculated using the posterior distribution of θ and τ from the analysis A00 carried out in BPP (Yang 2015). Analysis A00 estimates divergence times and population sizes using a guided specie tree, under the Multi-Species Coalescent (MSC) model. As for the species delimitation test, we ran the analysis for each of the six combinations of priors for θ and τ, and four runs for each combination. The details of the MCMC run were the same as for the species delimitation test. The posterior distribution of the *gdi*-index was calculated by computing the *gdi*-index for each of the 200,000 MCMC samples, across runs, using *equation 7* from Leaché et al. (2019): *gdi* = 1 − *e*^−2τ/*θ*^. The *gdi*-index analysis was carried out only to test the species hypothesis for *Eliurus sp. nova*, therefore we used the estimated divergence time (τ) between *Eliurus sp. nova* and *E. myoxinus* (its sister-species), and alternatively used as population size (θ) either the one estimated for *Eliurus sp. nova* or for *E. myoxinus*. This index provides a continuous measure, that uses information on the estimated divergence times and population sizes for a fixed species tree under the MSC, to assess where, a given taxon, lies on the path of speciation. Although the *gdi* index does not really solve the dichotomy of population structure *versus* species delimitation in MSC models, it allows to go beyond the discrete model selection (e.g., one-species *versus* two-species).

Complementary to the two abovementioned species delimitation methods, which ignore the importance of geographically-limited dispersal in determining genetic differences between populations, we compared the individual-based genomic and geographic (Euclidean) distances within (e.g., *E. myoxinus* - *E. myoxinus*) and among taxa (e.g., *E. myoxinus* - *Eliurus* sp. nova). We performed this analysis on the STACKS generated datasets (including only variant sites) by estimating pairwise genetic distances using the *stamppNeisD* function in R (R Core Team 2022) which computes the Nei’s genetic distance (Nei 1972).

It has been shown that across animal genomes, GC-content correlates with various aspects of genome architecture such as genome size, gene density, gene expression, replication timing, and recombination (Vinogradov 1998; Bernardi and Bernardi 1986; Bernardi 2007; Figuet et al. 2015). Few case studies have shown that GC-content may also be informative on the evolutionary relationships of species (e.g., Du et al. 2010; Huttener et al. 2019). Accordingly, we asked whether differences in GC content could be observed among the five *Eliurus* species in our dataset, as a proxy for differences in genomic architecture among *Eliurus* species. For measuring the percentage of GC content within each *Eliurus* species, we used a multialignment that included *catalog* loci present in at least 100 out of the 124 individuals in the dataset. All sites (variant and invariant) were extracted and the %GC was computed using the *GC.content* function implemented in the *ape* R package (Paradis and Schliep 2019).

### Morphological Analyses

Four morphological variables (head length, tail length, body length and weight) were consistently recorded for the *Eliurus* individuals. They were used here to assess whether morphological differences could be detected between the five *Eliurus* taxa. For increasing the power of the morphological analyses, we used a dataset composed of 325 *Eliurus* individuals, which included the 124 individuals used in phylogenetic and species validation analyses, as well as 200 individuals that were excluded from the above analysis due to computational cost. We carried out Discriminant Analysis of Principal Component (DAPC) to identify which morphological variable contributes the most to the phenotypic structure observed across the five *Eliurus* taxa. DAPC analysis was performed using the *dapc* function in *adegenet* R package (Jombart 2008; Jombart and Ahmed 2011), fixing species identity as prior grouping. Furthermore, we compared the distribution of each morphological variable across *Eliurus* species and assess significant differences through multiple comparison Tukey’s test, as implemented in the *TukeyHSD* R function (R Core Team 2022). In addition, we assessed whether morphology would correlate with altitude and 19 bioclimatic variables. For each individual, altitude and bioclimatic data were extracted from the WorldClim database (www.worldclim.org; Fick and Hijmans 2017) at a spatial resolution of 30 seconds of a degree using the *getData* function in the *raster* R package (Hijmans 2022). Correlations between each morphological and bioclimatic variable were measured using Phylogenetic Generalized Least Squares (PGLS) analysis (Harvey and Pagel 1991). Briefly, PGLS analysis correlates two variables while accounting for the phylogenetic relatedness among individuals, given that closely related taxa tend to have more similar traits than random. The PGLS analysis was carried out in R using the *gls* function of *nlme* R package (Pinheiro and Bates 2000; Pinheiro et al. 2022). Covariance structure was based on the model of Brownian evolution through the *corBrownian* function. The phylogenetic tree used to account for phylogenetic relatedness was generated from the full *Eliurus* dataset (N = 347), including also few individuals for which morphological data were not available or incomplete (N = 23) (Supplementary Appendix S3: Fig. S3). All individuals from the dataset were assumed to belong to the same ‘population’ (p = 1) and loci were included in the final dataset only if 80% of the individuals were genotyped at a given locus. Moreover, in order to reduce linkage disequilibrium among genotyped SNPs, we considered only one random SNP per RAD locus. The last filtering step was carried out in *plink* v1.9 (Purcell et al. 2007) for removing loci that were missing for > 10% of individuals and removed individuals that contained >40% of missing data. The phylogenetic tree was constructed in R using neighbour-joining method as implemented in *njs* R function (Paradis and Schliep 2019).

### Eliurus Species Distribution and Co-Occurrence

We estimated species range areas for the five *Eliurus* taxa by computing the minimum convex hull among all sampling sites across Madagascar for which species identity has been genetically determined in the present and previous studies. The minimum convex hull was computed using the *Minimum Bounding Geometry* algorithm in QGIS 3.22 (QGIS.org 2022). We also quantified *Eliurus* species co-occurrence across northern Madagascar, defined as the percentage of sampling sites for each species where at least one other genetically identified *Eliurus* species was recorded.

## Results

### Species Identification from mtDNA cytb Reference Phylogeny

Using Maximum-likelihood and Bayesian based phylogenetic analyses, the 20 individuals genotyped at the mtDNA *cytb* grouped with four previously described *Eliurus* species: *E. carletoni* (N=2), *E. ellermani* (N=3), *E. minor* (N=2) and *E. myoxinus* (N=6) (Fig. 2a; Supplementary Appendix S3: Fig. S3, S4). We observed major topological differences between ML and Bayesian trees at the deeper nodes. The evolutionary relationships reconstructed using Bayesian-based phylogenetic method (MRBAYES) were consistent with previous *Eliurus* phylogenies (e.g., Jansa et al. 2019). The phylogenetic analyses revealed also a novel mtDNA *Eliurus* lineage (N=7), sister-clade of *E. myoxinus*, which hereafter we will refer to as *Eliurus* sp. nova.

**Figure 2.**
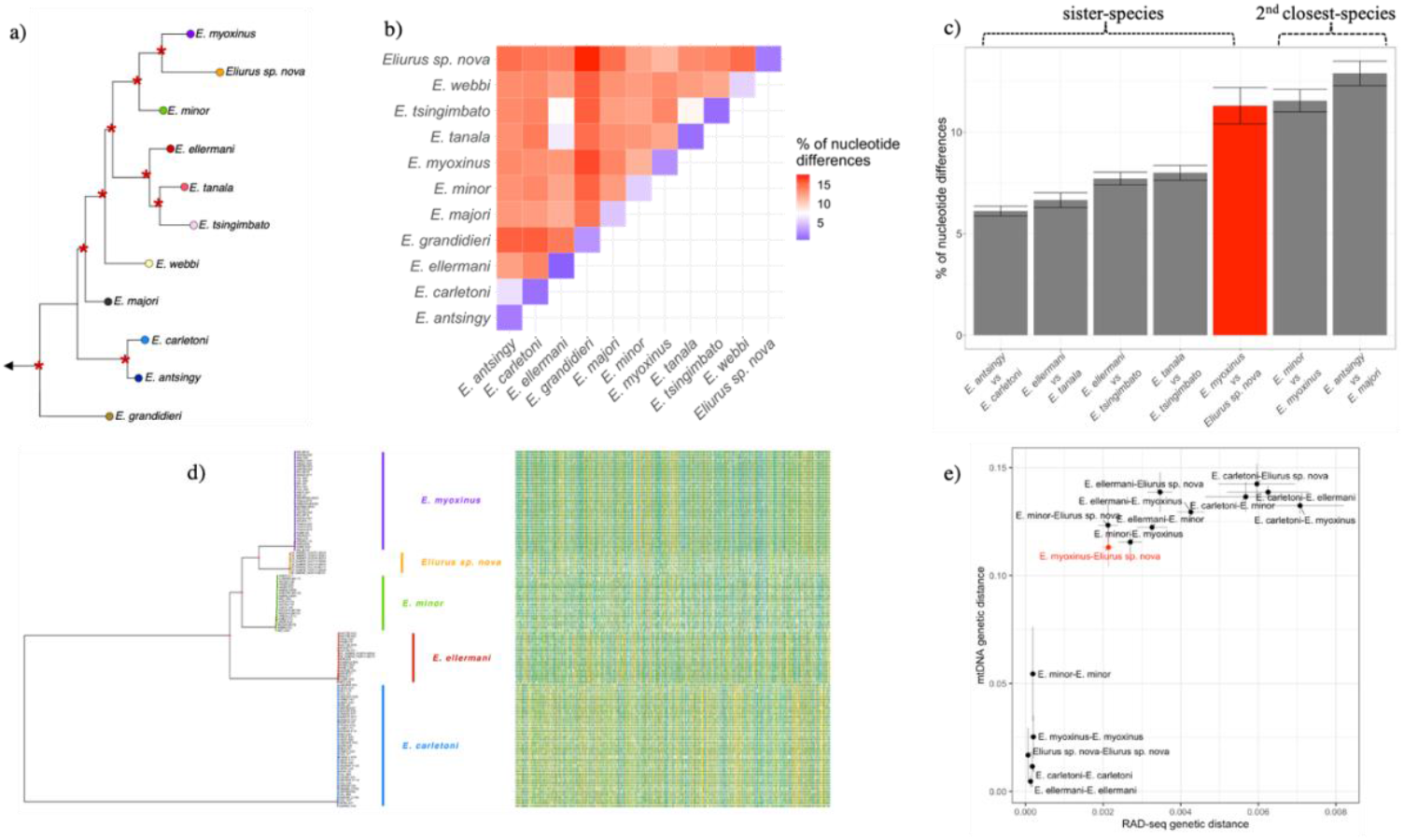
Inference of the phylogenetic relationships within the *Eliurus* genus. a) Bayesian-based phylogeny obtained from MRBAYES. The outgroup EU349782.1 corresponds to mtDNA *cytb* of *Rattus norvegicus*. b) Matrix of the mtDNA *cytb* p-distances between *Eliurus* species pairs. c) mtDNA *cytb* p-distances between closely-related *Eliurus* species pairs. The red bar represents the *Eliurus* sp. nova – *E. myoxinus* mtDNA *cytb* p-distance, which is larger than the other four comparisons of closely-related *Eliurus* species pairs. d) Maximum-likelihood based phylogeny of *Eliurus* samples based on nuclear RAD-seq data (*N124.pop5.p4.min100*; 4.351 SNPs). On the right side is showed the multialignment from which the phylogeny has been inferred. e) Correlation of pairwise *Eliurus* sp. genetic differences between mtDNA *cytb* and nuclear RAD-seq data. The results suggest a strong correlation between mtDNA *cytb* and nuclear RAD-seq phylogenies (Spearman rho: 0.896; p-value<2.2e-16), and that *Eliurus* sp. nova could be considered a newly discovered *Eliurus* species, holding among the highest level of *Eliurus* sister-species genetic divergence.

The level of average genetic differences (*p* − *distance*) across all *Eliurus* species pairs ranged between 6.12% (*E. antsingy* – *E. carletoni*) and 17.6% (*E. grandidieri* – *Eliurus* sp. nova), with a median of 13.4% (Fig. 2b). Among all recognised closely-related *Eliurus* species pairwise comparisons, the *p* − *distance* value of *E. antsingy* – *E. carletoni* was the lowest (6.12%), whereas the value of *E. antsingy* – *E. majori* was the highest (12.9%) (Fig. 2c). The two sister taxa *Eliurus* sp. nova and *E. myoxinus* showed a high level of average genetic differences among sister-species (*p* − *distance* = 11.3%), similar to *E. minor* - *E. myoxinus* (*p* − *distance* = 11.6%).

### RAD-data Processing: de novo Assembly of Nuclear Loci

After the three filtering steps described in the *Material and Methods* section (*process_radtags, clon_filter, kmer_filter*) and data processing (*ustacks, cstacks, sstacks, tsv2bam, gstacks*) of each RAD-seq dataset, we obtained in total about 500 million short-reads (half paired-end and half single-end), with an average of ∼4 M reads per individual. The minimum number of forward reads recovered per individual was 201 K and the maximum was 5.2 M reads (Table 1). The optimization analysis of *de novo* assembly parameters showed that the highest amount of polymorphism for *r* = 80% was at *m* = 2, *M* = 5, *n* = 8 (Supplementary Appendix S3: Fig. S5). The cleaned *catalog* of *Eliurus spp*. was composed of 327,870 loci with an average length of ∼780 bp. Across the whole RAD-seq dataset, the weighted mean individual coverage was between ∼5X and ∼40X (Supplementary Appendix S3: Fig. S6). Across the eleven STACKS datasets, we recovered between ∼9,000 and ∼89,662 variant sites, whereas the ANGSD dataset included ∼392,000 sites.

### Phylogenomic Analysis

The maximum-likelihood based phylogeny inferred using the full RAD-seq dataset (*N124.pop5.p4.min100*) was consistent with the mtDNA *cytb* phylogeny (Spearman rho: 0.896, p-val < 2.2e-16; Fig. 2a, d, e), although not all *Eliurus* species were present in the RAD-seq dataset. The maximum-likelihood based phylogenies were topologically identical across all RAD-seq datasets, regardless of the sampling scheme, sample size, or list of included loci (Supplementary Appendix S3: Fig. S7). For the dataset including the lowest sample size (*N15.pop5.p4.min15*), we also performed partitioned analysis on RAXML, in which, for each partition (equivalent to RAD-locus), the method estimates the model parameters of the most likely evolutionary model, while fixing the global set of branch lengths (Supplementary Appendix S3: Fig. S8a). In addition, we also inferred the maximum likelihood tree for each partition independently (Supplementary Appendix S3: Fig. S8b). A qualitative comparison of the tree recovered using the partitioned analysis on RAXML and the ML trees inferred for each partition showed consistencies between concatenated, partitioned and gene phylogenies (Supplementary Appendix S3: Fig. S7 and Fig. S8).

### Genomic Structure

Clustering analyses conducted with ADMIXTURE and NGSadmix identified the best number of genetic clusters at K = 6 (Supplementary Appendix S3: Fig. S9). The coefficients of individual ancestry estimated from the two approaches showed that at K = 4, *E. myoxinus, E. carletoni, E. ellermani* and *E. minor* were primarily separated, whereas *Eliurus* sp. nova showed a mixed ancestry between *E. minor* and *E. myoxinus*, with all individuals exhibiting the exact same pattern (Fig. 3). These results may be due to the fact that, with the exception of *Eliurus* sp. nova and *E. myoxinus*, our genomic data do not include sister-species for the other taxa. At K = 5, ADMIXTURE results identified population structure within *E. carletoni* as an additional genetic cluster, and at K= 6, *Eliurus* sp. nova individuals could be genetically separated from the two closely-related species *E. minor* and *E. myoxinus*. NGSadmix results showed clear genetic differences between *Eliurus* sp. nova and the other *Eliurus* species at K = 5, while revealing population structure within *E. carletoni* at K= 6 (Supplementary Appendix S3: Fig. S10). The PCA analyses performed on genotype calls (STACKS dataset) and likelihood (ANGSD dataset) showed five distinct clusters of individuals (Fig. 3 and Supplementary Appendix S3: Fig. S10), corresponding to the genetic clades identified in the phylogenetic analyses. The first principal component (PC1) explained ∼ 50% of the variance, contributing mostly to the delimitation of *E. carletoni* against the other *Eliurus* species. The PC2 explained ∼ 15% of the variance, clearly separating the remaining four *Eliurus* taxa. A similar pattern of separation could be seen with PC3 (∼ 5%), while at PC4 (∼ 1%) major differences were observed between *Eliurus sp. nova* and the other four *Eliurus* species.

**Figure 3.**
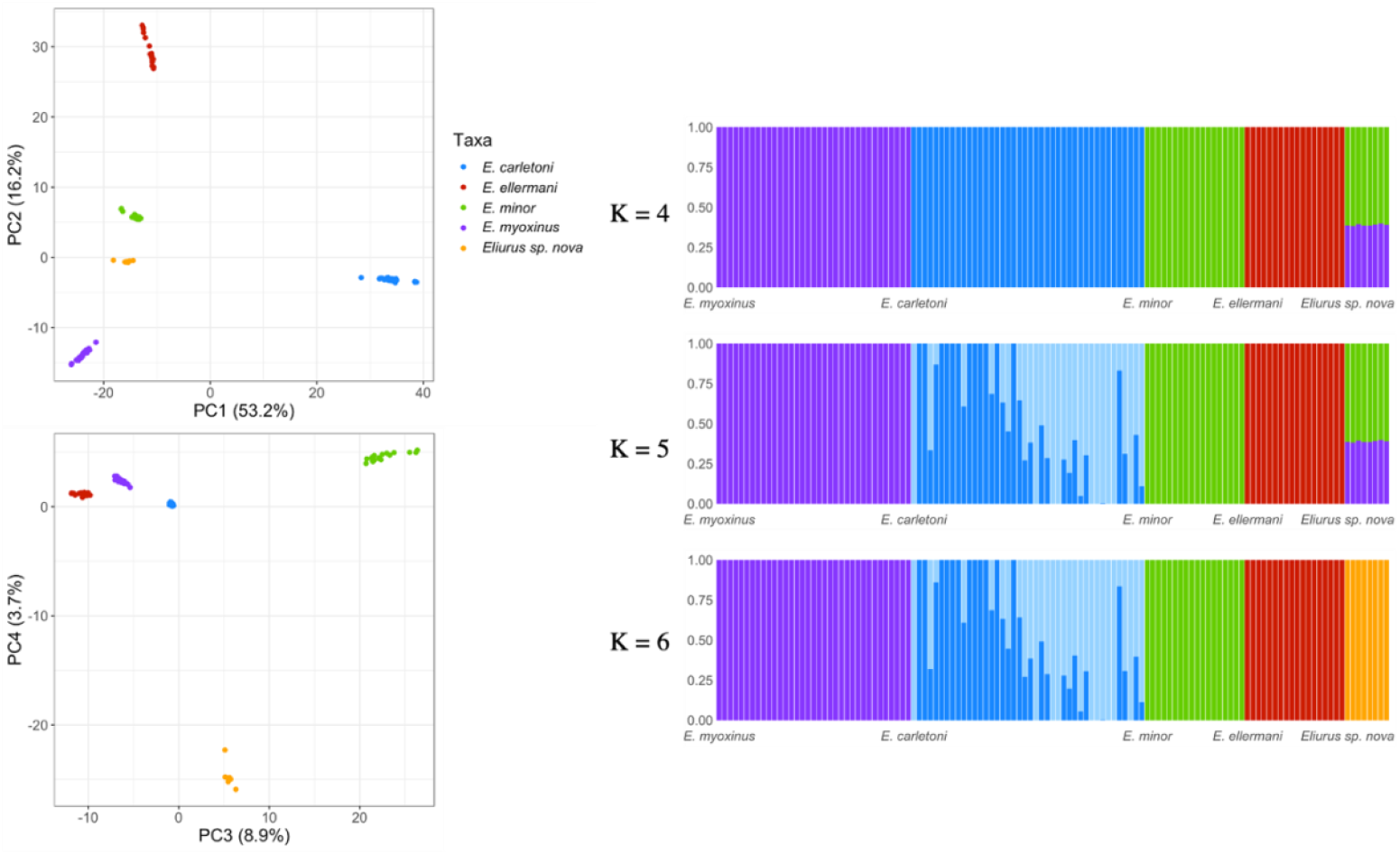
Genetic structure of sampled *Eliurus* individuals in northern Madagascar. PCA and ADMIXTURE analyses separate individuals according to the hypothesized species identification (*E. carletoni*; *E. myoxinus*; *E. ellermani*; *E. minor*) while identifying the hypothetically new *Eliurus* species (*Eliurus sp. nova*) as being genetically distinct from the others, and, in particular, from the sister-species *E. myoxinus*. Both analyses have been performed using the *N124.pop5.p4.min100* dataset which includes 9,040 variant sites.

### Eliurus Species Delimitation

The phylogenomic and genomic structure analyses above appear to support the hypothesis of a new distinct *Eliurus* taxon: *Eliurus* sp. nova. The species delimitation analyses carried out with BPP, using all sites (variant and invariant) of the *N15.pop5.p4.min15* dataset, for *E. myoxinus, E. minor* and *Eliurus* sp. nova, strongly supported the delimitation of the three taxa as distinct species. This topology was always the best species tree with a posterior probability of 1.0, across all runs, algorithms and priors (Fig. 4a). The *N15.pop5.p4.min15* dataset was also used to estimate divergence time and population size parameters using BPP. The estimated values were then used to calculate the *gdi*-index, which was essentially equal to 1 across all runs and combination of priors (Fig. 4b). For both BPP analyses, MCMC convergence was reached and posterior parameters distributions were concordant across runs, priors’ combinations and MCMC algorithms (Supplementary Appendix S3: Fig. S11, S12).

**Figure 4.**
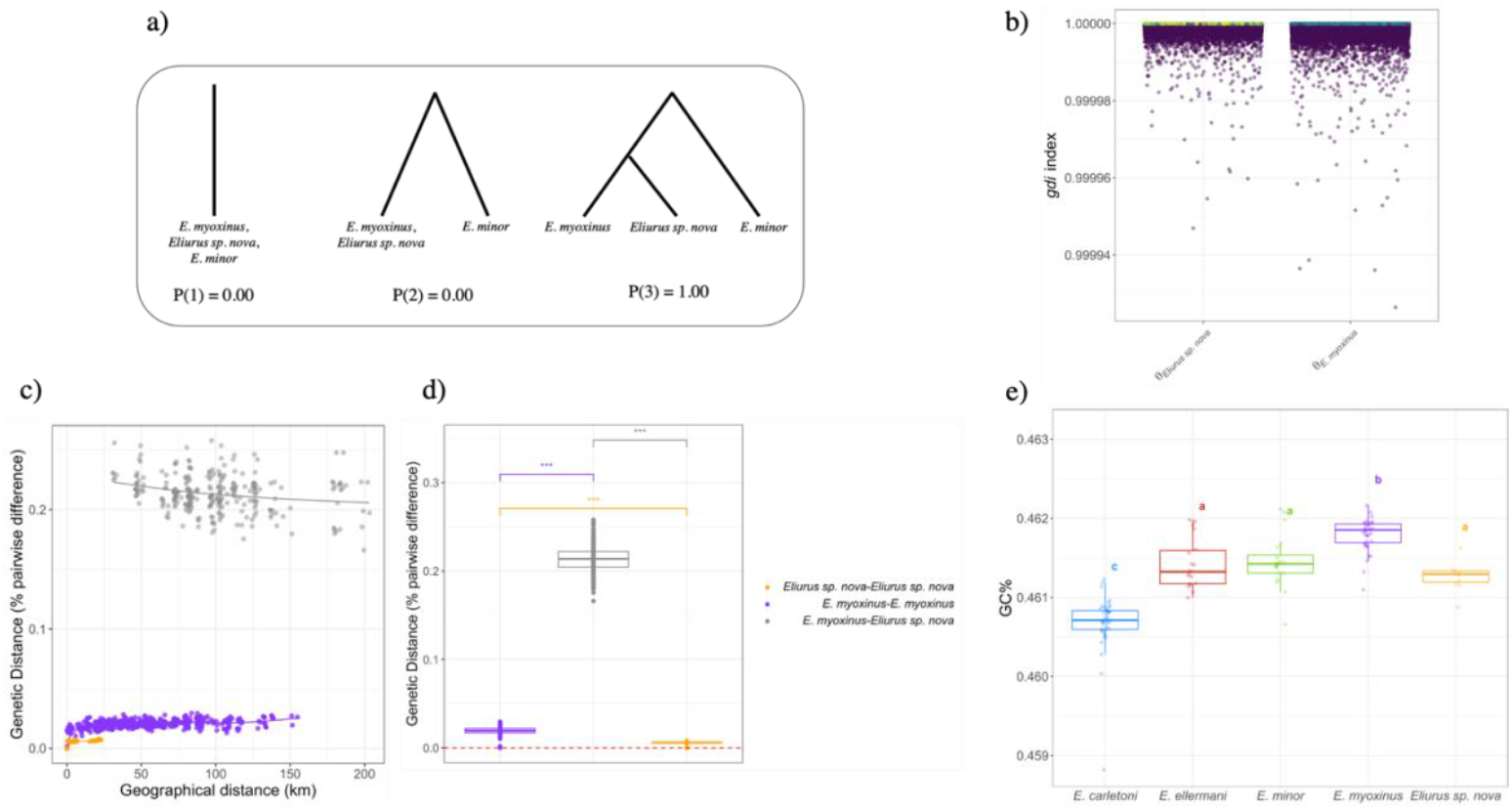
Species delimitation analyses for *E. myoxinus, Eliurus* sp. nova and *E. minor*. a) Alternative species delimitation models generated from the hypothetical fixed guide tree ((*E. myoxinus, Eliurus sp. nova*), *E. minor*) in BPP. P(1), P(2), P(3) refers to the posterior probabilities for the best species tree model across four runs for each of the two tested rjMCMC algorithms (finietune0 and finetune1) and six combinations of population size and divergence time priors; b) *gdi*-index calculated using the estimated parameters (*θ*_*E. myoxinus*_; *θ*_*Eliurus sp.nova*_, τ_*E.myoxinus*−*Eliurus sp.nova*_) under the multispecies-coalescent model implemented in BPP. Dots represent gdi-values estimated for each run, combination of parameters and MCMC steps; c) Isolation by distance within taxon (*E. myoxinus, Eliurus sp. nova*) and between taxa; d) Comparison of genetic distance within and between taxa; e) percentage of GC content (%GC) across *Eliurus* species, estimated from all sites (variant and invariant) extracted from the *N124.pop5.p4.min100* dataset (6,162,356 bp).

The *ad hoc* test that compares isolation-by-distance (IBD) patterns for individuals belonging to the same taxon (*E. myoxinus* - *E. myoxinus* or *Eliurus* sp. nova - *Eliurus* sp. nova) with IBD patterns for individuals assigned to different taxa (*E. myoxinus* - *Eliurus* sp. nova) (Fig. 4c) strongly suggest that *Eliurus* sp. nova does not fit in a model of intraspecific spatial genetic structure within *E. myoxinus*. Indeed, we found positive and significant IBD for *E. myoxinus* (Mantel’R= 0.484; p-value=0.001) and *Eliurus* sp. nova (Mantel’R= 0.7386; p-value=0.001), whereas negative significant IBD was detected for *Eliurus* sp. nova-*E. myoxinus* (Spearman *rho*= -0.187; p-value= 0.0017). The results showed also that, for similar geographical distances, *Eliurus* sp. nova - *E. myoxinus* genetic differences were significantly higher than within taxa genetic differences (Fig. 4d). Similar patterns could be observed for two well-recognized *Eliurus* species, *E. ellermani* and *E. minor*, for which the average proportion of genetic differences is slightly higher than *Eliurus* sp. nova-*E. myoxinus* species pairs (*p* − *distance*_*cytb*_ = 12.3%; Supplementary Appendix S3: Fig. S13).

The percentage of GC content across the *Eliurus* species included in our dataset ranged between 46.06% (*E. carletoni*) and 46.18% (*E. myoxinus*) (Fig. 4e). We did not observe significant differences among *E. ellermani, E. minor* and *Eliurus* sp. nova, however we found significantly different (but partially overlapping) GC content distributions between *Eliurus* sp. nova and its sister-taxa *E. myoxinus* (Fig. 4e).

### Morphological Differences among Eliurus Species

The DAPC analysis showed that among the four morphological variables available in our dataset, “head length” had the most weight for discriminating the five *Eliurus* species (Fig. 5a; Supplementary Appendix S3: Fig. S14b). In particular, the Tukey multiple comparison of means revealed significant differences in head length between two groups of species, namely *E. carletoni* - *E. ellermani versus E. minor* - *E. myoxinus* - *Eliurus* sp. nova (p-value<0.01). The test was also significant for the comparison between *E. myoxinus* and *E. minor* - *Eliurus* sp. nova (p-value<0.01) (Fig. 5b). Similar differences were detected also for “weight” and “body length” (Supplementary Appendix S3: Fig. S14c, e), as expected since they are likely positively correlated. Morphological analyses thus detected significant differences between the two sister taxa *E. myoxinus* and *Eliurus* sp. nova, the latter being consistently smaller than *E. myoxinus* across the three significant morphological variables. Comparisons with previously published morphological data confirmed the interspecies morphological differences observed in our dataset (Supplementary Appendix S3: Fig. S15). Moreover, the PGLS analysis on each morphological variable showed that “body length”, “tail length” and “weight” were negatively correlated with altitude (slope = -0.01, -0.03, - 0.03; p-value = 0.05, <0.01, <0.01, respectively), whereas no correlation could be detected for “head length” (Fig. 5; Supplementary Appendix S2: Table S5). The PGLS analysis carried out on 19 bioclimatic variables exhibited results consistent with those obtained from altitude, overall showing a decline in body size with decreasing temperature. However, we found significant relationships only between “tail length” or “weight” and part of the bioclimatic variables, respectively 10 and 16 out of 19 bioclimatic variables (Supplementary Appendix S3: Fig. S16-S19; Supplementary Appendix S2: Table S5).

**Figure 5.**
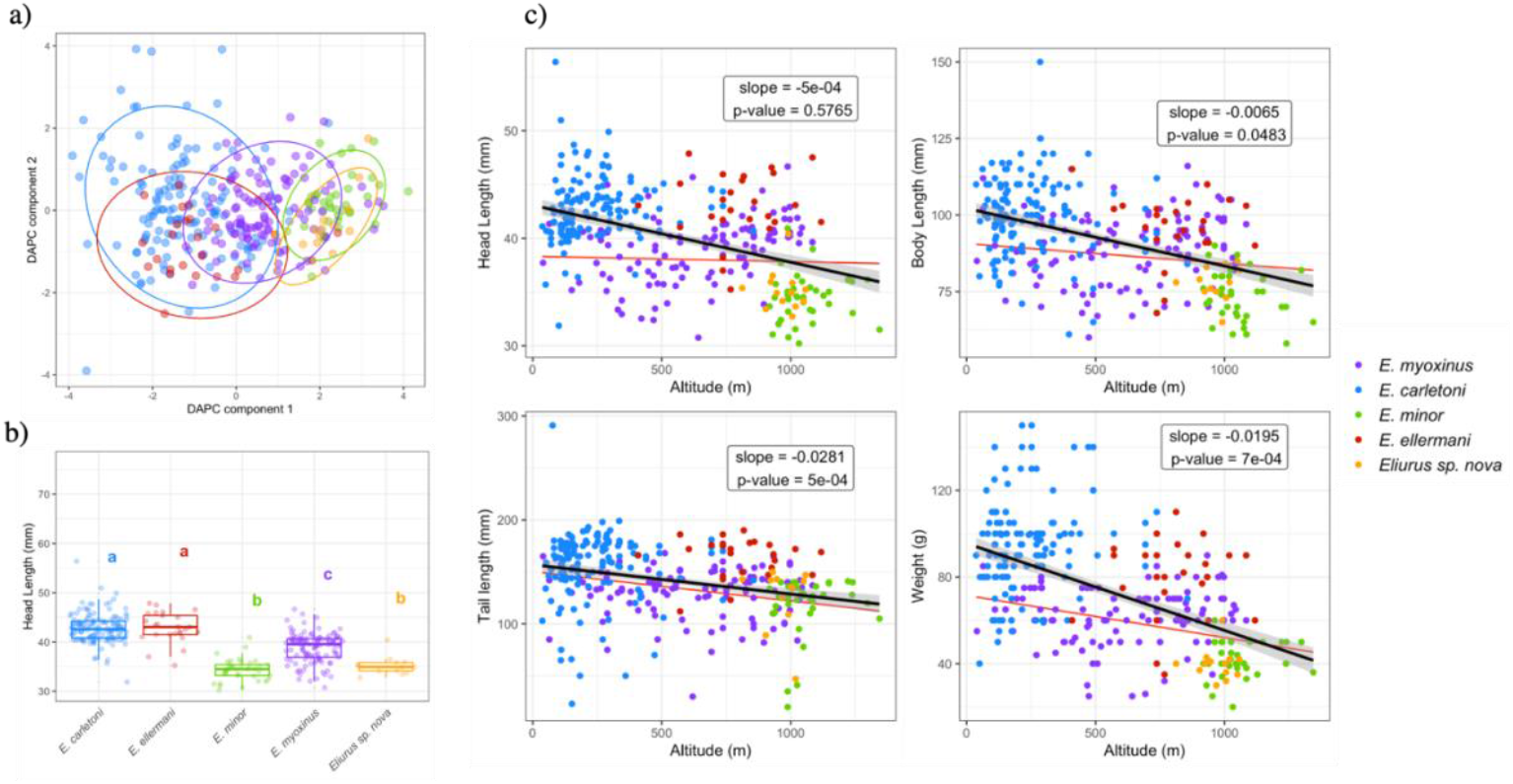
Morphological analysis and correlation with bioclimatic variables. a) DAPC plot for the first two principal components based on four morphological variables across genetically determined *Eliurus* individuals (N = 325) sampled across northern Madagascar; b) Multiple comparison Tukey test on “head length”, the variable that contribute the most on interspecies morphological structure; c) PGLS analysis between each morphological variable and altitude. Black line indicates linear regression without accounting for phylogenetic relatedness. Red line indicates linear regression based on PGLS analysis.

### Eliurus Species Distribution in Northern Madagascar

In the present study, we also report new records of *Eliurus* species occurrence across northern Madagascar (Supplementary Appendix S3: Fig. S20), leading to an update of the species distribution in area and elevation. Among the four previously known *Eliurus* species, *E. myoxinus* and *E. minor* presented the widest distribution in Madagascar, ∼378,000 km^2^ and ∼261,423 km^2^, respectively, followed by *E. ellermani* and *E. carletoni* with, ∼45,500 km^2^ and ∼6,980 km^2^. *Eliurus* sp. nova was found in Montagne d’Ambre (∼ 249 km^2^), and never identified in the other forests we visited. The new records presented in this study led to an update in species range area (minimum convex hull) of 69% for *E. carletoni*, 8.26% for *E. myoxinus*, 7.38% for *E. ellermani* and 2.51% for *E. minor*. The altitudinal range varied across *Eliurus* species (Supplementary Appendix S3: Fig. S21): between 37 m and 1,002 m for *E. carletoni*; between 40 m and 1,284 m for *E. myoxinus*; between 289 m and 1,558 m for *E. ellermani*; between 289 m and 2,016 m for *E. minor*; between 812 m and 1,057 m for *Eliurus* sp. nova. We found significant differences in elevational range among all *Eliurus* sp. pairwise comparisons, except for *E. minor* - *Eliurus* sp. nova (p-value=0.14) and *E. ellermani* - *Eliurus* sp. nova (p-value=0.78).

## Discussion

In the present study, we analysed mitochondrial and nuclear genomic data of 124 Malagasy endemic tuft-tailed rats (*Eliurus* genus) collected in northern Madagascar. We showed that the species delineation previously defined using mitochondrial data were supported by nuclear genomic data. Our analyses have also identified an undescribed *Eliurus* clade (*Eliurus* sp. nova), recovered from individuals collected in the humid forests of Montagne d’Ambre, in the far north of Madagascar. We hypothesized that the newly identified *Eliurus* genetic clade would correspond to an evolutionary unit distinct from the other known *Eliurus* species. We tested our hypothesis by performing a series of phylogenetic inferences using ML and MSC based approaches, and by evaluating pattern of genomic and morphological diversity. We assessed the support for the *Eliurus sp. nova* species hypothesis by using five approaches: i) the species delimitation tests based on MSC model, ii) *gdi*-index, iii) the *ad-hoc* test of isolation-by-distance within *versus* between sister-taxa; iv) GC-content analysis; and v) morphological analyses. Our results support the delimitation of the newly discovered *Eliurus* clade as a distinct species. In addition, we showed that, in northern Madagascar, the dry-habitat associated *E. myoxinus* can actually be mostly found in humid habitats. We discuss below the implications of such findings in the context of *Eliurus* species evolution and diversification. Lastly, we confirm that *Eliurus* species co-occurrence is common in northern Madagascar, and update the distribution of previously known species.

### Eliurus Mitochondrial Genetic Diversity

Among the 124 individuals sampled in the dry, humid and transition forests across a region of ∼12,000 km^2^, we genetically assigned the sampled individuals to four previously known *Eliurus* species (*E. carletoni, E. myoxinus, E. minor, E. ellermani*). In addition, we identified a new *Eliurus* taxon (*Eliurus* sp. nova). The phylogenetic relationships inferred from mtDNA *cytb* sequences were in agreement with previous phylogenies (Jansa et al. 1999, 2019), and placed the new taxon, *Eliurus* sp. nova, as sister-taxon of *E. myoxinus* (Fig. 2a). The mtDNA *cytb* genetic divergence between *Eliurus* species pairs suggested strong level of genetic differentiation within the *Eliurus* genus (6.12% - 17.6%; Fig. 2b). Previous analyses at the mtDNA *cytb* of six out of thirteen *Eliurus* species showed similarly high values of pairwise species genetic differences (4.7% - 13.2%; Jansa et al. 2019). As a comparison, the average species genetic differences measured at the same mitochondrial locus in a small primate genus endemic of Madagascar (*Microcebus* spp.) ranged between 0.9% and 15.3% (e.g., Sgarlata et al. 2019). The *Eliurus* sp. nova - *E. myoxinus* sister-taxa showed the highest level of genetic differences compared to the other *Eliurus* sister-species comparisons, and the second highest when also the second most closely-related species were considered (Fig. 2c), strongly supporting, on the basis of the mtDNA *cytb*, the species hypothesis for *Eliurus* sp. nova. While no new *Eliurus* species has been described in Montagne d’Ambre, Tysdal and Jansa (2014) reported in a poster the presence of a new *Eliurus* taxon sampled in the northwest of Madagascar. They also found it to be closely related to *E. myoxinus* and *E. minor* clade, but not as sister-taxon of *E. myoxinus*. Future work should confirm whether the *Eliurus* sp. nova identified in the present study and the new taxon in Tysdal and Jansa (2014) belongs to the same lineage.

### Nuclear Genomic Diversity of Eliurus Species in Northern Madagascar

The comparison of individuals genotyped at both mtDNA and nuclear DNA have confirmed the presence of five *Eliurus* taxa in our dataset (Supplementary Appendix S3: Fig. S3). The results based on nuclear genome-wide data confirmed the tree topology recovered using MrBayes from mtDNA *cytb*, and were in agreement with the topology obtained in previous studies (e.g., Jansa et al. 2019). Genome-wide PCA and clustering analysis have further confirmed the genomic differentiation across the five *Eliurus* taxa (Fig. 3 and Supplementary Appendix S3: Fig. S10). We did not find clear signature of admixed co-ancestry among *Eliurus* species, which may lead to exclude events of recent admixture or hybridization among them, at this stage (Fig. 3). The congruence obtained among our different analyses can be considered a good indicator of true evolutionary signal in the data (Hillis 1995; Rubin et al. 2012), and therefore we propose that, according to the phylogenetic species concept (Rosen 1979; Baum and Shaw 1995), *Eliurus* sp. nova should correspond to an undescribed *Eliurus* species, possibly the same as the one mentioned by Tysdal and Jansa (2014).

### Eliurus Species Delimitation

Classical concatenation models of gene sequences, which assumes topologically congruent genealogies across the genome, oversimplify the genome-wide variability of the biological processes (e.g., incomplete lineage sorting, recombination) (Rannala and Yang 2003; Edwards et al. 2016). The multispecies-coalescent (MSC) model has been proposed as an alternative model able to capture such variability and estimate gene trees, together with species trees, by taking into account the diversity of genealogical histories in the genome (Rannala and Yang 2020, 2003). The MSC model has been shown to perform better for species identification than genetic distance approaches that rely on empirical distance thresholds (Yang and Rannala 2017). Moreover, recent studies have shown that MSC-based approaches outperform concatenation models for species tree inference (Jiang et al. 2020). The additional analyses performed under the MSC model implemented in BPP strongly supported the separation of *E. minor, E. myoxinus* and *Eliurus* sp. nova as separate species, with ((*E. myoxinus, Eliurus sp. nova*), *E. minor*) as the species tree with highest support (*P*_(3)_ = 1; Fig. 4), across all runs and combinations of priors.

Two recent studies (Jackson et al. 2017; Leaché et al. 2019) have shown that even in cases where two populations (or putative species) exchange genes at a relatively high rate (*Nm* = 10), such that they should be considered nearly one population, MSC-based species delimitation approaches would select a two-species model over a one-species model. In other words, MSC-based approaches may favour species over-splitting even in the case of ongoing gene-flow between populations, and thus mis-interpret population structure as evidence for species delimitation. To overcome this limitation in MSC-based species delimitation models, Jackson et al. (2017) have proposed the *gdi* index, which calculates the degree of non-monophyly in a set of gene-trees. Using the parameters estimated under MSC with ((*E. myoxinus, Eliurus sp. nova*), *E. minor*) as species tree, we found that the *gdi*-index was consistently higher than 0.999 across runs and combinations of priors. Based on Pinho and Hey (2010) meta-analysis, Jackson et al. (2017) proposed that *gdi* < 0.2, should be interpreted as evidence for one single species, a *gdi* > 0.7 as evidence for two-species model, and 0.2 ≤ *gdi* ≤ 0.7 as ambiguous delimitation. According to this rule of thumb, our *gdi* results strongly supported the delimitation of *Eliurus* sp. nova a separate species.

We also performed an *ad hoc* test, aimed at evaluating if the *Eliurus* sp. nova clade could be the result of insufficient geographical and taxon sampling, thus be part of the spatial genetic structure within *E. myoxinus*. The reasoning underlying such test is that species are often characterized by geographically limited dispersal (e.g., Sexton et al. 2014), such that geographically close individuals tend to be genetically more related than individuals far apart, called isolation-by-distance (IBD) (Wright 1943; Malécot 1948). The signal of spatial genetic structure due to limited dispersal can create cline in genetic diversity that if not taken into account, especially when taxa are insufficiently sampled, can lead to misinterpret intraspecies population structure as evidence for species delimitation (Barley et al. 2018; Hausdorf and Hennig 2020; Mason et al. 2020). We found that individuals mtDNA identified as *E. myoxinus* exhibit signatures of IBD at the nuclear genome, suggesting that *E. myoxinus* belong to those species characterised by geographically-limited dispersal. Therefore, we compared IBD patterns within *E. myoxinus* and *Eliurus* sp. nova, with IBD computed only among *E. myoxinus* - *Eliurus* sp. nova individual pairs. If *Eliurus* sp. nova individuals are not a population of *E. myoxinus*, we would expect i) no IBD for *E. myoxinus* - *Eliurus* sp. nova comparisons; and ii) significantly higher genetic distances between taxa than within taxa, keeping in mind that genetic differentiation is expected to reach plateau (stationary value of differentiation) for large geographical distances (Rousset 1997, 2000). In this regard, our results showed that *Eliurus* sp. nova is likely a distinct species and not the result of insufficient taxon sampling and spatial genetic structure within the sister-taxa *E. myoxinus* (Fig. 4c, d).

Finally, we found significant differences in GC content for all *Eliurus* species pairwise comparisons, except *E. ellermani*-*E. minor*-*Eliurus* sp. nova (Fig. 4e). Most importantly, we showed significant differences between *Eliurus* sp. nova and its sister-species *E. myoxinus*. The small differences recovered among *Eliurus* species (GC = 46.06% - 46.18%) are consistent with GC content pattern across placental mammals (Romiguier et al. 2010). For instance, Romiguier et al. (2010) showed similar small differences in GC content between *Homo sapiens* (GC3 = 46.1%) and *Pan troglodytes* (GC3 = 46.09%), or between *Rattus norvegicus* (GC3 = 51.46%) and *Mus musculus* (GC3 = 51.24%). Quantifying GC content across species can be informative on differences in genomic architecture, since it has been shown that GC content correlates with several genomic features, such as gene density, methylation rate, recombination rate, and expression levels (Eyre-Walker and Hurst 2001; Kudla et al. 2006). Thus, differences in %GC between the two sister-taxa *Eliurus* sp. Nova and *E. myoxinus* suggest that the genomic architecture of these two taxa may be significantly different, further supporting the initial hypothesis that *Eliurus* sp. nova should be considered a new *Eliurus* species. Moreover, this observation points out that important “islands of differentiation” may be present in the genome of these two taxa and should be considered in future work.

### Morphological Diversity of the Eliurus genus in Northern Madagascar

Morphological analyses provided further evidence for distinguishing *Eliurus* sp. nova from *E. myoxinus* (Fig. 5b and Supplementary Appendix S3: Fig. S14c-e). Overall, morphological data suggested that *Eliurus* sp. nova and *E. minor* have similar size, and are smaller than *E. carletoni, E. myoxinus* and *E. ellermani*. Such patterns were consistent with those obtained from previously published morphological data for *E. carletoni, E. ellermani, E. myoxinus* and *E. minor* (Supplementary Appendix S3: Fig. S15). Interestingly, we found that across *Eliurus* species, “body length”, “tail length” and “weight” negatively correlated with altitude (Fig. 5c), suggesting that *Eliurus* body size tend to decrease at higher altitude. Since higher elevation correlates with colder climate (e.g., Supplementary Appendix S3: Fig. S16), the pattern observed in the *Eliurus* genus rejects Bergman’s rule, which states that species occurring in colder areas have larger body size than species occurring in warmer areas (Bergmann 1847). It has been proposed that such clines in body size minimize heat loss by decreasing the surface-to-volume ratio for species in colder areas (Mayr 1956). Although support for Bergman’s rule has been found across mammal and bird taxa (e.g., Blackburn et al. 1999; Ashton et al. 2000; Meiri and Dayan 2003; Blackburn and Hawkins 2004; Romano et al. 2020), several other studies have shown that this is not always the case (e.g., Adams and Church 2008; Rodríguez et al. 2008; Feldman and Meiri 2014; Maestri et al. 2016; Hendges et al. 2021). For example, Rodríguez et al. (2008) and Maestri et al. (2016) have reported positive correlation between body size and ambient temperature across more than 250 mammal species sampled in the Neotropics (most of which were sigmodontine rodents). The conclusion drawn by Rodríguez et al. (2008) is that the Bergmann’s heat-conservation mechanism, which underlies the increase in body size with latitude (i.e., towards colder areas), should apply only to regions with cold climate (e.g., Nearctic). Instead, in regions with warm climate (e.g., Neotropics), topography would play a major role in shaping body size across mammals (i.e., mountainous *versus* lowland). One possible explanation, discussed by Rodríguez et al. (2008), is that climate and habitat quality are similar in mountains and lowlands of cold regions, while showing a stronger gradient in warm region. Because lower temperature correlates with lower habitat quality in mountains of the warm regions, a possible explanation for the positive correlation between body size and temperature is that the mountainous habitats of the Neotropics are not good enough for sustaining many species of large body size (Alhajeri and Steppan 2016). The trend found by Rodríguez et al. (2008) and Maestri et al. (2016) in the Neotropics recall the negative correlation that we found between body size and altitude in *Eliurus* individuals sampled across the diverse topography of northern Madagascar. Accordingly, we propose that the decline in body size observed in *Eliurus* individuals sampled at higher altitude may be explained by the putative lower quality of the mountainous habitats compared to lowland habitats.

### Eliurus Species Distribution and Biogeography in Northern Madagascar

The importance of exhaustive geographic sampling has been a central debate in taxonomy, but it has only recently received more attention (Meyer and Paulay 2005; Wiemers and Fiedler 2007; Bergsten et al. 2012). Regarding the *Eliurus* genus, for instance, the study of Jansa et al. (2019) showed that increasing the geographic coverage of samples belonging to the *Eliurus tanala* group (*E. ellermani, E. tanala* and *E. tsingimbato*) provided a better representation of population variation in species morphology. In particular, Jansa et al. (2019) revealed that a trait thought to be unique within *E. ellermani* was actually shared with its close relative *E. tanala*.

In the light of the crucial role that geographical sampling plays, we combined the new data reported in the present work with those from previous studies, and carried out an updated analysis of the geographic distribution and elevational range of the *Eliurus* species found in northern Madagascar. We found that among the five *Eliurus* taxa herein studied, *E. myoxinus* and *E. minor* are the species with the widest distribution across Madagascar, respectively ∼378,000 km^2^ and ∼261,423 km^2^ (Supplementary Appendix S3: Fig. S20). Prior to this study, *E. myoxinus* was mostly known from the dry deciduous forests and xerophytic scrub bush of western Madagascar, with few reported occurrences in the humid forests of northern-east Madagascar (Puy and Moat 1996; Soarimalala and Goodman 2003; Shi et al. 2013). The present study reveals frequent occurrences of *E. myoxinus* also in northern Madagascar, in both dry and lowland humid forests at altitudes ranging from 40 – 1284 m (Supplementary Appendix S3: Fig. S21). In particular, we found that among the sites where *E. myoxinus* individuals have been sampled for the present study, 77% (14/18) are characterised by humid habitat and 22% (4/18) by dry habitat (Fig. 1 and Table 1). In northern Madagascar, the first reporting of *E. myoxinus* individuals in humid habitats dates back to the work of Goodman and Soarimalala (2002) and Soarimalala and Goodman (2003), documenting the presence of this species in the Special Reserve of Manongarivo (in the northwest), in the humid forests of Loky-Manambato protected area (in the north) and National Park Marojejy (in the northeast). Morphological comparisons of *E. myoxinus* individuals collected from the south to the north of Madagascar showed a cline in body size, from the largest size in the south to the smallest in the north (Carleton and Goodman 2007). Such decline in body size in the direction of the most humid northern populations, coincides with the increase in morphological similarity of *E. myoxinus* with *E. minor*, known to be associated with humid habitats of north and east Madagascar. The increasing similarity with the humid-associated *E. minor*, and the occurrence of *E. myoxinus* in both dry and mesic habitats, may point out to a mechanism of local adaptation or phenotypic plasticity. However, the fact that the other two closely-related *Eliurus* lineages (*E. minor* and *Eliurus* sp. nova) are humid-associated species may suggest that the ancestor of *E. myoxinus* was adapted to humid habitat and that *E. myoxinus* expanded southward. A similar hypothesis has been proposed in the unpublished work of Tysdal and Jansa (2014), where the authors compared *Eliurus* phylogeny with the habitat type to which each *Eliurus* species is associated. They found that i) seven out of eleven *Eliurus* species occurs in humid habitats, and that ii) dry-associated *Eliurus* species are not necessarily phylogenetically related. They proposed that most speciation in the *Eliurus* genus occurred in humid and evergreen forests, and that at least three independent events of adaptation to dry habitats occurred within the genus, specifically for: i) *E. myoxinus*; ii) *E. tsingimbato*; and iii)

### E. carletoni + E. antsingy

With this study, we also report, for the first-time, evidence for the presence of *E. carletoni* in humid habitats of northern Madagascar, corresponding to 24% (5/21) of *E. carletoni* sampling sites (e.g., ANDRA, ANDR, SAL, BEZ and ANALV; Fig. 1). Similar findings have been reported by Goodman et al. (2009), which sampled *E. carletoni* individuals in mesic vegetations within the deep canyons of the Ankarana Massif. The presence of *E. carletoni* in both dry and humid habitats could be explained by i) ecological tolerance, allowing this species to occupy areas of moister habitats; or ii) the presence of dry microclimatic conditions that favour *E. carletoni* persistence in mostly humid localities. *E. myoxinus* and *E. carletoni* represent two interesting case studies for possibly studying adaptation to dry and humid habitats. At this stage, this scenario is largely speculative and it will require genetic and detailed phenotypic data across the whole species range to test such hypothesis and assess to which extent dry-adaptation is due to genetic adaptation or phenotypic plasticity.

In the dry southern Madagascar, previous studies have reported the presence of *E. myoxinus* in the small moisty areas of the canyons of Isalo and Analavelona Massif (Carleton et al. 2001). These two localities have been suggested to be Quaternary relics of wetter climatic conditions across this region (Burney 1993; Goodman and Rakotozafy 1997; Raxworthy and Nussbaum 1997; Carleton et al. 2001), further supported by evidence of ancient connectivity between the present-day humid eastern and dry western Madagascar (Goodman and Rakotozafy 1997; Jansa et al. 1999; Muldoon et al. 2009; Goodman et al. 2013b; Yoder et al. 2016). In fact, Jansa et al. (1999) found in the canyons of Isalo (in the southwest), together with *E. myoxinus*, also an *Eliurus* form related to the eastern *E. majori*. Muldoon et al. (2009) found in the Ankilitelo Cave (in the south-west) subfossils of shrew tenrecs, carnivorans and rodents which have modern distributions in areas more humid than the present-day dry southwestern Madagascar. Yoder et al. (2016) found stronger genetic proximity between two mouse lemurs (*Microcebus*) occurring in different ecoregions, the humid forests of the east (*M. rufus*) and the dry deciduous forests of the west (*M. berthae*), than mouse lemurs occurring in the same ecoregion, suggesting that forest-dwelling species such as *Microcebus* could disperse across the Central Highlands in the ancient past.

Assuming that *E. myoxinus* was originally restricted to humid habitats, we propose that also the humid forest fragments where *E. myoxinus* has been found in northern Madagascar represents relics of an ancient wetter period. In fact, these humid forest fragments coincide with sites that have been suggested to represent humid refugia (Sgarlata et al. 2019) when dry conditions increased in northern Madagascar during the Holocene (Jungers et al. 1995; Simons et al. 1995; Godfrey et al. 1999). For instance, Sgarlata et al. (2019) have reported the presence in these fragmented humid forests of another small mammal, the Arnhold’s mouse lemur (*Microcebus arnholdi*), which until 2019 was known only from Montagne d’Ambre (in the far north) and from one site in north-west Madagascar (Weisrock et al. 2010). The discovery of new *M. arnholdi* populations in fragmented humid forests surrounded by dry forests has led the authors, supported by fossil evidences (Jungers et al. 1995; Simons et al. 1995; Godfrey et al. 1999), to hypothesise that in the recent past northern Madagascar was most likely covered by humid habitat, and that following the aridification of the Holocene only few mountain forests could maintain humid conditions suitable to the persistence of humid-adapted species. According to this scenario, only some populations of *E. myoxinus* and *M. arnholdi* have persisted during the dry conditions of the Holocene. Similar scenarios have been proposed to explain the demographic history, and distribution of two *Eliurus* species in northern Madagascar (*Eliurus carletoni* and *Eliurus tanala*; Rakotoarisoa et al. 2013a, b), sifaka (*Propithecus tattersalli*; Quéméré et al. 2012; Salmona et al. 2017) and the leaf chameleons (e.g., *Brookesia ebenaui* and *Brookesia minima*; Raxworthy and Nussbaum 1995). Future population genomic analyses may shed light on the time scale of forest cover change in northern Madagascar and on the environmental changes that have shaped its present-day biodiversity.

### Eliurus Sympatry: more common than rare

In the present study, we also document frequent cases of *Eliurus* species co-occurrence (sympatry and sintopy) in northern Madagascar (Table 1; Fig. 1). In particular, we found a co-occurrence frequency of 43%, 90%, 38% and 56%, respectively for *E. carletoni, E. ellermani, E. minor* and *E. myoxinus* with at least one other *Eliurus* species occurring in the study region. *Eliurus* sp. nova is currently only known from Montagne d’Ambre, and thus in co-occurrence with *E. ellermani. Eliurus* records collected in previous studies have shown that sympatry between *Eliurus* species is common (Carleton 1994; Jansa et al. 2019; Supplementary Appendix S2: Table S6). As pointed out by Goodman et al. (2013a), however, *Eliurus* sympatry rarely involves closely-related species. The genetic analyses we performed, such as ADMIXTURE or PCA, did not reveal signature of ongoing admixture between sympatric *Eliurus* species, suggesting that even if in co-occurrence, these species are most likely reproductively isolated. Not many studies have looked at how several *Eliurus* species can co-exist in the same sites and what are the mechanisms behind reproductive isolation. A PhD thesis by Katrin Marquart (Marquart 2014), has however shown that niche repartition may be one of the mechanisms that make *Eliurus* species coexistence possible. In fact, Marquart (2014) observed that, among the four *Eliurus* taxa found in Maromizaha, north-eastern Madagascar (*E. minor, E. tanala, E. webbi, E. grandidieri*), some species could be preferentially captured at ground level and medium high vegetation (e.g., *E. tanala*), while others exclusively in high vegetation (*E. webbi*). The author also identified different behaviours across *Eliurus* taxa, some considered great climbers (*E. minor*) whereas others seen using ground level or swimming along riverside (*E. tanala*). Some of these differences in behaviour and microhabitat-use were qualitatively correlated with differences in the morphology of the posterior hind (*E. webbi* versus *E. tanala-E. grandidieri-E. minor*), which led the author to conclude that certain morphological structures of *Eliurus* feet give evidence of a specific way of life and that foot pad morphology in particular, mirrors special adaptations to a species’ habitat.

### Conclusions and Future Directions

In the present work, we have assessed the species genomic diversity of the *Eliurus* genus in northern Madagascar, representing the first multi-species genomic study of the genus. We document the discovery of a new species, *Eliurus* sp. nova, in the mountain humid forest of Montagne d’Ambre, supporting the idea that this mountain forest contains extraordinary levels of biodiversity across several taxa and represent an ‘island’ of humid habitat surrounded by drier habitat (Raxworthy and Nussbaum 1994; Louis et al. 2008; Rasolonjatovo et al. 2020). The concordance between gene trees and species tree, together with the differences in %GC detected among the studied species, emphasize major differences in genomic architecture among *Eliurus* taxa. Future work may focus on detecting ‘islands of genomic differentiation’ among closely-related *Eliurus* species, in order to identify genomic regions involved in reproductive isolation or in the adaptation to dry *versus* humid habitats. We also showed that, in northern Madagascar, *E. myoxinus* occurs mostly in humid habitats, in contrast to what was previously known for this species. We also document the presence of the dry-associated *E. carletoni* in humid forests of northern Madagascar.

These findings might have important implications for our understanding of *Eliurus* evolution and diversification. In fact, the *E. myoxinus* populations identified in the humid forests of northern Madagascar may contain genetic information that could contribute to our understanding of the genomic basis of adaptation to dry and humid habitats. Ultimately, we confirmed that *Eliurus* species often co-exist in the same forest fragments, suggesting niche separation of *Eliurus* species within the same environment. Additional work is required to uncover the eco-evolutionary mechanisms that make such co-existence possible.

## Data Availability

Raw sequencing data is available at NCBI BioProject XXXX.

## Supplementary Appendix

Data available from the Dryad Digital Repository: http://dx.doi.org/10.5061/dryad.[NNNN]

## Acknowledgments

We thank all the Malagasy MSc students, field assistants, volunteers, local guides and cooks that contributed significantly to this work, by sharing their knowledge and participating at the collection of the data. We thank in particular Jacquis Randriamaroson for his valuable help and support over several fieldwork seasons. We also thank Filipa Borges for her help with PCR amplification of mtDNA data. We thank the Direction Générale du Ministère de l’Environnement et des Forêts de Madagascar, Direction Régionale de l’Environnement, Ecologie et des Forêts (DREEF DIANA and SAVA), Madagascar’s Ad Hoc Committee for Fauna and Flora and Organizational Committee for Environmental Research (CAFF/CORE), Conservation International, WWF, Missouri Botanical Garden and the Fanamby NGO. This study benefited from the continuous support of the Department of Animal Biology and Ecology, University of Mahajanga, the Department of Animal Biology, University of Antananarivo, Department of Natural and Envornmental Sciences at the University of Antsiranana.

## References

Adams D.C., Church J.O. 2008. Amphibians Do Not Follow Bergmann’s Rule. Evolution. 62:413–420.

Alexander D.H., Novembre J., Lange K. 2009. Fast model-based estimation of ancestry in unrelated individuals. Genome Res. 19:1655–1664.

Alhajeri B.H., Steppan S.J. 2016. Association between climate and body size in rodents: A phylogenetic test of Bergmann’s rule. Mamm. Biol. 81:219–225.

Andrews S. 2010. A quality control tool for high throughput sequence data. http://www.bioinformatics.babraham.ac.uk/projects/fastqc/.

Andrews K.R., Good J.M., Miller M.R., Luikart G., Hohenlohe P.A. 2016. Harnessing the power of RADseq for ecological and evolutionary genomics. Nat. Rev. Genet. 17:81–92.

Ashton K.G., Tracy M.C., Queiroz A. de. 2000. Is Bergmann’s Rule Valid for Mammals? Am Nat. 156:390–415.

Baird N.A., Etter P.D., Atwood T.S., Currey M.C., Shiver A.L., Lewis Z.A., Selker E.U., Cresko W.A., Johnson E.A. 2008. Rapid SNP Discovery and Genetic Mapping Using Sequenced RAD Markers. PLOS ONE 3:e3376.

Barley A.J., Brown J.M., Thomson R.C. 2018. Impact of Model Violations on the Inference of Species Boundaries Under the Multispecies Coalescent. Syst. Biol. 67:269–284.

Baum D.A., Shaw K.L. 1995. Genealogical perspectives on the species problem. In: Hoch P.C., Stephenson A.G. editors. Experimental and Molecular Approaches to Plant Biosystematics. St. Louis: Missouri Botanical Garden. p. 53:289–303.

Bergmann C. 1847. Über die Verhältnisse der Wärmeökonomie der Thiere zu ihrer Größe.

Bergsten J., Bilton D.T., Fujisawa T., Elliott M., Monaghan M.T., Balke M., Hendrich L., Geijer J., Herrmann J., Foster G.N., Ribera I., Nilsson A.N., Barraclough T.G., Vogler A.P. 2012. The effect of geographical scale of sampling on DNA barcoding. Syst. Biol. 61:851–869.

Bernardi G., Bernardi G. 1986. Compositional constraints and genome evolution. J. Mol. Evol. 24:1–11.

Bernardi G. 2007. The neoselectionist theory of genome evolution. Proc. Natl. Acad. Sci. U. S. A. 104:8385–8390.

Bésairie H. 1964. Carte géologique à 1:1.000.000 de Madagascar. Tananarive: Serv. Geol. Madagascar.

Bickford D., Lohman D.J., Sodhi N.S., Ng P.K.L., Meier R., Winker K., Ingram K.K., Da I. 2007. Cryptic species as a window on diversity and conservation. Trends Ecol. Evol. 22:148–155.

Blackburn T.M., Gaston K.J., Loder N. 1999. Geographic gradients in body size: a clarification of Bergmann’s rule. Diversity and Distributions. 5:165–174.

Blackburn T.M., Hawkins B.A. 2004. Bergmann’s rule and the mammal fauna of northern North America. Ecography. 27:715–724.

Burney D.A. 1993. Late Holocene Environmental Changes in Arid Southwestern Madagascar. Quat. Res. 40:98–106.

Carleton M., Goodman S. 1998. New taxa of nesomyine rodents (Muroidea: Muridae) from Madagascar’s northern Highlands, with taxonomic comments on previously described forms. In: Goodman S.M. editor. A Floral and Faunal Inventory of the Réserve Spéciale d’Anjanaharibe-Sud, Madagascar With Reference to Elevational Variation. Zoology, New Series No. 90 Fieldiana. p. 163–200.

Carleton M.D. 2003. Eliurus, tufted-tailed rats. In: Goodman S.M., Benstead J.P. editors. The Natural History of Madagascar. Chicago: University of Chicago Press. p. 1373–1380.

Carleton M.D. 1994. Systematic studies of Madagascar’s endemic rodents (Muroidea: Nesomyinae): revision of the genus Eliurus. Am. Mus. Novit. 3087:1–55.

Carleton M.D., Goodman S.M. 2007. A new species of the Eliurus majori complex (Rodentia, Muroidea, Nesomyidae) from south-central Madagascar, with remarks on emergent species groupings in the genus Eliurus. Am. Mus. Novit. 3547:1–21.

Carleton M.D., Goodman S.M., Rakotondravony D. 2001. A new species of tufted-tailed rat, genus Eliurus (Muridae: Nesomyinae), from western Madagascar, with notes on the distribution of E. myoxinus. Proc. Biol. Soc. Wash. 114:972–987

Chenuil A., Cahill A.E., Délémontey N., Du Salliant du Luc E., Fanton H. 2019. Problems and Questions Posed by Cryptic Species. A Framework to Guide Future Studies. In: Casetta E., Marques da Silva J., Vecchi D. editors. From Assessing to Conserving Biodiversity: Conceptual and Practical Challenges, History, Philosophy and Theory of the Life Sciences. Springer International Publishing, Cham. p. 77–106.

Du H., Hu H., Meng Y., Zheng W., Ling F., Wang J., Zhang X., Nie Q., Wang X. 2010. The correlation coefficient of GC content of the genome-wide genes is positively correlated with animal evolutionary relationships. FEBS Lett. 584:3990–3994.

Du Puy D.J., Moat J. 1996. A refined classification of the primary vegetation of Madagascar based on the underlying geology: Using GIS to map its distribution and to assess its conservation status. In: Lourenco, W.R., editor. Biogeography of Madagascar. Paris: ORSTOM. p. 205–218.

Edwards S.V., Xi Z., Janke A., Faircloth B.C., McCormack J.E., Glenn T.C., Zhong B., Wu S., Lemmon E.M., Lemmon A.R., Leaché A.D., Liu L., Davis C.C. 2016. Implementing and testing the multispecies coalescent model: A valuable paradigm for phylogenomics. Mol. Phylogenet. Evol. 94:447–462.

Etter P.D., Bassham S., Hohenlohe P.A., Johnson E.A., Cresko W.A., 2011. SNP discovery and genotyping for evolutionary genetics using RAD sequencing. In: Orgogozo V., Rockman, M.V., editors. Molecular Methods for Evolutionary Genetics. New York: Humana Press. p. 157–178.

Evanno G., Regnaut S., Goudet J. 2005. Detecting the number of clusters of individuals using the software structure: a simulation study. Mol. Ecol. 14:2611–2620.

Ewels P., Magnusson M., Lundin S., Käller M. 2016. MultiQC: summarize analysis results for multiple tools and samples in a single report. Bioinformatics 32:3047–3048.

Eyre-Walker A., Hurst L.D. 2001. The evolution of isochores. Nat. Rev. Genet. 2:549–555.

Feldman A., Meiri S. 2014. Australian Snakes Do Not Follow Bergmann’s Rule. Evol Biol. 41:327–335.

Felsenstein, J. 1981. Evolutionary trees from DNA sequences: A maximum likelihood approach. J. Mol. Evol. 17:368–376.

Fick S.E., Hijmans R.J. 2017. WorldClim 2: new 1-km spatial resolution climate surfaces for global land areas. Int. J. Clim. 37:4302–4315.

Figuet E., Ballenghien M., Romiguier J., Galtier N. 2015. Biased Gene Conversion and GC-Content Evolution in the Coding Sequences of Reptiles and Vertebrates. Genome Biol. Evol. 7:240–250.

Flouri T., Jiao X., Rannala B., Yang Z. 2020. A Bayesian Implementation of the Multispecies Coalescent Model with Introgression for Phylogenomic Analysis. Mol. Biol. Evol. 37:1211–1223.

Flouri T., Jiao X., Rannala B., Yang Z. 2018. Species Tree Inference with BPP Using Genomic Sequences and the Multispecies Coalescent. Mol. Biol. Evol. 35:2585–2593.

Glez-Peña D., Gómez-Blanco D., Reboiro-Jato M., Fdez-Riverola F., Posada D. 2010. ALTER: program-oriented conversion of DNA and protein alignments. Nucleic Acids Res. 38:W14–W18.

Godfrey L.R., Jungers W.L., Simons E.L., Chatrath P.S., Rakotosamimanana B. 1999. Past and Present Distributions of Lemurs in Madagascar. In: Rakotosamimanana B., Rasamimanana H., Ganzhorn J.U., Goodman S.M. editors. New Directions in Lemur Studies. Boston: Springer US. p. 19–53.

Goodman S.M., Raheriarisena M., Jansa S.A. 2009. A new species of Eliurus Milne Edwards, 1885 (Rodentia: Nesomyinae) from the Réserve Spéciale d’Ankarana, northern Madagascar. Bonn. Zool. Beitr. 56:133–149.

Goodman S.M., Raherilalao M.J., Muldoon K. 2013b. Bird fossils from Ankilitelo Cave: inference about Holocene environmental changes in southwestern Madagascar. Zootaxa 3750:534–548.

Goodman S.M., Rakotozafy L.M.A. 1997. Subfossil birds from coastal sites in western and southwestern Madagascar: a paleoenvironmental reconstruction. In: Goodman S.M., Patterson B.D. editors. Natural Change and Human Impact in Madagascar. Washington, D.C.: Smithsonian Institution Press. p. 257–279.

Goodman S.M., Raselimanana A.P., Wilmé L. 2007. Inventaires de la faune et de la flore du couloir forestier d’Anjozorobe-Angavo. Recherches pour le Developpement, Série Sciences Biologique no. 24. Centre d’Information et de Documentation Scientifique et Technique, Antananarivo, Madagascar

Goodman S.M., Rasolonandrasana B.P.N. 1999. Inventaire biologique de la Réserve spéciale du Pic d’Ivohibe et du couloir forestier qui la relie au Parc national d’Andringitra. Rech. dév., Sér. Sci. biol. 15:1–180

Goodman S.M., Soarimalala V. 2002. Les petits mammifères de la Réserve Spéciale de Manongarivo, Madagascar. Boissiera 59: 384–401.

Goodman S. M., Soarimalala V., Raheriarisena M., Rakotondravony D. 2013a. Small mammals or tenrecs (Tenrecidae) and rodents (Nesomyinae). In: Goodman S.M., Raherilalao, M.J. editors. Atlas of Selected Land Vertebrates of Madagascar. Antananarivo: Association Vahatra. p. 211–269.

Harvey P.H., Pagel M.D. 1991. The Comparative Method in Evolutionary Biology. Oxford ; New York: Oxford University Press.

Hausdorf B., Hennig C. 2020. Species delimitation and geography. Mol. Ecol. Res. 20:950–960

Hendges C.D., Patterson B.D., Cáceres N.C. 2021. Big in the tropics: Ecogeographical clines in peccary size reveal the converse of Bergmann’s rule. J. Biogeogr. 48:1228–1239.

Hijmans R. 2022. raster: Geographic Data Analysis and Modeling. R package version 3.5-21.

Hillis D.M., 1995. Approaches for Assessing Phylogenetic Accuracy. Syst. Biol. 44:3–16.

Huttener R., Thorrez L., in’t Veld T., Granvik M., Snoeck L., Van Lommel L., Schuit F. 2019. GC content of vertebrate exome landscapes reveal areas of accelerated protein evolution. BMC Evol. Biol. 19:144.

Jackson N.D., Carstens B.C., Morales A.E., O’Meara B.C. 2017. Species Delimitation with Gene Flow. Syst. Biol. 66:799–812.

Jansa S.A., Carleton M.D. 2003. Systematics and phylogenetics of Madagascar’s native rodents. In: Goodman S.M., Benstead J.P. editors. The Natural History of Madagascar. Chicago: iversity of Chicago Press. p. 1257–1265

Jansa S.A., Carleton M.D., Soarimalala V., Rakotomalala Z., Goodman S.M. 2019. A review of the Eliurus tanala complex (Rodentia, Muroidea, Nesomyidae), with description of a new species from dry forests of western Madagascar. Bull. Am. Mus. Nat. Hist. 430:1–69.

Jansa S.A., Goodman S.M., Tucker P.K. 1999. Molecular Phylogeny and Biogeography of the Native Rodents of Madagascar (Muridae: Nesomyinae): A Test of the Single-Origin Hypothesis. Cladistics 15:253–270.

Jansa, S.A., Weksler, M., 2004. Phylogeny of muroid rodents: relationships within and among major lineages as determined by IRBP gene sequences. Mol. Phylogenet. Evol. 31:256–276.

Jiang X., Edwards S.V., Liu L. 2020. The Multispecies Coalescent Model Outperforms Concatenation Across Diverse Phylogenomic Data Sets. Syst. Biol. 69:795–812.

Jombart T. 2008. adegenet: a R package for the multivariate analysis of genetic markers. Bioinformatics 24:1403–1405.

Jombart T., Ahmed I. 2011. adegenet 1.3-1: new tools for the analysis of genome-wide SNP data. Bioinformatics 27:3070–3071.

Jungers W.L., Godfrey L.R., Simons E.L., Chatrath P.S. 1995. Subfossil Indri indri from the Ankarana Massif of northern Madagascar. Am. J. Phys. Anthropol. 97:357–366.

Kearse M., Moir R., Wilson A., Stones-Havas S., Cheung M., Sturrock S., Buxton S., Cooper A., Markowitz S., Duran C., Thierer T., Ashton B., Meintjes P., Drummond A. 2012. Geneious Basic: an integrated and extendable desktop software platform for the organization and analysis of sequence data. Bioinformatics 28:1647–1649.

Kopelman N.M., Mayzel J., Jakobsson M., Rosenberg N.A., Mayrose I. 2015. Clumpak: a program for identifying clustering modes and packaging population structure inferences across K. Mol. Ecol. Resour. 15:1179–1191.

Kudla G., Lipinski L., Caffin F., Helwak A., Zylicz M. 2006. High Guanine and Cytosine Content Increases mRNA Levels in Mammalian Cells. PLOS Biology. 4:e180.

Leaché A.D., Zhu T., Rannala B., Yang Z. 2019. The Spectre of Too Many Species. Syst. Biol. 68:168–181.

Lewis P.O. 2001. A likelihood approach to estimating phylogeny from discrete morphological character data. Syst. Biol. 50:913–925.

Louis E.E.L., Engberg S.E., McGuire S.M., McCormick M.J., Randriamampionona R., Ranaivoarisoa J.F., Bailey C.A., Mittermeier R.A., Lei R. 2008. Revision of the Mouse Lemurs, Microcebus (Primates, Lemuriformes), of Northern and Northwestern Madagascar with Descriptions of Two New Species at Montagne d’Ambre National Park and Antafondro Classified Forest. Primate Cons. 23:19–38.

Maestri R., Luza A.L., de Barros L.D., Hartz S.M., Ferrari A., de Freitas T.R.O., Duarte L.D.S. 2016. Geographical variation of body size in sigmodontine rodents depends on both environment and phylogenetic composition of communities. Journal of Biogeography. 43:1192–1202.

Malécot G. 1948. Les mathématiques de l’hérédité. Barnéoud frères.

Marquart K. 2014. Habitat use and morphological adaptations of endemic rodents (Muroidea: Nesomyinae) of East Madagascar. Universität Hohenheim, Munich.

Mason N.A., Fletcher N.K., Gill B.A., Funk W.C., Zamudio K.R. 2020. Coalescent-based species delimitation is sensitive to geographic sampling and isolation by distance. System. Biodivers. 18:269–280.

Mayr E. 1956. Geographical Character Gradients and Climatic Adaptation. Evolution. 10:105–108.

Meiri S., Dayan T. 2003. On the Validity of Bergmann’s Rule. Journal of Biogeography. 30:331–351.

Meyer C.P., Paulay G. 2005. DNA Barcoding: Error Rates Based on Comprehensive Sampling. PLoS Biol. 3:e422.

Michaux J., Reyes A., Catzeflis F. 2001. Evolutionary History of the Most Speciose Mammals: Molecular Phylogeny of Muroid Rodents. Mol. Biol. Evol. 18: 2017–2031.

Milne Edwards A., 1885. Description d’une nouvelle espèce de rongeur provenant de Madagascar. Ann. Sci. nat., Zoo. Paléontol.

Muldoon K.M., de Blieux D.D., Simons E.L., Chatrath P.S. 2009. The Subfossil Occurrence and Paleoecological Significance of Small Mammals at Ankilitelo Cave, Southwestern Madagascar. J. Mammal. 90:1111–1131.

Musser, G.G., Carleton, M.D., 2005. Superfamily Muroidea., in: Mammal Species of the World: A Taxonomic and Geo-Graphical Reference. Johns Hopkins University Press, Baltimore, pp. 894–1531.

Nei M. 1972. Genetic Distance between Populations. Am. Nat. 106:283–292.

Paradis E., Schliep K. 2019. ape 5.0: an environment for modern phylogenetics and evolutionary analyses in R. Bioinformatics. 35:526–528.

Paris J.R., Stevens J.R., Catchen J.M. 2017. Lost in parameter space: a road map for stacks. Methods Ecol. Evol. 8:1360–1373.

Pinheiro J., Bates D. 2000. Mixed-Effects Models in S and S-PLUS. Springer Science & Business Media.

Pinheiro J., Bates D., R Core Team. 2022. nlme: Linear and Nonlinear Mixed Effects Models. R package version 3.1-158.

Pinho C., Hey J. 2010. Divergence with Gene Flow: Models and Data. Annu. Rev. Ecol. Evol. Syst. 41:215–230.

Poux C., Madsen O., Marquard E., Vieites D.R., de Jong W.W., Vences M. 2005. Asynchronous colonization of Madagascar by the four endemic clades of primates, tenrecs, carnivores, and rodents as inferred from nuclear genes. Syst. Biol. 54: 719–730.

Pritchard J.K., Stephens M., Donnelly P. 2000. Inference of population structure using multilocus genotype data. Genetics 155:945–959.

Purcell S., Neale B., Todd-Brown K., Thomas L., Ferreira M.A.R., Bender D., Maller J., Sklar P., de Bakker P.I.W., Daly M.J., Sham P.C. 2007. PLINK: A Tool Set for Whole-Genome Association and Population-Based Linkage Analyses. Am. J. Hum. Genet. 81:559–575.

Quéméré E., Amelot X., Pierson J., Crouau-Roy B., Chikhi L. 2012. Genetic data suggest a natural prehuman origin of open habitats in northern Madagascar and question the deforestation narrative in this region. Proc. Natl. Acad. Sci. U.S.A. 109:13028–13033.

R Core Team 2022. R: A language and environment for statistical computing. R Foundation for Statistical Computing, Vienna, Austria. https://www.R-project.org/

Rakotoarisoa J.E., Raheriarisena M., Goodman S.M. 2013a. Late Quaternary climatic vegetational shifts in an ecological transition zone of northern Madagascar: insights from genetic analyses of two endemic rodent species. J. Evol. Biol. 26:1019–1034.

Rakotoarisoa J.E., Raheriarisena M., Goodman S.M. 2013b. A phylogeographic study of the endemic rodent Eliurus carletoni (Rodentia: Nesomyinae) in an ecological transition zone of Northern Madagascar. J. Hered. 104:23–35.

Rakotondravony R., Radespiel U. 2009. Varying patterns of coexistence of two mouse lemur species (Microcebus ravelobensis and M. murinus) in a heterogeneous landscape. Am. J. of Primatol. 71:928–938.

Rannala B., Yang Z. 2020. Species delimitation. In: Scornavacca C., Delsuc F., Galtier N., editors. Phylogenetics in the Genomic Era. p. 1–18.

Rannala B., Yang Z. 2003. Bayes Estimation of Species Divergence Times and Ancestral Population Sizes Using DNA Sequences From Multiple Loci. Genetics 164:1645–1656.

Rasolonjatovo S.M., Scherz M.D., Hutter C.R., Glaw F., Rakotoarison A., Razafindraibe J.H., Goodman S.M., Raselimanana A.P., Vences M. 2020. Sympatric lineages in the Mantidactylus ambreensis complex of Malagasy frogs originated allopatrically rather than by in-situ speciation. Mol. Phylogenet. Evol. 144:106700.

Raxworthy C.J., Nussbaum R.A. 1997. Biogeographic patterns of reptiles in eastern Madagascar. In: Goodman S.M., Patterson B.D. editors. Natural Change and Human Impact in Madagascar. Washington, D.C.: Smithsonian Institution Press. p. 124–141.

Raxworthy C.J., Nussbaum R.A. 1995. Systematics, speciation and biogeography of the dwarf chameleons (Brookesia; Reptilia, Squamata, Chamaeleontidae) of northern Madagascar. J. Zool. 235:525–558.

Raxworthy C.J., Nussbaum R.A. 1994. A rainforest survey of amphibians, reptiles and small mammals at Montagne d’Ambre, Madagascar. Biol. Conserv. 69:65–73.

Rochette N.C., Catchen J.M. 2017. Deriving genotypes from RAD-seq short-read data using Stacks. Nat. Protoc. 12:2640–2659.

Rochette N.C., Rivera-Colón A.G., Catchen J.M. 2019. Stacks 2: Analytical methods for paired-end sequencing improve RADseq-based population genomics. Mol. Ecol. 28:4737–4754.

Rodríguez M.Á., Olalla-Tárraga M.Á., Hawkins B.A. 2008. Bergmann’s rule and the geography of mammal body size in the Western Hemisphere. Global Ecology and Biogeography. 17:274–283.

Romano A., Séchaud R., Roulin A. 2020. Geographical variation in bill size provides evidence for Allen’s rule in a cosmopolitan raptor. Global Ecology and Biogeography. 29:65–75.

Romiguier J., Ranwez V., Douzery E.J.P., Galtier N. 2010. Contrasting GC-content dynamics across 33 mammalian genomes: relationship with life-history traits and chromosome sizes. Genome Res. 20:1001–1009.

Ronquist F., Teslenko M., van der Mark P., Ayres D.L., Darling A., Höhna S., Larget B., Liu L., Suchard M.A., Huelsenbeck J.P. 2012. MrBayes 3.2: efficient Bayesian phylogenetic inference and model choice across a large model space. Syst. Biol. 61:539–542.

Rosen D.E. 1979. Fishes from the uplands and intermontane basins of Guatemala: revisionary studies and comparative geography. Bull. Am. Mus. Nat. Hist. 162:267–376.

Rousset 2000. Genetic differentiation between individuals. J. Evol. Biol. 13:58–62.

Rousset F. 1997. Genetic Differentiation and Estimation of Gene Flow from F-Statistics Under Isolation by Distance. Genetics 145:1219–1228.

Rubin B.E.R., Ree R.H., Moreau C.S. 2012. Inferring Phylogenies from RAD Sequence Data. PLOS ONE 7:e33394.

Salmona J., Heller R., Quéméré E., Chikhi L. 2017. Climate change and human colonization triggered habitat loss and fragmentation in Madagascar. Mol. Ecol. 26:5203–5222.

Seutin G., White B.N., Boag P.T. 1991. Preservation of avian blood and tissue samples for DNA analyses. Can. J. Zool. 69:82–90.

Sexton J.P., Hangartner S.B., Hoffmann A.A. 2014. Genetic Isolation by Environment or Distance: Which Pattern of Gene Flow Is Most Common? Evolution. 68:1–15.

Sgarlata G.M., Salmona J., Pors B.L., Rasolondraibe E., Jan F., Ralantoharijaona T., Rakotonanahary A., Randriamaroson J., Marques A.J., Aleixo-Pais I., Zoeten T. de, Ousseni D.S.A., Knoop S.B., Teixeira H., Gabillaud V., Miller A., Ibouroi M.T., Rasoloharijaona S., Zaonarivelo J.R., Andriaholinirina N.V., Chikhi L. 2019. Genetic and morphological diversity of mouse lemurs (Microcebus spp.) in northern Madagascar: The discovery of a putative new species? Am. J. Primatol. 81:e23070.

Shi J.J., Chan L.M., Rakotomalala Z., Heilman A.M., Goodman S.M., Yoder A.D. 2013. Latitude drives diversification in Madagascar’s endemic dry forest rodent Eliurus myoxinus (subfamily Nesomyinae). Biol. J. Linn. Soc. 110:500–517.

Sievers F., Wilm A., Dineen D., Gibson T.J., Karplus K., Li W., Lopez R., McWilliam H., Remmert M., Söding J., Thompson J.D., Higgins D.G. 2011. Fast, scalable generation of high-quality protein multiple sequence alignments using Clustal Omega. Mol. Syst. Biol. 7:539.

Simons E.L., Burney D.A., Chatrath P.S., Godfrey L.R., Jungers W.L., Rakotosamimanana B. 1995. AMS 14C Dates for Extinct Lemurs from Caves in the Ankarana Massif, Northern Madagascar. Quat. Res. 43:249–254.

Skotte L., Korneliussen T.S., Albrechtsen A. 2013. Estimating individual admixture proportions from next generation sequencing data. Genetics 195:693–702.

Soarimalala V., Goodman S.M. 2003. Diversité biologique des micromammifères non volants (Liptotyphla et Rodentia) dans le complexe Marojejy-Anjanaharibe-Sud. In: Goodman S.M., Wilmé, L., editors. Nouveaux résultats d’inventaires biologiques faisant référence à l’altitude dans la région des massifs montagneux de Marojejy et d’Anjanaharibe-Sud: Recherches pour le Développement, Série Sciences Biologiques. p. 231–278.

Stamatakis A. 2014. RAxML version 8: a tool for phylogenetic analysis and post-analysis of large phylogenies. Bioinformatics 30:1312–1313.

Steppan S.J., Schenk J.J., 2017. Muroid rodent phylogenetics: 900-species tree reveals increasing diversification rates. PLOS ONE 12: e0183070.

Struck T.H., Feder J.L., Bendiksby M., Birkeland S., Cerca J., Gusarov V.I., Kistenich S., Larsson K.H., Liow L.H., Nowak M.D., Stedje B., Bachmann L., Dimitrov D. 2018. Finding Evolutionary Processes Hidden in Cryptic Species. Trends Ecol. Evol. 33:153–163.

tattersall I. 2007. Madagascar’s Lemurs: Cryptic diversity or taxonomic inflation? Evol. Anthropol. 16:12–23.

tattersall I., Cuozzo F.P. 2019. Systematics of the extant Malagasy lemurs (order Primates). In: Goodman S.M., Raherilalao M.J., Wohlhause S. editors. The Terrestrial Protected Areas of Madagascar: Their History, Description, and Biota. Antananarivo: Association Vahatra. p. 403–424.

tysdal J.A., Jansa S.A. 2014. Phylogeny and biogeography of an endemic Madagascar rodent.

Vinogradov A.E. 1998. Genome size and GC-percent in vertebrates as determined by flow cytometry: the triangular relationship. Cytometry. 31:100–109.

Webb C.S. 1954. The odyssey of an animal collector. London: Longmans

Weisrock D.W., Rasoloarison R.M., Fiorentino I., Ralison J.M., Goodman S.M., Kappeler P.M., Yoder A.D. 2010. Delimiting Species without Nuclear Monophyly in Madagascar’s Mouse Lemurs. PLOS ONE. 5:e9883.

Wiemers M., Fiedler K. 2007. Does the DNA barcoding gap exist? – a case study in blue butterflies (Lepidoptera: Lycaenidae). Front. Zool. 4:1–16.

Wright S. 1943. Isolation by Distance. Genetics. 28:114–138.

Yang Z. 2015. The BPP program for species tree estimation and species delimitation. Curr. Zool. 61:854–865.

Yang Z., Rannala B. 2017. Bayesian species identification under the multispecies coalescent provides significant improvements to DNA barcoding analyses. Mol. Ecol. 26:3028–3036.

Yang Z., Rannala B. 2010. Bayesian species delimitation using multilocus sequence data. Proc. Natl. Acad. Sci. U.S.A. 107: 9264–9269.

Yoder A.D., Campbell C.R., Blanco M.B., Reis M. dos, Ganzhorn J.U., Goodman S.M., Hunnicutt K.E., Larsen P.A., Kappeler P.M., Rasoloarison R.M., Ralison J.M., Swofford D.L., Weisrock D.W. 2016. Geogenetic patterns in mouse lemurs (genus Microcebus) reveal the ghosts of Madagascar’s forests past. Proc. Natl. Acad. Sci. U.S.A. 113:8049–8056.

Yu G. 2020. Using ggtree to Visualize Data on Tree-Like Structures. Curr. Protoc. Bioinformatics. 69:e96.

Yu G., Smith D.K., Zhu H., Guan Y., Lam T.T.-Y. 2017. ggtree: an r package for visualization and annotation of phylogenetic trees with their covariates and other associated data. Methods Ecol. Evol. 8:28–36.

