## Supplementary Appendix S1 for "The Genomic Diversity of the *Eliurus* genus in northern Madagascar with a Putative New Species"

Data available from the Dryad Digital Repository: http://dx.doi.org/10.5061/dryad.[NNNN]

*Study Region*

*North of the Loky river*.—At the north of the Loky river, there are mainly fragmented dry forests, frequently surrounded by riparian corridors along narrow canyons or by more extended riparian forests in proximity of the Irodo river. In this region, we mainly visited forest sites within three major massifs: Ankarana, Analamerana and Andrafiamena-Andavakoeira. The Special Reserve (SR) of Ankarana (IUCN category IV) is a limestone massif of (~250 km$^2$) covered mostly by pinnacle karst ("tsingy") and dry forests. However, a large diversity of vegetation types can be also observed, for instance, moist primary seasonal forest and specialized xerophytic scrub (Rossi 1974; Fowler et al. 1989). Surface water is rare and only present along canyons, where underground rivers get to the surface. The thick soil on the basalt formation of the massif, the underground water and the protection from drying winds create optimal conditions for the persistence of moist and humid vegetation within canyons (Fowler et al. 1989; Nicoll and Legrand 1989). There are also humid deciduous forests on the basaltic soil of the northern side of massif, extending down from Montagne d'Ambre (Goodman et al., 2021a). The surface area of the moist forests is limited to 20 – 50 km$^2$, representing a 'refugium' during the dry season for some animal taxa (e.g., lemurs) that, then during the wet season, also use more extensively the deciduous forests (Hawkins et al. 1990). The SR of Ankarana is currently managed by Madagascar National Parks (Goodman et al., 2021a). The SR of Analamerana (~470 km$^2$) is characterised by the same vegetational types as Ankarana Special Reserve. It is drier than Ankarana forests and lies entirely on limestone. The dominant vegetation is dry deciduous forest. Moist semi-deciduous vegetation can be found near river streams, with tree species typical from rainforest habitats (Hawkins et al. 1990). Two major blocks of forests can be identified within the Analamerana SR, one in the northeast that is well preserved, and one in the southwest which has been heavily fragmented by fires (Goodman et al., 2021b). The Analamerana SR is currently managed by Madagascar National Parks. The Andrafiamena-Andavakoeira Protected Harmonious Landscape (IUCN category V; ~730 km$^2$) is composed of two large forests, Andrafiamena and Andavakoeira differing in geological substrate types: limestone karst and sandstone, respectively (Buřivalová 2011). As a consequence, also the vegetation occurring in the two forests is different. In fact, Buřivalová (2011) sampled large trees in forested habitats found on both substrates and determined that only 4% of the species were shared between the two forests. In particular, Andrafiamena forest has an alkaline soil layer, and is more water deprived and support a more xeric and deciduous vegetation than Andavakoeira forest (Fowler et al. 1989), which occurs on sandstone, where trees are less water deprived and often closely related to the tree species occurring in the eastern rainforest. The Andrafiamena-Andavakoeira Protected area is managed by the NGO Fanamby.

*Loky-Manambato region.*—The Loky-Manambato (LM) region is a biogeographical transition zone between dry deciduous and humid forests (Goodman and Wilmé 2006), which is delimited by the Loky and Manambato rivers in the north and south, respectively. This region is crossed by the relatively shallow Manankolana River, bordered by riparian forests along most of its course, and by a national dirt road. It consists of an area of ~2,500 km$^2$ covered by ~360 km$^2$ of forests (Goodman et al., 2021c), fragmented into a dozen major forest patches surrounded by human-altered grasslands, dry scrub and agricultural lands. Most forests are situated at low- to mid-elevations and mostly consist of dry deciduous vegetation. In contrast, some mountain forests on the southern edge (Binara and Antsahabe, plus Bobankora to a lower extent) are covered by a gradient of dry deciduous, transition, humid and ericoid vegetation (Gautier et al. 2006). The LM region is protected as a Protected

Harmonious Landscape (IUCN category V) and managed by the Malagasy NGO "Fanamby" since 2005 (Goodman et al., 2021, pp. 528–545).

*South Manambato region.*— In this region, forest fragments are currently un-protected (Goodman et al., 2021d). Between the Manambato and the Manambery rivers, there are the lowland dry deciduous forest of Analafiana and the mountain humid forest of Salafaina, which topology and vegetation are similar to the Binara massif of the LM region. Between the Manambery and the Fanambana rivers, there is the evergreen humid forest of Bezavona-Ankirendrina. Ultimately, the southernmost visited forest of Analalava, south of the Fanambana River, is an evergreen humid forest. Little detailed ecological and vegetation information on these forest fragments can be found in the literature. However, recurrent survey by members of our research team have observed over years, a decrease in habitat amount and quality (Salmona J. and Le Pors B. pers. comm.).

*COMATSA region.*— Within what we call ‘COMATSA region’, we surveyed sites that are located in the continuous forest of the Corridor of Marojejy- Anjanaharibe Sud-Tsaratanana Nord protected area (COMATSA Nord) and those lying in the small unprotected forest fragments, at the north-east of the protected area (Ankinjanala and Anketrakabe). The vegetation composition of the COMATSA region is poorly studied. What is currently known is that it is mostly covered by continuous medium altitude evergreen forests, which only in few locations of the north-central portion transit to an ericoid vegetation. The forest of the Andravory Massif is isolated from the continuous forest by a 3-4 km stretch of secondary grassland along the Fanambana River (Goodman et al., 2021e). The Ankinjanala and Anketrakabe forests are disconnected from the north-east portion of the COMATSA protected area by secondary grassland, with only some riparian forests along river streams. The COMATSA Nord protected area is classified as a Natural Resources Reserve (VI IUCN category) and is currently managed by WWF (Goodman et al., 2021e).

*North Marojejy.*— We visited three sites that are located in the unprotected forests at the north of the Anakarangona river and Marojejy National Park: Lohananjialava, Beamalonkely, Andranomenabe, Ambatomainty, Andohanibemarivo. They are moist semi-deciduous forest fragments separated by secondary grassland. Deforestation in this area is most likely recent.

*Montagne d'Ambre.*— Montagne d'Ambre is an isolated volcanic massif with a peak of altitude at 1475 m. It is placed in a dry ecoregion, but due to its exposition to the trade winds, it receives more precipitation than neighbouring areas. Most of the massif is covered by medium-altitude moist evergreen forest, and only close to the summit the moist vegetation tends to transit to an ericoid vegetation (Nicoll and Legrand 1989; Raxworthy and Nussbaum 1994). Montagne d'Ambre is protected under the status of National Park (IUCN category II) and is managed by Madagascar National Parks (Goodman et al., 2021f).

*Study Species*

*E. carletoni* (Goodman et al. 2009) is a relatively large member of the *Eliurus* genus (head-body length: 143 – 150 mm; Goodman et al. 2009) known to occur in the extreme north of Madagascar (Fig. S17). It inhabits the relatively mesic forest conditions between the pinnacle karsts (called in Malagasy *tsingy*) (Bésairie 1964; Puy and Moat 1996; Veress et al. 2008) found at the SR of Ankarana, SR of Analamerana and a nearby zone of metamorphic rock in the PHL-V of Loky-Manambato (Daraina region) (Rakotoarisoa et al. 2010, 2013). *E. carletoni* can be considered a scansorial forest-dwelling species of dry deciduous habitats and it has been caught in both undisturbed and disturbed forests. Little is known on the ecology of this species. Documented records report an elevation range of occurrence between 20 m and 500 m (Jansa et al. 2019). *E. carletoni*, as also its sister-species *E. antsingy*, is characterised by a tuft that cover 45% – 55% of tail length. The tuft is dark blackish – brown and monochromatic over the entire tail (Goodman et al. 2009; Jansa et al. 2019). The dorsal pelage is typically dark-brown, which contrasts with the grayish-white ventral pelage. The tail length and pilosity distinguish *E. carletoni* from other *Eliurus* species with similarly monochromatic dark-blackish brown tails (*E. myoxinus* and *E. webbi*) (Carleton 1994; Carleton et al. 2001; Carleton 2003; Goodman et al. 2009).

*E. ellermani* (Carleton 1994) is another large member of the *Eliurus* genus (head-body length: 126 – 174 mm; Jansa et al. 2019) restricted to the north and north-eastern montane humid forests of Madagascar (Fig. S17). In particular, individuals of *E. ellermani*, whose species identity has been genetically confirmed, have been collected from the northernmost Montagne d’Ambre to the central mountains of the Masoala Peninsula. The habitat of *E. ellermani* consists of evergreen humid forest in montane formations, characterised by moist or sclerophyllous plant communities. It occurs in an elevation range between 440 m and 1570 m (Jansa et al. 2019). *E. ellermani* composes, together with *E. tanala* and *E. tsingimbato*, the so-called *E. tanala* species group. The three species are morphological difficult to distinguish, and also 'remarkably similar morphologically to members of the *E. antsingy* group (*E. antsingy* and *E. carletoni*) (Jansa et al. 2019). The tuft covers about 35% – 40% of tail length. The traits that most distinguish *E. ellermani* from the other species of the *E. tanala* group is the bicolored hairs and overall grayish appearance of the ventral pelage in *E. tanala*, in contrast to monochromatic ventral hairs and creamy white underparts in *E. ellermani.* The tail tuft with a brown tip distinguishes in most cases *E. tsingimbato* from *E. ellermani* (Jansa et al. 2019).

*E. minor* (Major 1896) is the smallest member of the *Eliurus* genus (head-body length: 101 – 124 mm; Goodman and Carleton 1996) (Fig. S17). The litter size of *E. minor* ranges from 2 to 4 (Carleton 2003). The breeding season of this species occurs in the last quarter of the year (October to December; Goodman and Carleton 1996; Carleton and Goodman 1998; Goodman et al. 1999). However, this information is limited by the biased time period in which fieldwork data have been collected. Earlier sparse observations (Carleton 1994) showed a male with scrotal testes recorded for May, a specimen in juvenile pelage for July, an embryo count of two for August and a lactating female for August. Trap capture data suggest that *E. minor* can be found on lianas, small trees, bamboo grass, on tree ferns as well as on dead wood, filigree structures or trees with a small trunk diameter. It has been observed to climb along filigree vegetation, lianas and in dense filigree undergrowth (Marquart 2014), up to heights of several meters (Goodman and Carleton 1996; Carleton and Goodman 1998). Observations by Katrin Marquart (Marquart 2014) suggests that in comparison to other *Eliurus* species (*E. tanala* and *E. webbi*), *E. minor* can be 'considered both a good large-tree-climbers as well as a good 'small-branch and bamboo- grass-climbers'. *E. minor* occurs in a wide elevation range (near sea level to 1875 m; Garbutt 1999; Carleton 2003) in correspondence of moist evergreen forests. Individuals whose species identity has been genetically confirmed show that *E. minor* has a wide distribution, extending from the south-east to the north-east of Madagascar (Everson et al. 2020). Morphologically, *E. minor* can be distinguished from the other *Eliurus* species by its smaller cranial and body-size, the smallest of the genus. In addition, it is characterised by a typically dark-brown or blackish tuft-tail, a bright brown colouration of the dorsal pelage and the lack of a marked difference in dorsal-ventral coloration. The tuft covers between 60% - 66% of the tail (Carleton 1994).

*E. myoxinus* (Milne Edwards 1885) is a medium-size tuft-tailed rat (head-body length: 117 – 136 mm; Carleton et al. 2001) (Fig. S17). It occurs in the dry forests of the west and in the lowland humid forests of the far north/north-east Madagascar, which have a pronounced annual dry season (Soarimalala and Goodman 2003). Litter size between 1- 3 (Carleton 2003). It has been documented from localities in xerophytic scrub (spiny bush), dry deciduous forest, dry-humid transitional forest (Carleton 1994; Goodman and Ganzhorn 1994; Goodman and Rasoloarison 1997; Goodman et al. 1999). It is strictly forest-dependent and occurs in dry and humid forests up to 900 m and 1240 m in elevation, respectively (Soarimalala and Goodman 2011). *E. myoxinus* individuals have been collected also from the mesic (moist) canyons of Isalo and the upper portion of the Analavelona Massif, which have distinctly more humid vegetation than surrounding lowland areas (Goodman et al. 1999; Carleton et al. 2001; Soarimalala and Goodman 2003; Goodman et al. 2009; Shi et al. 2013). *E. myoxinus* is characterised by a relatively shorter tail length compared to the other *Eliurus* species (105% - 110% of head-body length). *E. myoxinus*, although being a western-dry associated species, has a body size smaller than any of the eastern-humid *Eliurus* species. It is covered by a generally lighter brown dorsal pelage. *E. myoxinus* individuals present a decline in body size, going from the south to the north of its distribution, e.g., head-body-length is 127.7 mm $\pm$ 6.4 in the south and 123.6 mm $\pm$ 8.7 in the north (Table 3 in Carleton et al.

2001). The tuft is brown to brownish-black, long and bushy, covering between 60% - 70% of the tail (Carleton 1994).

*RAD-seq Data Processing and de novo Assembly*

The *ustacks* program is run on each individual in the dataset, separately, to build loci from single-end short-reads. It can be distinguished in two steps which allows to i) create *stacks* (or putative alleles) of exactly matching reads; and ii) match putative alleles (*stacks*) into a locus. These two steps are controlled by two parameters: *m* (minimum depth of coverage), corresponding to the minimum number of raw reads required to form a *stack*; and *M* (maximum distance between stacks), equivalent to the maximum number of nucleotides differences allowed between stacks to merge them into a putative locus.

The *de-novo* analysis creates a 'synthetic' reference-genome (*catalog*) which is made of all loci and alleles assembled in the *ustacks* program across the individuals of the dataset. In other words, it creates a set of consensus loci by merging homologous alleles of different individuals together. The key parameter in this step is *n* (distance allowed between *catalog* loci), corresponding to the number of mismatches allowed between sample loci when building the *catalog* consensus loci. This parameter is fundamental since it determines the threshold for considering a given allele a homologous of other alleles in the whole dataset.

The program *sstacks* is used to match individuals' loci to the *catalog*, and loci matching more than one *catalog* locus are excluded because their true matching locus in the *catalog* is ambiguous, either due sequencing error or to true paralogy. Multiple loci, however, can still uniquely match the same *catalog* locus, however later analyses (in *gstacks* and *populations* program) can exclude these loci (Catchen et al. 2011). The *tsv2bam* program rearrange the format of the matched loci and retrieve the paired-end reads associated with the single-end reads that have been used for individuals' loci assembly. The *gstacks* program assembles the paired-end reads retrieved in the previous step into contigs which are then merged with the assembled single-end loci. After that, *gstacks* calls SNPs within the whole dataset using the information on the matched loci, and genotype each individual accordingly. For *gstacks*, we used a --*var-alpha* and --*gt-alpha* equal to 0.01. The *var-alpha* parameter defines the p-value threshold for calling a given nucleotide *polymorphic* rather than *invariant* across all individuals in the dataset. The *gt-alpha* parameter defines the p-value threshold of the likelihood-ratio test statistic between the significant heterozygous genotype (called depending on *var-alpha*) and the genotype homozygous for the more abundant nucleotide in the individual at a given site (Maruki and Lynch 2017). If the likelihood of the heterozygous genotype is significantly greater than the likelihood of the homozygous genotype, then the heterozygous genotype is considered, otherwise a homozygous genotype is called. In practice, a *gt-alpha* parameter of 0.01 would be equivalent to a Genotype Quality score $\geq 20$, calculated as $-\log_{10}(\text{likelihood ratio test p} - \text{value})$.

The *gstacks* program generates the consensus *catalog* FASTA file which correspond to the 'synthetic' reference genome used for phylogenomic analyses. The *catalog* FASTA file was cleaned from possible contaminant sequences using *DeconSeq* software (Schmieder and Edwards 2011). We decided to use *DeconSeq* on the *catalog* loci and not on the raw reads, because it requires long-read datasets (> 150 bp mean read length) and also because several steps in the STACKS v2.2 pipeline are made such to reduce the probability of calling nucleotides that are rarely represented in the dataset. Briefly, *DeconSeq* assesses sequence similarity between *catalog* loci and Bacteria, Virus and Human databases of potential contaminants. It uses the parameters *coverage* and *identity*, that is how many nucleotides of a *catalog* locus are covered by and identical to the reference contaminant sequence, respectively. We set *coverage* equal to 95 and *identity* equal to 94. Finally, we used the *populations* program to analyse the genomic data from the whole dataset allowing to convert this information, for instance, into a dataset of single SNPs per locus or to FASTA sequences. In doing so, additional filtering parameters need to be specified, which allow to further increase confidence on the recovered genomic information by reducing biases due to sequencing errors, paralogy or repetitive regions. In the *populations* program, as conditions for processing a nucleotide at a locus, we set i) a minimum minor allele frequency (*min_maf*) equal to 5%; ii) a maximum observed heterozygosity (*max_obs_het*) equal to 70%; iii) restrict data analysis to one random SNP per locus, in order to reduce linkage disequilibrium among genotyped SNPs (*write-random-snp*); and iv) a minimum percentage of individuals in a population required to process a locus equal to 80% (-*r*).

*RAD-seq Datasets Building*

Here we give a detailed description of the criteria of selection. In particular, we generated four types of datasets, differing in the number and geographical distribution of the individuals: i) all 124 individuals (*N124*); ii) one individual per sampling site, per species (*N62*); iii) 10 individuals per species (*N48*); and iv) 3 individuals per species (*N15*). For each dataset, we created a *white list*, which included loci that were genotyped: *i*) if at least 80% of the individuals across taxa contained that locus (*pop1.r80*); or *ii*) if at least 80% of the individuals within each taxon contained that locus (*pop5.p0.r80*); or *iii*) if at least 80% of the individuals within each taxon and 80% of the taxa (N = 4) contained that locus (*pop5.p4.r80*).

The *white lists* were then used for generating the genomic datasets on which to perform phylogenomic and genomic analyses. Most of the datasets used for phylogenomic analysis (maximum-likelihood tree inference) were composed of sites that are fixed within individuals but variables across individuals, since we are first interested in the topology of the species tree, and not in the branch lengths. For two datasets (*N124.pop5.p4.min100* and *N15.pop5.p4.min15*), we also generated a multialignment composed of all sites (variant + fixed). The *N124.pop5.p4.min100* dataset was used for computing average genetic difference between and within taxa, whereas the *N15.pop5.p4.min15* dataset was used for phylogenetic inference, species delimitation test and estimation of divergence times and species population size (see below). Finally, we generated a dataset, with all individuals across taxa, composed by one random SNPs per *catalog* locus, which needs to be present in at least 80% of the individuals within taxa and in at least four out of the five taxa (*N124.pop5.p4.r80*).

For comparison, we also generated one genomic dataset (*N124.pop5.p4.r80*) using a genotype-likelihood (GL) based approach implemented in ANGSD (Nielsen et al. 2012; Korneliussen et al. 2014). One difference between ANGSD and STACKS v2.2 is that the former does not call genotypes but estimates the posterior probability of all possible genotypes, given the data. Using the GLs based approach would allow thus to strengthen results by using a larger amount of information (Heller et al. 2021). The ANGSD pipeline was run on the full dataset (N=124) by first aligning cleaned sequences to the *de-novo catalog* generated in STACKS v2.2. The alignment was carried out with *bwa-mem* (Li 2013). We estimated locus depth as the F1 read starting position (Sbf1 cutting site) using the "samtools-view" command (Li et al. 2009). We then estimated genotype likelihoods with the SAMtools approach ('-GL 1') using only loci with *i*) a total depth of at least five (-setMinDepth), *ii*) a total depth of at most the sum of the 0.95 quantiles of the depth distribution of considered individuals (-setMaxDepth), and *iii*) an individual depth of at most the 0.95 quantiles of its depth distribution (-setMaxDepthInd). This strategy allows discarding loci present only in a very low number of individuals, or over-covered loci that are likely to be repeated or paralog regions (Heller et al. 2021). In addition, we considered only bases with a minimum quality of 20 and paired reads that mapped to one unique *catalog* locus with a minimum quality of 30. Finally, we only kept biallelic variants with a SNP probability lower than 1e-6, a minor allele frequency (MAF) of at least 0.05, and that were present in at least 100 out of 124 individuals.

*Mitochondrial DNA Analyses*

In MRBAYES v3.2.1, we used four MCMC chains, each at the default temperature and running for 15,000,000 generations. Trees were sampled every 15,000 generations and 25% of total sampled trees were discarded as the "burn-in." To assess MCMC convergence, we checked for stationarity of the log probability of the data over time, we verified that the potential scale reduction factor was close to 1 and we used Tracer v1.6 (Rambaut et al. 2014) to confirm that all estimated sample size values were >200. In RAXML v8.2.X, we used the rapid Bootstrap option with automatic detection of the sufficient number of bootstrap replicates (option: "autoMR"). Retained posterior distributions of trees are summarized in MRBAYES to build a consensus tree, while in RAXML majority-rule consensus tree is estimated from the Bootstrap replicates. We estimated the best partitioning scheme and best-fit model of evolution for mtDNA *cytb* in PARTITIONFINDER v2.1.1 (Lanfear et al. 2017) using the greedy algorithm and selecting the best model according to AICc criteria. We performed a separate analysis for each phylogenetic method, given that MRBAYES and RAXML accept only some of the models of evolution that can be estimated in PARTITIONFINDER.

**REFERENCES**


Bésairie H. 1964. Carte géologique à 1:1.000.000 de Madagascar. Tananarive: Serv. Geol. Madagascar.

Buřivalová Z. 2011. Remote sensing of vegetation in conservation: A case study from the dry and transitional forests of Andrafiamena, northern Madagascar. Master's dissertation, Universite de Geneve and Conservatoire et Jardin botaniques de la Ville de Geneve, Genève.

Carleton M., Goodman S. 1998. New taxa of nesomyine rodents (Muroidea: Muridae) from Madagascar's northern Highlands, with taxonomic comments on previously described forms. In: Goodman S.M. editor. A floral and faunal inventory of the Réserve Spéciale d' Anjanaharibe-Sud, Madagascar: with reference to elevational variation. Zoology, New Series No. 90 Fieldiana. p. 163–200.

Carleton M.D. 1994. Systematic studies of Madagascar's endemic rodents (Muroidea: Nesomyinae): revision of the genus *Eliurus*. Am. Mus. Novit. 3087:1–55.

Carleton M.D., Goodman S.M., Rakotondravony D. 2001. A new species of tufted-tailed rat, genus *Eliurus* (Muridae: Nesomyinae), from western Madagascar, with notes on the distribution of *E. myoxinus*. Proc. Biol. Soc. Wash. 114:972–987

Carleton M.D. 2003. *Eliurus*, tufted-tailed rats. In: Goodman S.M., Benstead J.P. editors. The Natural History of Madagascar. Chicago: University of Chicago Press. p. 1373–1380.

Carleton M.D., Goodman S.M., Rakotondravony D. 2001. A new species of tufted-tailed rat, genus *Eliurus* (Muridae: Nesomyinae), from western Madagascar, with notes on the distribution of *E. myoxinus*. Proc. Biol. Soc. Wash. 114:972–987

Catchen J.M., Amores A., Hohenlohe P., Cresko W., Postlethwait J.H. 2011. Stacks: Building and Genotyping Loci De Novo From Short-Read Sequences. G3 (Bethesda). 1:171–182.

Du Puy D.J., Moat J. 1996. A refined classification of the primary vegetation of Madagascar based on the underlying geology: Using GIS to map its distribution and to assess its conservation status. In: Lourenco, W.R., editor. Biogeography of Madagascar. Paris: ORSTOM. p. 205–218.

Everson K.M., Goodman S.M., Olson L.E. 2020. Speciation and gene flow in two sympatric small mammals from Madagascar, *Microgale fotsifotsy* and *M. soricoides* (Mammalia: Tenrecidae). Mol. Ecol. 29:1717–1729.

Fowler S.V., Chapman P., Checkley D., Hurd S., McHale M., Ramangason G.-S., Randriamasy J.-E., Stewart P., Walters R., Wilson J.M. 1989. Survey and management proposals for a tropical deciduous forest reserve at Ankarana in Northern Madagascar. Biol. Conserv. 47:297–313.

Garbutt N. 1999. Mammals of Madagascar. Yale University Press.

Gautier L., Ranirison P., Nusbaumer L., Wohlhauser S. 2006. Aperçu des massifs forestiers de la région Loky-Manambato. In: Goodman S.M. and Wilmé L. editors. Inventaires de La Faune et de La Flore Du Nord de Madagascar Dans La Région Loky-Manambato, Analamerana et Andavakoera. Rech. Développement. Sér. Sci. Biol. 23:81–99

Goodman S.M., Carleton M.D. 1996. The rodents of the Réserve Naturelle Intégrale d'Andringitra, Madagascar. In: Goodman S.M. editor. A floral and faunal inventory of the eastern slopes of the Réserve Naturelle Intégrale d'Andringitra, Madagascar: with reference to elevational variation. Zoology, New Series No. 85 Fieldiana. p. 257–283.

Goodman S.M., Carleton M.D., Pidgeon M. 1999. The rodents of the Réserve Naturelle Intégrale d'Andohahela, Madagascar. In: Goodman S.M. editor. A floral and faunal inventory of the eastern slopes of the Réserve Naturelle Intégrale d'Andohahela, Madagascar: with reference to elevational variation. Zoology, New Series No. 94 Fieldiana. p. 217–249.

Goodman S.M., Ganzhorn J.U. 1994. Les petits mammiferes. In: Goodman S.M., Langrand O. editors. Inventaire biologique: Foret de Zombitse. Recherches pour le developpement, serie sciences biologiques. Antananarivo, Madagascar: Centre d'Information et de Documentation Scientifique et Technique. p. 58–63.

Goodman S.M., Raheriarisena M., Jansa S.A. 2009. A new species of Eliurus Milne Edwards, 1885 (Rodentia: Nesomyinae) from the Réserve Spéciale d'Ankarana, northern Madagasca. Bonn. Zool. Beitr. 56:133–149.

Goodman S.M., Raherilalao M.J., Wohlhauser S. 2021a. Site 8. Ankarana. In: Goodman S.M., Raherilalao M.J., Wohlhauser S. editors. The Terrestrial Protected Areas of Madagascar: Their History, Description, and Biota, Volume 2: Northern and eastern Madagascar - Synthesis. Antananarivo: Association Vahatra. p. 558–572.

Goodman S.M., Raherilalao M.J., Wohlhauser S. 2021b. Site 5. Analamerana. In: Goodman S.M., Raherilalao M.J., Wohlhauser S. editors. The Terrestrial Protected Areas of Madagascar: Their History, Description, and Biota, Volume 2: Northern and eastern Madagascar - Synthesis. Antananarivo: Association Vahatra. p. 515–527.

Goodman S.M., Raherilalao M.J., Wohlhauser S. 2021c. Site 6. Loky Manambato. In: Goodman S.M., Raherilalao M.J., Wohlhauser S. editors. The Terrestrial Protected Areas of Madagascar: Their History, Description, and Biota, Volume 2: Northern and eastern Madagascar - Synthesis. Antananarivo: Association Vahatra. p. 528–545.

Goodman S.M., Raherilalao M.J., Wohlhauser S. 2021d. The Terrestrial Protected Areas of Madagascar: Their History, Description, and Biota, Volume 2: Northern and eastern Madagascar - Synthesis. Antananarivo: Association Vahatra.

Goodman S.M., Raherilalao M.J., Wohlhauser S. 2021e. Site 11. COMATSA Nord. In: Goodman S.M., Raherilalao M.J., Wohlhauser S. editors. The Terrestrial Protected Areas of Madagascar: Their History, Description, and Biota, Volume 2: Northern and eastern Madagascar - Synthesis. Antananarivo: Association Vahatra. p. 597–611.

Goodman S.M., Raherilalao M.J., Wohlhauser S. 2021f. Site 4. Montagne d'Ambre. In: Goodman S.M., Raherilalao M.J., Wohlhauser S. editors. The Terrestrial Protected Areas of Madagascar: Their History, Description, and Biota, Volume 2: Northern and eastern Madagascar - Synthesis. Antananarivo: Association Vahatra. p. 500–514.

Goodman S.M., Rasoloarison R. 1997. Les petits mammiferes. In: Goodman S.M., Langrand O. editors. Inventaire biologique: Foret de Vohibasia et d'Isoky-Vohimena. Recherches pour le developpement, serie sciences biologiques. Antananarivo, Madagascar: Centre d'Information et de Documentation Scientifique et Technique. p. 144–155.

Goodman S.M., Wilmé L. 2006. Inventaires de la faune et de la flore du nord de Madagascar dans la région Loky-Manambato, Analamerana et Andavakoera. Rech. dév., Sér. Sci. biol. 23:1–238.

Hawkins A., Chapman P., Ganzhorn J., Bloxam Q., Barlow S., Tonge S. 1990. Vertebrate conservation in Ankarana special reserve, Northern Madagascar. Biol. Conserv. 54:83–110

Heller R., Nursyifa C., Garcia-Erill G., Salmona J., Chikhi L., Meisner J., Korneliussen T.S., Albrechtsen A. 2021. A reference-free approach to analyse RADseq data using standard next generation sequencing toolkits. Mol. Ecol. Resour. 21:1085–1097.

Jansa S.A., Carleton M.D., Soarimalala V., Rakotomalala Z., Goodman S.M. 2019. A review of the *Eliurus tanala* complex (Rodentia, Muroidea, Nesomyidae), with description of a new species from dry forests of western Madagascar. Bull. Am. Mus. Nat. Hist. 430:1–69.

Korneliussen T.S., Albrechtsen A., Nielsen R. 2014. ANGSD: Analysis of Next Generation Sequencing Data. BMC Bioinformatics. 15:356.

Lanfear R., Frandsen P.B., Wright A.M., Senfeld T., Calcott B. 2017. PartitionFinder 2: New Methods for Selecting Partitioned Models of Evolution for Molecular and Morphological Phylogenetic Analyses. Mol. Biol. Evol. 34:772–773.

Li H. 2013. Aligning sequence reads, clone sequences and assembly contigs with BWA-MEM. arXiv:1303.3997.

Li H., Handsaker B., Wysoker A., Fennell T., Ruan J., Homer N., Marth G., Abecasis G., Durbin R., 1000 Genome Project Data Processing Subgroup. 2009. The Sequence Alignment/Map format and SAMtools. Bioinformatics. 25:2078–2079.

Major C.I.F. 1896. Descriptions of four additional new mammals from Madagascar. Ann. Mag. Nat. Hist.108:461–463.

Marquart K. 2014. Habitat use and morphological adaptations of endemic rodents (Muroidea: Nesomyinae) of East Madagascar.

Maruki T., Lynch M. 2017. Genotype Calling from Population-Genomic Sequencing Data. G3: Genes, Genomes, Genet. 7:1393–1404.

Milne Edwards A., 1885. Description d’une nouvelle espèce de rongeur provenant de Madagascar. Ann. Sci. nat., Zoo. Paléontol.

Nicoll M.E., Legrand. 1989. Madagascar : revue de la conservation et des aires protégées. Gland: WWF.

Nielsen R., Korneliussen T., Albrechtsen A., Li Y., Wang J. 2012. SNP Calling, Genotype Calling, and Sample Allele Frequency Estimation from New-Generation Sequencing Data. PLOS ONE. 7:e37558.

Rakotoarisoa J.E., Raheriarisena M., Goodman S.M. 2010. Phylogeny and species boundaries of the endemic species complex, *Eliurus antsingy* and *E. carletoni* (Rodentia: Muroidea: Nesomyidae), in Madagascar using mitochondrial and nuclear DNA sequence data. Mol. Phylogenet. Evol. 57:11–22.

Rakotoarisoa J.E., Raheriarisena M., Goodman S.M. 2013. Late Quaternary climatic vegetational shifts in an ecological transition zone of northern Madagascar: insights from genetic analyses of two endemic rodent species. J. Evol. Biol. 26:1019–1034.

Rambaut A., Drummond A.J., Xie D., Baele G., Suchard M.A. 2014. Tracer v1.6. http://beast. bio.ed.ac.uk

Raxworthy C.J., Nussbaum R.A. 1994. A rainforest survey of amphibians, reptiles and small mammals at Montagne d'Ambre, Madagascar. Biol. Conserv. 69:65–73.

Rossi G. 1974. Morphologie et évolution d’un karst en milieu tropical: l’Ankarana (Madagascar). Mém. et Doc. du C.N.R.S. vol. 15. Phénomènes karstiques II. Paris: p. 279–298.

Schmieder R., Edwards R. 2011. Fast Identification and Removal of Sequence Contamination from Genomic and Metagenomic Datasets. PLOS ONE. 6:e17288.

Shi J.J., Chan L.M., Rakotomalala Z., Heilman A.M., Goodman S.M., Yoder A.D. 2013. Latitude drives diversification in Madagascar’s endemic dry forest rodent Eliurus myoxinus (subfamily Nesomyinae). Biol. J. Linn. Soc. 110:500–517.

Soarimalala V., Goodman S.M. 2003. Diversité biologique des micromammifères non volants (Liptotyphla et Rodentia) dans le complexe Marojejy-Anjanaharibe-Sud. In: Goodman S.M., Wilmé, L., editors. Nouveaux résultats d'inventaires biologiques faisant référence à l'altitude dans la région des massifs montagneux de Marojejy et d'Anjanaharibe-Sud: Recherches pour le Développement, Série Sciences Biologiques. p. 231–278.

Soarimalala V., Goodman S.M. 2011. Les Petits Mammifères de Madagascar. Antananarivo, Madagascar: Association Vahatra.

Veress M., Lóczy D., Zentai Z., Tóth G., Schläffer R. 2008. The origin of the Bemaraha tsingy (Madagascar). Int. J. of Speleol. 37:131–142.
