## Supplementary Appendix S2 for "The Genomic Diversity of the *Eliurus* genus in northern Madagascar with a Putative New Species"

Data available from the Dryad Digital Repository: http://dx.doi.org/10.5061/dryad.[NNNN]

**Table S1. List of *Eliurus* individuals used for the parameter's optimization analysis and for the construction of the *de-novo* catalog.**

| Forest | Site | ID | Altitude (m) | NFWReads* | Nloci* | Coverage | WCoverage | Species |
|---|---|---|---|---|---|---|---|---|
| Mahamasina/Ankarana | MAHAR | MAHAR.F19 | 110 | 2,988 | 149 | 20.0 | 21.3 | *E. carletoni* |
| Salafaina | SAL | SAL.B64 | 336 | 4,923 | 147 | 33.4 | 35.4 | *E. carletoni* |
| Solaniampilana | SOL | SOL.K35 | 138 | 3,453 | 148 | 23.4 | 24.7 | *E. carletoni* |
| Andrafiamena | ANDF | ANDF.A11 | 408 | 5,024 | 149 | 33.7 | 36.0 | *E. carletoni* |
| Ambohitsitondroina | AMBO | AMBO.F57 | 132 | 2,948 | 148 | 19.9 | 21.0 | *E. carletoni* |
| Binara | BIN | BIN.K71 | 767 | 3,723 | 143 | 26.0 | 27.3 | *E. ellermani* |
| Antsahabe | ANTSB | ANTSB.H21 | 737 | 3,562 | 147 | 24.3 | 25.8 | *E. ellermani* |
| Anketrakabe | KABE | KABE.G09 | 818 | 1,886 | 142 | 13.3 | 13.9 | *E. ellermani* |
| Ankinjanala | ANKI | ANKI.G52 | 1,084 | 2,306 | 144 | 16.1 | 17.0 | *E. ellermani* |
| Ambalasoa | AMBALA | AMBALA.B25 | 918 | 1,756 | 132 | 13.3 | 13.9 | *E. ellermani* |
| Andoharano | ANDR | ANDR.E32 | 1,025 | 2,696 | 149 | 18.1 | 19.1 | *E. minor* |
| Ambalasoa | AMBALA | AMBALA.B72 | 1,075 | 1,812 | 143 | 12.7 | 13.4 | *E. minor* |
| Andoharano | ANDR | ANDR.E30 | 930 | 2,500 | 137 | 18.3 | 19.3 | *E. minor* |
| Maladialina | MAL | MAL.A69 | 1,014 | 3,433 | 153 | 22.4 | 24.0 | *E. minor* |
| Binara | BIN | BIN.BE78 | 822 | 4,531 | 147 | 30.8 | 32.8 | *E. myoxinus* |
| Andavakoera | TSAKA | TSAKA.D75 | 474 | 3,635 | 145 | 25.0 | 26.5 | *E. myoxinus* |
| Andrafiamena | ANDF | ANDF.B51 | 475 | 3,458 | 146 | 23.7 | 24.9 | *E. myoxinus* |
| Anketrakabe | KABE | KABE.G6 | 818 | 4,504 | 145 | 31.0 | 32.5 | *E. myoxinus* |
| Salafaina | SAL | SAL.B81 | 669 | 3,655 | 147 | 24.9 | 26.3 | *E. myoxinus* |
| Montagne d'Ambre | M_AMBRE_SOUTH | M_AMBRE_SOUTH.BE35 | 1,002 | 1,803 | 145 | 12.5 | 13.0 | *Eliurus* sp. nova |

Forest: forest name; Site: forest abbreviation; ID: identifier of the individual; Nloci: number of genotyped loci; Altitude: altitude of the capture site; NFWReads: number of forward reads; Coverage: mean coverage; WCoverage: weighted mean coverage, with more weight to loci that are more frequent; *: values are divided by $10^3$.

**Table S2. Accession numbers of the mtDNA cytb sequences used for mitochondrial phylogenetic analysis.**

| Species | N | Accession Numbers | References |
|---|---|---|---|
| *E. webbi* | 16 | AF160520-27; HM223718; KF058046 - 47; KF170838 - 42; KY753980 | Jansa et al., 1999; Rakotoarisoa et al., 2010; Shi et al., 2013; Steppan and Schenk, 2017 |
| *E. antsingy* | 6 | GQ420663 - 67; HM223711 | Goodman et al., 2009; Rakotoarisoa et al., 2010 |
| *E. grandidieri* | 3 | AF160571 - 2; KY753977 | Steppan and Schenk, 2017; Jansa et al., 1999 |
| *E. majori* | 16 | AF160549 - 60; GQ420668; HM223717; KF058048 - 49 | Jansa et al., 1999; Goodman et al., 2009; Rakotoarisoa et al., 2010; Shi et al., 2013 |
| *E. minor* | 11 | AF160539 - 48; KF058044; KF058050 | Jansa et al., 1999; Shi et al., 2013 |
| *E. tanala* | 14 | AF160530 - AF160538; KF058045; KY753979; MK806758; MK806763; MK806845 | Jansa et al., 1999; Shi et al., 2013; Steppan and Schenk, 2017; Jansa et al., 2019 |
| *E. tsingimbato* | 11 | MK806813; MK806816 - 17; MK806820 - 22; MK806825 - 26; MK806828; MK806832; MK806840 | Jansa et al., 2019 |
| *E. ellermani* | 38 | AF160528 - 29; AF160573;<br>KC433971 - 72, 74, 75, 77, 78, 80, 88, 96, 97, 99;<br>KC434002 - 04, 06, 07, 10, 24, 25, 26, 29, 31;<br>MK806723, 25, 27 - 29, 32, 34 - 36, 39 - 41, 43 | Jansa et al., 1999; Rakotoarisoa et al., 2013; Jansa et al., 2019 |
| *E. carletoni* | 67 | GQ420655, 57, 58, 61, 62;<br>HM223602 - 06, 08, 12, 13, 15, 16, 19, 21, 22, 26, 30, 32, 37, 38, 41, 43, 48, 49, 52, 54, 55, 66, 70, 71, 73, 74, 76, 78, 86 - 88, 91 - 93, 97;<br>HM223700, 04, 07;<br>HM223722; JQ866517 - 20, 26, 28, 36, 39 - 41, 43, 50, 55, 59, 64, 70, 71, 99; JQ866617 | Rakotoarisoa et al., 2010; Rakotoarisoa et al., 2013 |
| *E. myoxinus* | 60 | AF160561 - 63, 65, 67 - 69;<br>KF058051, 54 - 59, 61, 64, 68, 69, 74, 75, 78, 79, 85, 86, 88, 98, 99;<br>KF058104, 08, 10 - 12, 22 - 26, 39, 41, 42, 44 - 55, 67, 61, 62, 66 - 69;<br>KY753978 | Jansa et al., 1999; Shi et al., 2013; Steppan and Schenk, 2017 |

**Table S3. List of *Eliurus* individuals for which the mtDNA *cytb* has been sequenced.**

| Forest | Site | ID | CYTB (bp) | Species |
|---|---|---|---|---|
| Maladialina | MAL | MAL.A80 | 944 | *E. minor* |
| Ambalasoa | AMBALA | AMBAL.B78 | 928 | *E. minor* |
| Binara | BIN | BIN.K71 | 968 | *E. ellermani* |
| Anketrakabe | KABE | ANKET.G09 | 1,048 | *E. ellermani* |
| Ankinjanala | ANKI | ANKI.G52 | 991 | *E. ellermani* |
| Salafaina | SAL | SAL.B64 | 1,064 | *E. carletoni* |
| Analalava | ANALV | ANALV.A39 | 1,064 | *E. carletoni* |
| Montagne d'Ambre | M_AMBRE_NORTH | M_AMBRE_NORTH.BC51 | 1,067 | *Eliurus* sp. nova |
| Montagne d'Ambre | M_AMBRE_NORTH | M_AMBRE_NORTH.BD29 | 966 | *Eliurus* sp. nova |
| Montagne d'Ambre | M_AMBRE_SOUTH | M_AMBRE_SOUTH.BG77 | 956 | *Eliurus* sp. nova |
| Montagne d'Ambre | M_AMBRE_SOUTH | M_AMBRE_SOUTH.BG72 | 926 | *Eliurus* sp. nova |
| Montagne d'Ambre | M_AMBRE_SOUTH | M_AMBRE_SOUTH.BH10 | 952 | *Eliurus* sp. nova |
| Montagne d'Ambre | M_AMBRE_SOUTH | M_AMBRE_SOUTH.BE35 | 921 | *Eliurus* sp. nova |
| Montagne d'Ambre | M_AMBRE_SOUTH | M_AMBRE_SOUTH.BG61 | 919 | *Eliurus* sp. nova |
| Salafaina | SAL | SAL.B82 | 1,090 | *E. myoxinus* |
| Analalava | ANALV | ANALV.A28 | 993 | *E. myoxinus* |
| Bezavona - Ankirendrina | BEZ | BEZAV.C81 | 1,064 | *E. myoxinus* |
| Bezavona - Ankirendrina | BEZ | BEZAV.C58 | 1,064 | *E. myoxinus* |
| Bezavona - Ankirendrina | BEZ | BEZAV.C71 | 1,064 | *E. myoxinus* |
| Analafiana | ANALF | ANALF.B20 | 1,060 | *E. myoxinus* |

Forest: complete forest name; Site: forest abbreviation; ID: identifier of the individual; CYTB(bp): sequence length.

**Table S4. Nuclear RAD-seq datasets generated for phylogenomic and genomic analysis.**

| Dataset Name | White list | Sampling | Genotyping | N | #loci | #length (bp) | #geno/sites (bp) | #sites* | #variants* | #fixed* | pop rule | indv rule |
|---|---|---|---|---|---|---|---|---|---|---|---|---|
| N124.pop5.p4.min100 | pop5.p4.r80 | All | STACKS | 124 | 9,044 | 748 | 681 | 6,162 | 9 | 4 | 80% | 80% |
| N62.pop5.p0 | pop5.p0.r80 | 1 indv/site/species | STACKS | 62 | 71,879 | 740 | 626 | 44,977 | 72 | 36 | NA | NA |
| N62.pop5.p4 | pop5.p4.r80 | 1 indv/site/species | STACKS | 62 | 24,997 | 749 | 684 | 17,088 | 25 | 15 | 80% | NA |
| N62.pop5.p0.min50 | pop5.p0.r80 | 1 indv/site/species | STACKS | 62 | 22,289 | 746 | 628 | 14,000 | 22 | 12 | NA | 80% |
| N62.pop5.p4.min50 | pop5.p4.r80 | 1 indv/site/species | STACKS | 62 | 22,052 | 748 | 682 | 15,047 | 22 | 12 | 80% | 80% |
| N48.pop5.p0 | pop5.p0.r80 | 10 indv/species | STACKS | 48 | 89,210 | 742 | 636 | 56,776 | 89 | 44 | NA | NA |
| N48.pop5.p4 | pop5.p4.r80 | 10 indv/species | STACKS | 48 | 89,210 | 742 | 636 | 22,144 | 89 | 19 | 80% | NA |
| N48.pop5.p0.min38 | pop5.p0.r80 | 10 indv/species | STACKS | 48 | 25,884 | 745 | 635 | 16,449 | 26 | 13 | NA | 80% |
| N48.pop5.p4.min38 | pop5.p4.r80 | 10 indv/species | STACKS | 48 | 28,253 | 748 | 680 | 19,210 | 28 | 15 | 80% | 80% |
| N15.pop5.p4.min15 | pop5.p4.r80 | 3 indv/species | STACKS | 15 | 2,493 | 749 | 683 | 1,705 | 35 | 21 | 80% | 100% |
| N124.min100 | NA | All | ANGSD | 124 | 58,124 | NA | NA | 392 | NA | NA | NA | 80% |

*White list* indicates the lists of loci used for genotyping within each dataset. *pop5.p4.r80*: includes loci present in at least 80% of individuals within species (indv rule), and in at least four out of five species (Pop rule). *pop5.p0.r80*: includes loci present in at least 80% of individuals within species, with no constraints on the number of species in which a locus is present in order to be genotyped. Sampling indicates the criteria used to compose each dataset. *All*: all individuals; *1 indv/site/species*: one individual per sampling site per species; *10 indv/species* or *3 indv/species*: ten or three individuals per species. STACKS: defined genotype calls. ANGSD: genotype uncertainty is associated to each site. N: number of individuals included in the dataset. #loci: number of genotyped loci; #length (bp): mean length of the assembled loci; #geno/sites (bp): mean number of sites genotyped per locus; #sites*: total number of sites; #variants*: total number of SNPs. #fixed*: total number of fixed sites within but variable across individuals. * indicates that values are divided by$10^3$.

**Table S5. PGLS analysis on morphology and 20 (bio)climatic variables.**

| | Head | | | | Body | | | | Tail | | | | Weight | | | | Sign. |
|---|---|---|---|---|---|---|---|---|---|---|---|---|---|---|---|---|---|
| | slope | std-error | t-value | p-value | slope | std-error | t-value | p-value | slope | std-error | t-value | p-value | slope | std-error | t-value | p-value | |
| altitude | 0.00 | 0.00 | -0.56 | 0.58 | -0.01 | 0.00 | -1.98* | 0.05 | -0.03 | 0.01 | -3.50* | 0.00 | -0.02 | 0.01 | -3.41* | 0.00 | 3 |
| bio1 | 0.01 | 0.01 | 0.44 | 0.66 | 0.09 | 0.05 | 1.77 | 0.08 | 0.41 | 0.13 | 3.15* | 0.00 | 0.32 | 0.09 | 3.43* | 0.00 | 2 |
| bio2 | -0.03 | 0.04 | -0.74 | 0.46 | -0.34 | 0.13 | -2.62* | 0.01 | -1.25 | 0.32 | -3.88* | 0.00 | -0.83 | 0.23 | -3.62* | 0.00 | 3 |
| bio3 | -0.01 | 0.21 | -0.05 | 0.96 | -0.52 | 0.79 | -0.65 | 0.51 | 1.49 | 1.96 | 0.76 | 0.45 | 1.75 | 1.39 | 1.26 | 0.21 | 0 |
| bio4 | 0.00 | 0.00 | -0.27 | 0.79 | -0.01 | 0.01 | -1.59 | 0.11 | -0.04 | 0.01 | -2.84* | 0.00 | -0.04 | 0.01 | -4.50* | 0.00 | 2 |
| bio5 | 0.01 | 0.02 | 0.36 | 0.72 | 0.08 | 0.06 | 1.28 | 0.20 | 0.37 | 0.16 | 2.34* | 0.02 | 0.31 | 0.11 | 2.77* | 0.01 | 2 |
| bio6 | 0.01 | 0.01 | 0.55 | 0.58 | 0.09 | 0.04 | 2.06* | 0.04 | 0.39 | 0.11 | 3.53* | 0.00 | 0.29 | 0.08 | 3.74* | 0.00 | 3 |
| bio7 | -0.02 | 0.03 | -0.62 | 0.53 | -0.23 | 0.09 | -2.44* | 0.02 | -0.90 | 0.23 | -3.94* | 0.00 | -0.61 | 0.16 | -3.71* | 0.00 | 3 |
| bio8 | 0.01 | 0.02 | 0.55 | 0.58 | 0.11 | 0.06 | 1.87 | 0.06 | 0.48 | 0.15 | 3.24* | 0.00 | 0.35 | 0.11 | 3.35* | 0.00 | 2 |
| bio9 | 0.01 | 0.01 | 0.79 | 0.43 | 0.08 | 0.05 | 1.68 | 0.09 | 0.38 | 0.11 | 3.34* | 0.00 | 0.30 | 0.08 | 3.80* | 0.00 | 2 |
| bio10 | 0.01 | 0.02 | 0.47 | 0.64 | 0.11 | 0.06 | 1.76 | 0.08 | 0.46 | 0.15 | 3.15* | 0.00 | 0.33 | 0.10 | 3.21* | 0.00 | 2 |
| bio11 | 0.01 | 0.01 | 0.45 | 0.65 | 0.08 | 0.05 | 1.79 | 0.07 | 0.37 | 0.12 | 3.16* | 0.00 | 0.30 | 0.08 | 3.61* | 0.00 | 2 |
| bio12 | 0.00 | 0.00 | -1.53 | 0.13 | -0.01 | 0.01 | -0.58 | 0.56 | -0.04 | 0.03 | -1.32 | 0.19 | -0.04 | 0.02 | -1.88 | 0.06 | 0 |
| bio13 | 0.00 | 0.00 | -0.48 | 0.63 | 0.00 | 0.02 | 0.18 | 0.86 | 0.00 | 0.04 | 0.04 | 0.97 | 0.07 | 0.03 | 2.22* | 0.03 | 1 |
| bio14 | 0.00 | 0.03 | 0.14 | 0.89 | 0.07 | 0.13 | 0.57 | 0.57 | 0.17 | 0.32 | 0.52 | 0.61 | -0.60 | 0.23 | -2.66* | 0.01 | 1 |
| bio15 | 0.00 | 0.02 | 0.04 | 0.97 | 0.00 | 0.07 | -0.04 | 0.96 | -0.02 | 0.18 | -0.12 | 0.90 | 0.40 | 0.12 | 3.25* | 0.00 | 1 |
| bio16 | 0.00 | 0.00 | -0.77 | 0.44 | 0.00 | 0.01 | -0.25 | 0.80 | -0.01 | 0.02 | -0.55 | 0.58 | 0.02 | 0.02 | 1.38 | 0.17 | 0 |
| bio17 | 0.00 | 0.01 | 0.09 | 0.93 | 0.02 | 0.03 | 0.67 | 0.50 | 0.05 | 0.07 | 0.71 | 0.48 | -0.12 | 0.05 | -2.33* | 0.02 | 1 |
| bio18 | 0.01 | 0.00 | 2.05 | 0.04 | 0.01 | 0.01 | 0.82 | 0.41 | -0.04 | 0.03 | -1.39 | 0.17 | 0.07 | 0.02 | 3.34* | 0.00 | 1 |
| bio19 | 0.00 | 0.01 | 0.38 | 0.70 | 0.03 | 0.03 | 0.94 | 0.35 | 0.05 | 0.07 | 0.67 | 0.50 | -0.12 | 0.05 | -2.29* | 0.02 | 1 |

Head: head length; Body: body length; Tail: tail length; Sign: indicates the number of significant PGLS analysis across the four morphological variables.

**Table S6. Cases of sympatric occurrence in the *Eliurus* genus.**

| Species pairs | Region | Site | Reference |
|---|---|---|---|
| *E. antsingy - E. tanala* | West | / | Carleton et al., 2001 |
| *E. tsingimbato - E. antsingy* | North-Western, West | Foret d'Ambovonomby, Mahabo, Bekopaka | Jansa et al., 2019 |
| *E. tsingimbato - E. myoxinus* | West | / | Jansa et al., 2019 |
| *E. ellermani - E. carletoni* | North | Binara, Ambohibe | Jansa et al., 2019 |
| *E. ellermani - E. majori* | North | / | Jansa et al., 2019 |
| *E. tanala - E. majori* | Central Highlands | / | Jansa et al., 2019 |
| *E. tanala - E. minor* | Central Highlands | Perinet, Ranomafana | Carleton et al., 1994 |
| *E. minor - E. myoxinus* | North | Marojejy | Soarimalala and Goodman, 2003 |
| *E. tanala - E. webbi* | Central Highlands | Andrambovato, Ranomafana | Carleton et al., 1994 |
| *E. minor - E. webbi* | North-Eastern, South-Eastern | Maintimbato, Vondrozo | Carleton et al., 1994 |
| *Eliurus* sp. nova - *E. ellermani* | North | M. d'Ambre | Present study |
| *E. myoxinus - E. carletoni* | North | ANDF, SOL, SAL, ANALV, ANDRA | Present study |
| *E. myoxinus - E. carletoni - E. ellermani* | North | ANTSB, BIN, BEZ | Present study |
| *E. myoxinus - E. ellermani* | North | ANKI, KABE | Present study |
| *E. minor - E. ellermani* | North | AMBA, AMBALA | Present study |
| *E. myoxinus - E. carletoni - E. minor* | North | ANDR | Present study |

Species pairs: species in co-occurrence; Region: region of Madagascar in which co-occurrence has been documented; Site: locality of observation (forest or protected area); Reference: study where the co-occurrence has been documented or mentioned.
