## Supplementary Appendix S3 for "The Genomic Diversity of the *Eliurus* genus in northern Madagascar with a Putative New Species"

Data available from the Dryad Digital Repository: http://dx.doi.org/10.5061/dryad.[NNNN]

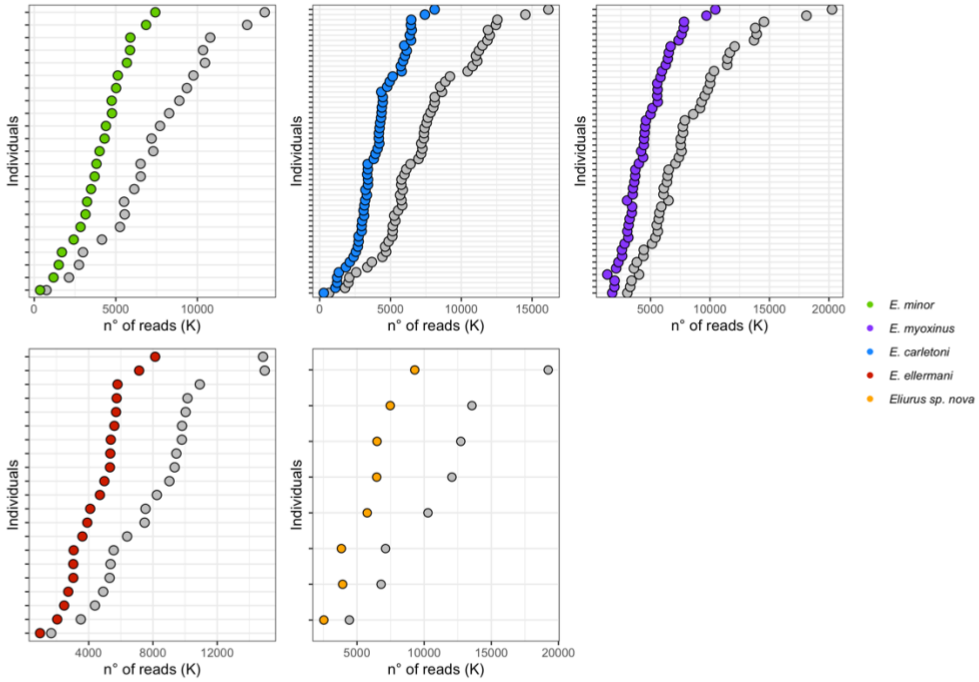


**Figure S1. Number of reads before and after filtering.** In the x-axis are indicated the number of reads and in the y-axis, individuals are ordered by increasing number of reads. Grey dots indicate number of reads before filtering.

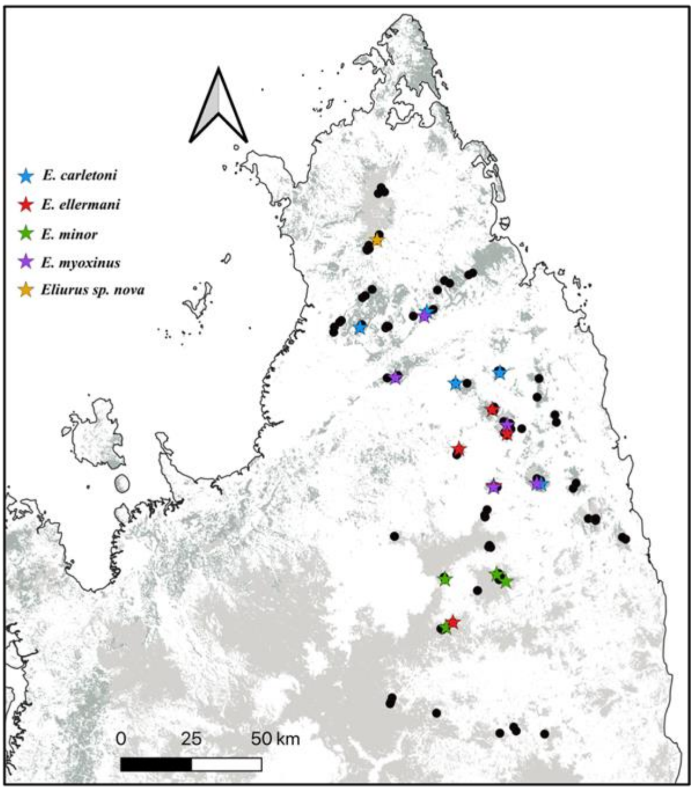


**Figure S2. *Eliurus* samples selected for building the *Eliurus* spp. catalog.** Black dots indicate the complete *Eliurus* dataset (N = 124), whereas coloured stars mark the individuals that have been selected for *catalog* assembly (Tab. S1). Individuals have been selected such to increase the geographic representation of each species, and use individuals with higher genomic coverage.

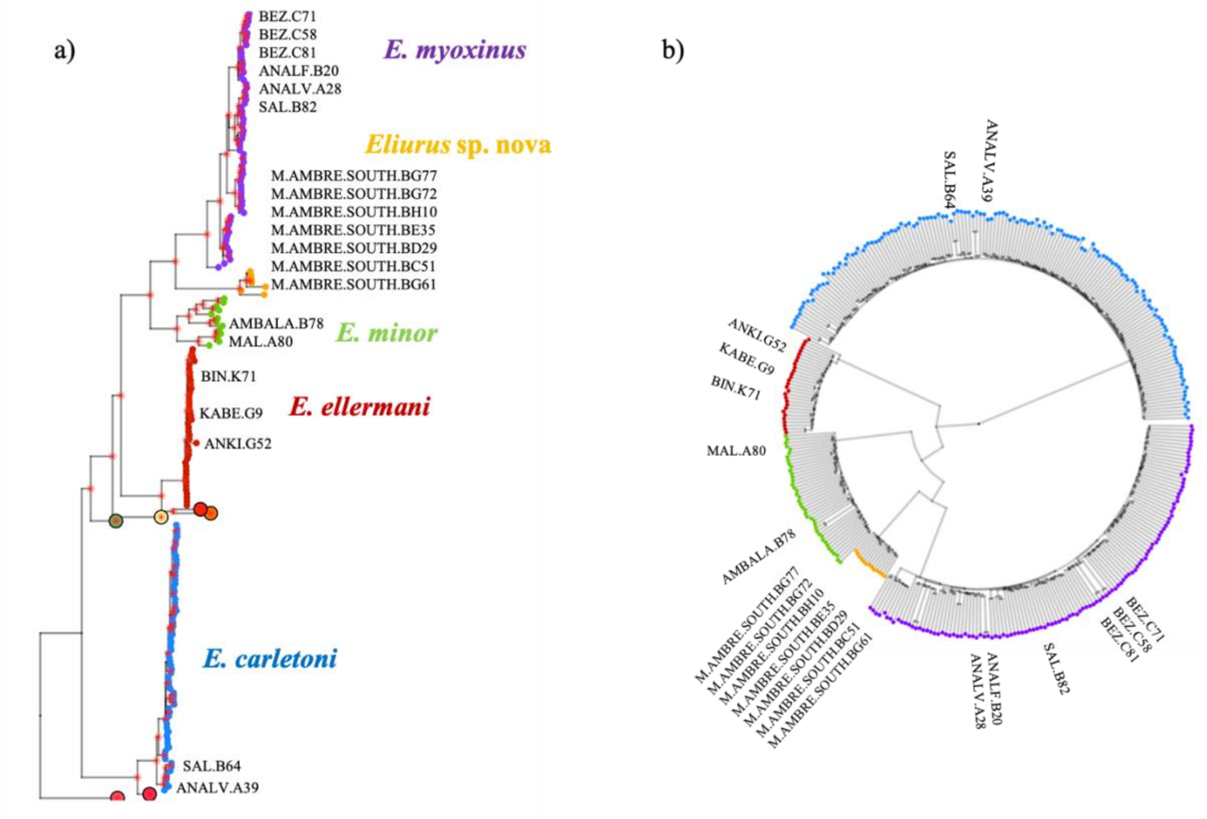


**Figure S3. Consistent inference of phylogenetic relationships from mtDNA *cytb* and nuclear genomic data.** Left: Bayesian mtDNA *cytb* phylogenetic tree of the *Eliurus* genus. The labelled individuals are the samples genotyped for *cytb* in the present study. Right: Neighbouring-joining tree of the RAD-seq nuclear data. This RAD-seq dataset included 347 individuals sampled across northern Madagascar, and loci that are present in at least 80% of the individuals (#SNPs: 8471). The concordant topological relationships recovered by the two phylogenetic trees suggest that our dataset is composed of five *Eliurus* taxa: *E. carletoni*; *E. myoxinus*; *E. minor*; *E. ellermani*; *Eliurus* sp. nova.

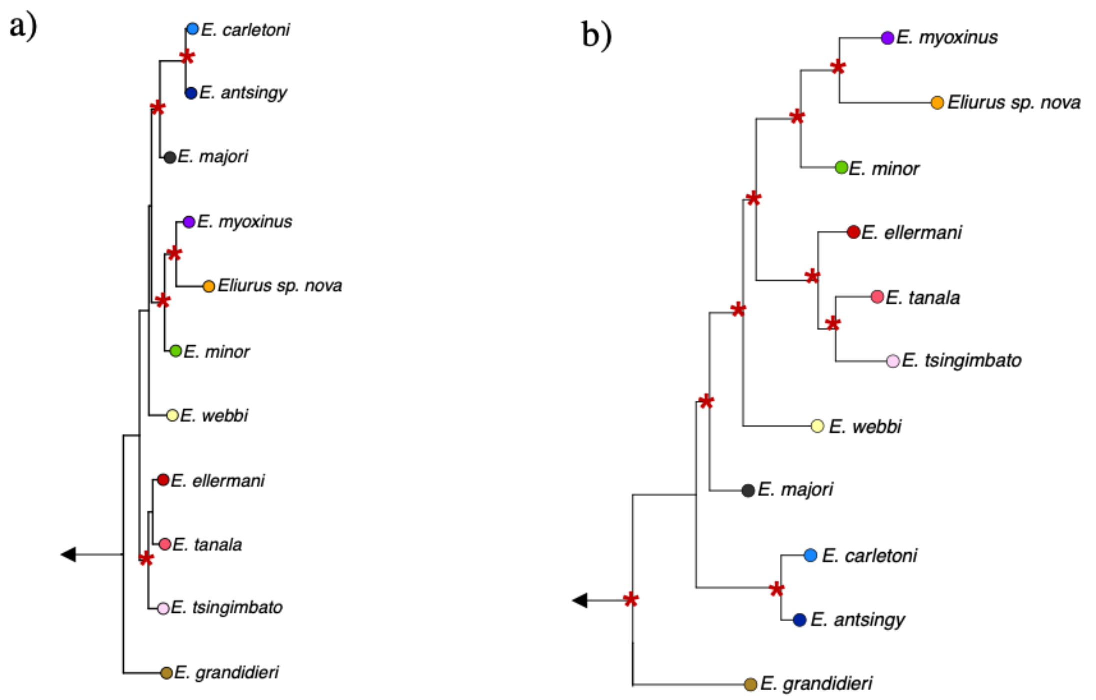


**Figure S4. Maximum-likelihood and Bayesian-based phylogenetic tree estimated from mtDNA *cytb*.** Left: Maximum-likelihood tree inferred in RAXML; Right: Bayesian-based tree inferred in MRBAYES. The two phylogenetic trees differ only on the topological relationships reconstructed at the deeper nodes, while confirming the phylogenetic units (taxa) identified in previous studies. MRBAYES phylogenetic tree is consistent with previous and well-established *Eliurus* phylogenies. Red asterisks indicate node support higher than 50%.

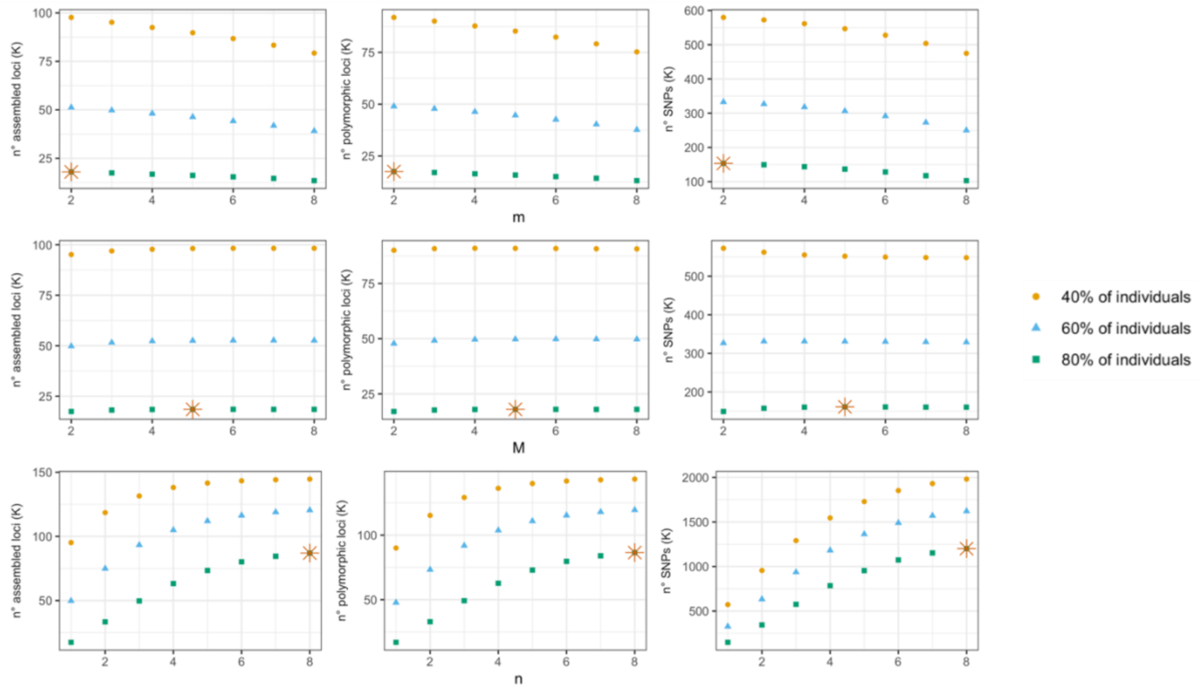


**Figure S5. Optimization of *Eliurus spp. de novo* assembly of RAD-seq loci using STACKS.** Iterating values of i) the minimum number of raw reads required to form a stack (*m*; First row), ii) the distance allowed between two stacks (*M*; Second row), and iii) the number of mismatches allowed between stacks during construction of the *catalog* (*n*; Third row). a) number of assembled loci; b) number of polymorphic loci; c) number of SNPs. The arrows indicate the best parameter combination, that is *m* = 2; *M* = 5 and *n* = 8.

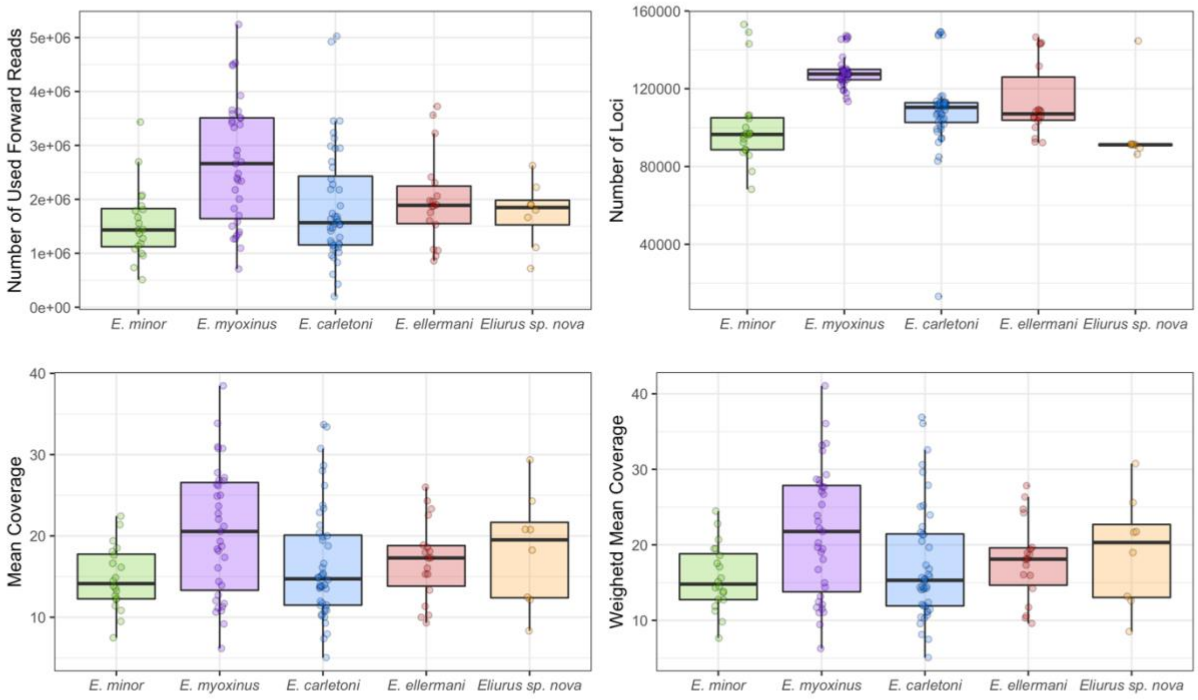


**Figure S6. Summary statistics of the assembled RAD-seq loci across *Eliurus* species.** Each panel shows individual variation within each *Eliurus* species in *i*) number of used forward reads; *ii*) number of assembled loci; *iii*) mean coverage; and *iv*) weighted mean coverage calculated by giving more weight to loci that are more represented across the dataset.

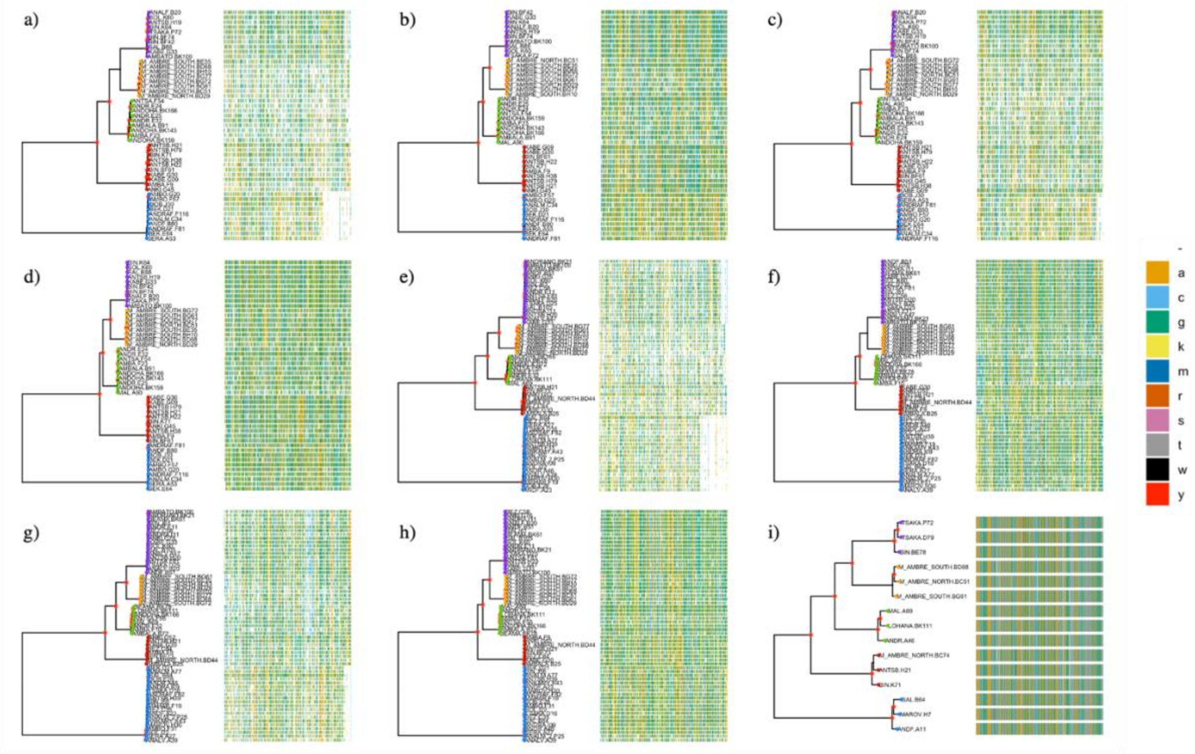


**Figure S7. Maximum-likelihood phylogenies inferred from concatenated sites in each RAD-seq datasets.** Phylogenies have been inferred using multialigments (on the right of each phylogeny) constructed by concatenating only sites that are fixed within individual and variants across individuals, with the exception of panel i) which includes all sites (variants and fixed). The phylogenies have been recovered using the following datasets (Tab. S4): a) N48.pop5.p0; b) N48.pop5.p0.min38; c) N48.pop5.p4; d) N48.pop5.p4.min38; e) N62.pop5.p0; f) N62.pop5.p0.min50; g) N62.pop5.p4; h) N62.pop5.p4.min50; i) N15.pop5.p4.min15. The legend indicates the color codes for each possible state in the alignment according to IUPAC nomenclature.

a)

b)

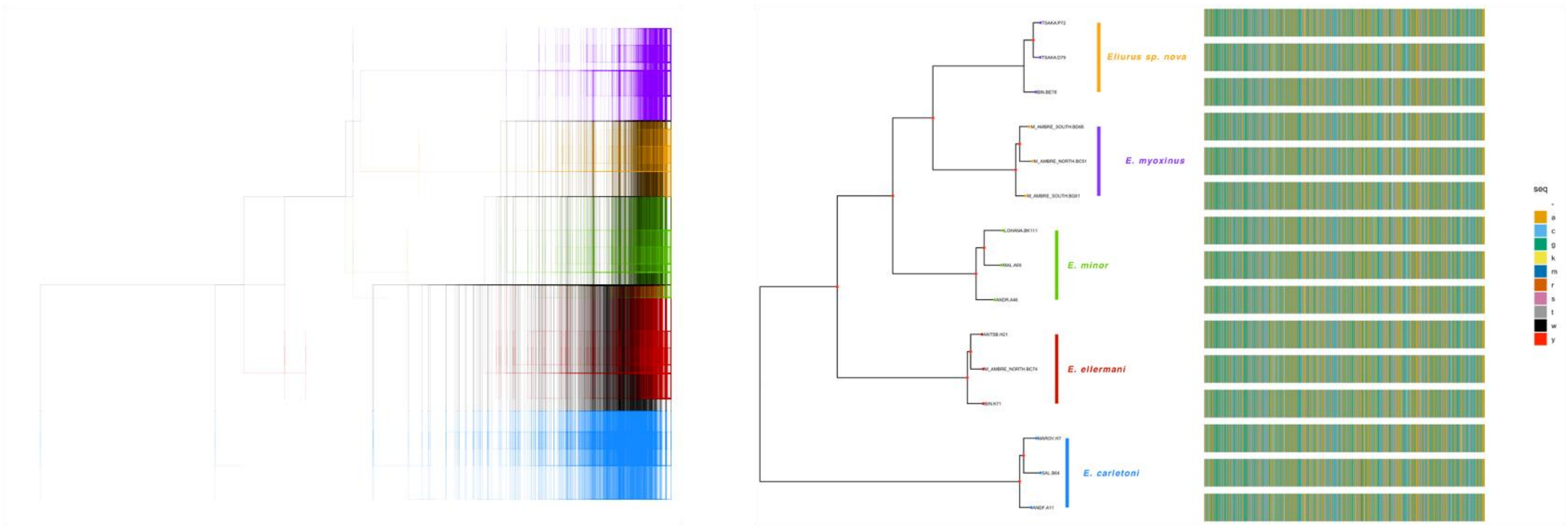


**Figure S8. Partitioned phylogenomic inference.** Maximum-likelihood phylogenies inferred for a) each partition (gene) independently, and b) using partitioned analysis, which estimates model parameters of the evolutionary model for each partition independently, while fixing the global set of branch lengths.

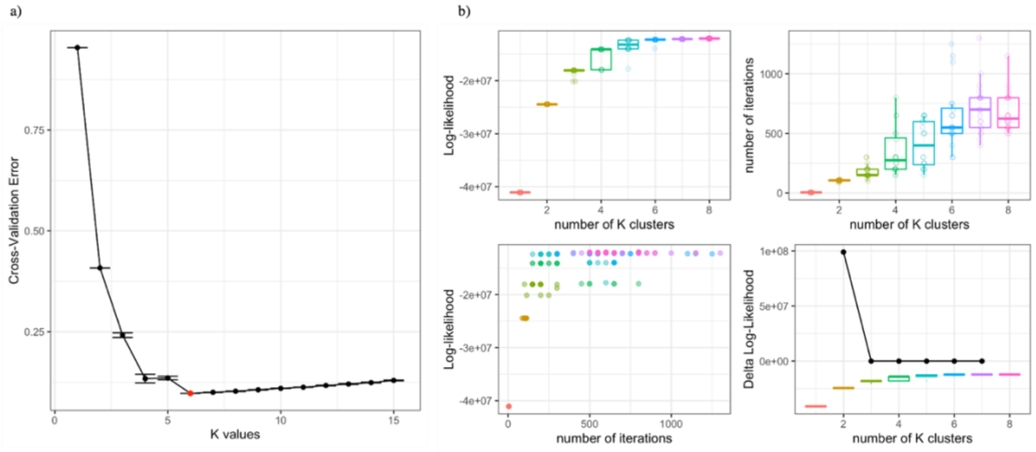


**Figure S9. ADMIXTURE cross validation error and NGSadmix log-likelihood across several K-cluster values.** a) ADMIXTURE model evaluation using 20-fold cross-validation (CV) error. The minimum CV error is identified for K = 6. b) Comparison of the log-likelihood of several K-values in NGS admix. The log-likelihood curve reaches plateau at K = 6. These results suggests that K = 6 is the minimum number of genetic cluster present in the dataset.

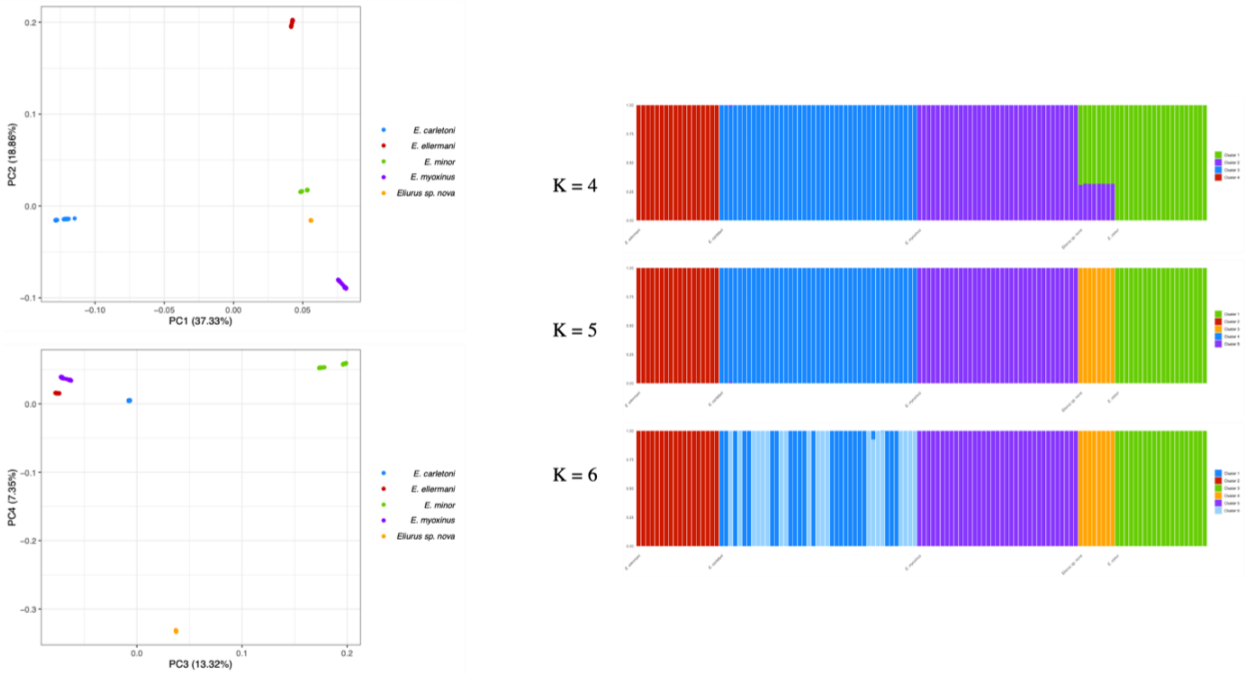


**Figure S10. *Eliurus* genomic structure using PCAangsd and NGSadmix.** PCA and ADMIXTURE analysis performed on genotype probabilities of the 124 *Eliurus* samples show five distinct clusters corresponding to the four previously identified species (*E. carletoni*; *E. myoxinus*; *E. ellermani*; *E. minor*) and the new taxon described in the present study (*Eliurus* sp. nova).

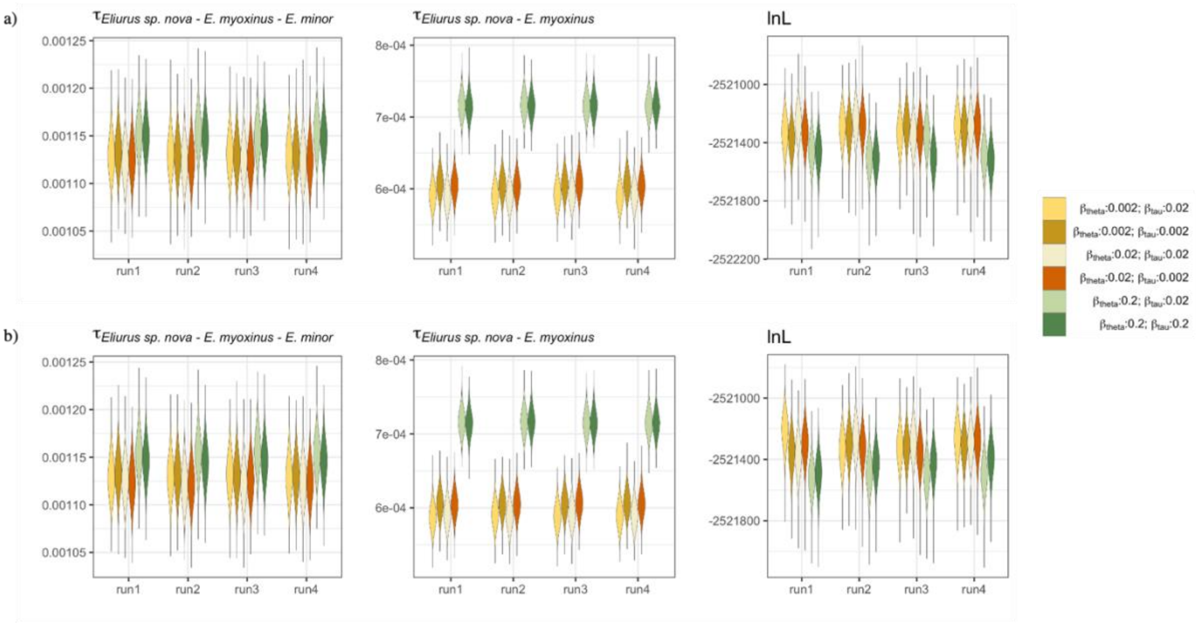


**Figure S11. MCMC samples for divergence times parameters across four MCMC runs of the species delimitation analysis.** Results obtained using algorithm0 (a) and algorithm1 (b) are concordant. The distribution of estimated $\tau_{E.myoxinus-Eliurus\ sp.nova-E.minor}$ values were similar across the six combinations of priors for divergence times and population size, whereas $\tau_{E.myoxinus-Eliurus\ sp.nova}$ showed significant differences for $\beta_{theta} = 0.2 - \beta_{tau} = 0.02$ and $\beta_{theta} = 0.2 - \beta_{tau} = 0.2$. Considering the large agreement across runs and algorithms, the results support the convergence of BPP analysis.

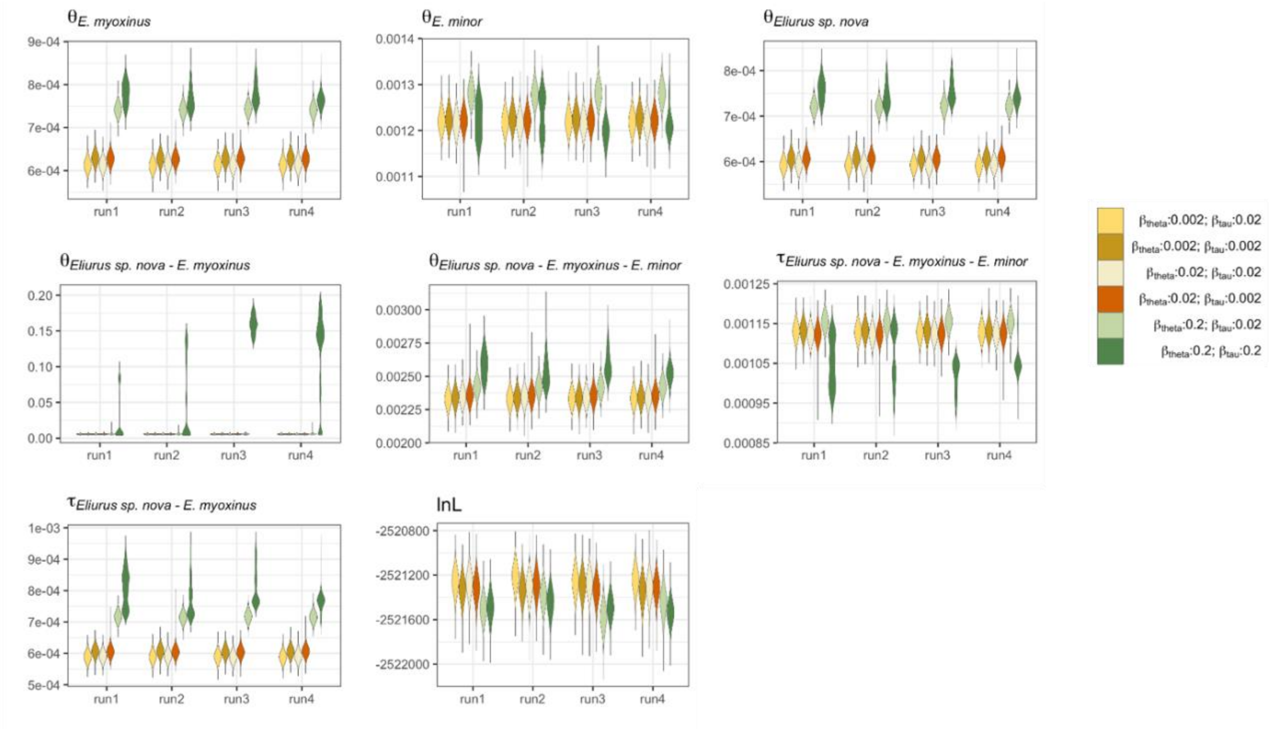


**Figure S12. Estimation of the divergence times and population size parameters under the multispecies-coalescent model for the hypothesized species phylogeny ((*E. myoxinus*, *Eliurus sp. nova*), *E. minor*).** The distribution of estimated values for $\theta_{E.minor}$, $\tau_{Eliurus\ sp.nova-E.myoxinus-E.minor}$, $\theta_{Eliurus\ sp.nova-E.myoxinus-E.minor}$, and log-likelihood were similar across the six combinations of priors and the four MCMC runs. $\theta_{E.myoxinus}$, $\tau_{E.myoxinus-Eliurus\ sp.nova}$, $\theta_{Eliurus\ sp.nova}$, $\theta_{Eliurus\ sp.nova-E.myoxinus}$ showed significant differences for $\beta_{theta} = 0.2 - \beta_{tau} = 0.02$ and $\beta_{theta} = 0.2 - \beta_{tau} = 0.2$. Considering the large agreement across runs and algorithms, the results support the convergence of BPP analysis,.

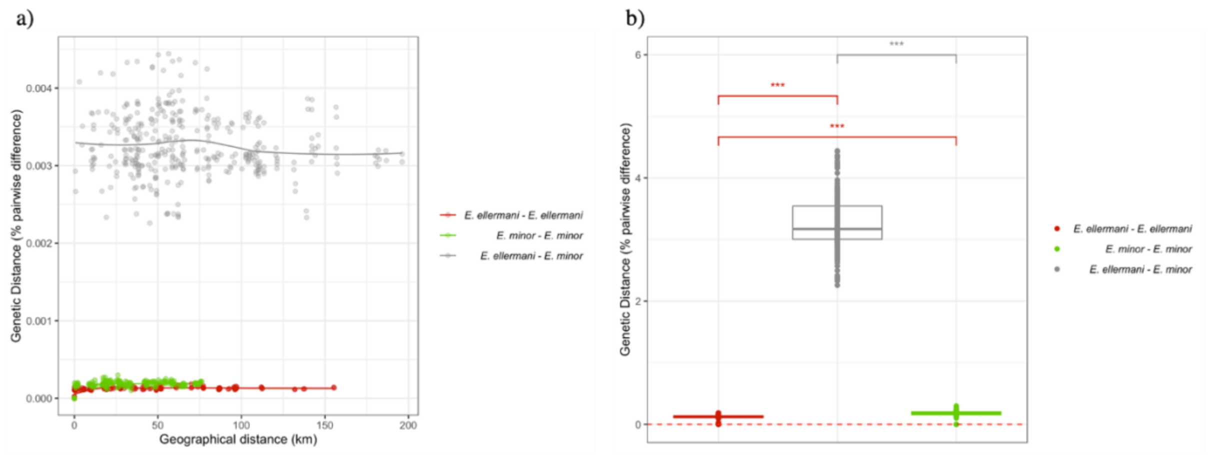


**Figure S13. Pairwise genomic differences and genomic covariance within and between *E. ellermani* and *E. minor*.** a) Isolation by distance within and between taxa. Genetic distance was measured as ‘percentage of genetic differences on variant sites’, calculated from the *N124.pop5.p4.min100* dataset (4.351 SNPs); b) Comparison of genetic distance within and between taxa.

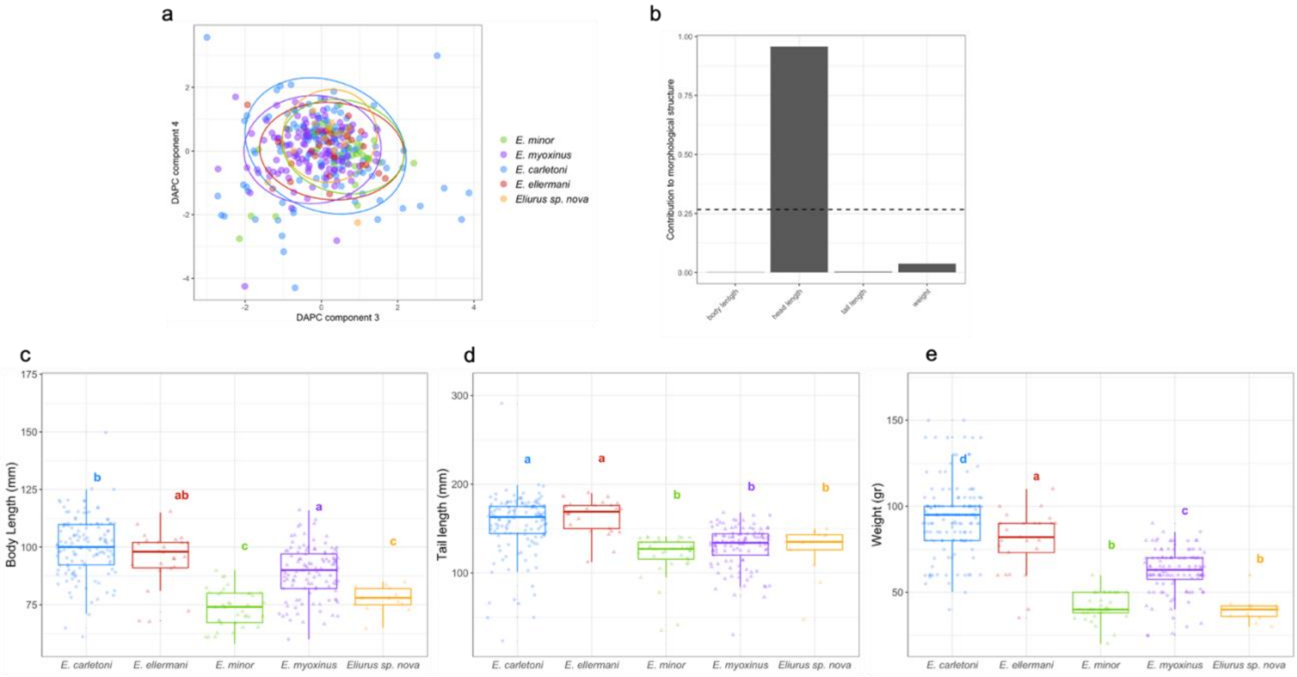


**Figure S14. Discriminant Analysis of Principal Components analysis on *Eliurus* taxa sampled across northern Madagascar.** a) DAPC plot obtained using the third and fourth discriminant components. c) loading plot of the first discriminant component shows that “head length” is the morphological variable that discriminate the most the five *Eliurus* taxa. The dashed line corresponds to the third quantile of the first component distribution and is used as default threshold value in *adegenet* for identifying significant variables. Panels d), e) and f) show the box plots with multiple comparison Tukey test results for body length, tail length and weight, respectively. Overall, the results show that the five *Eliurus* taxa can be partially distinguished using the four morphological variables here considered. It is worth to note that the new *Eliurus* taxa present significant differences across all variables with its sister taxa *E. myoxinus*

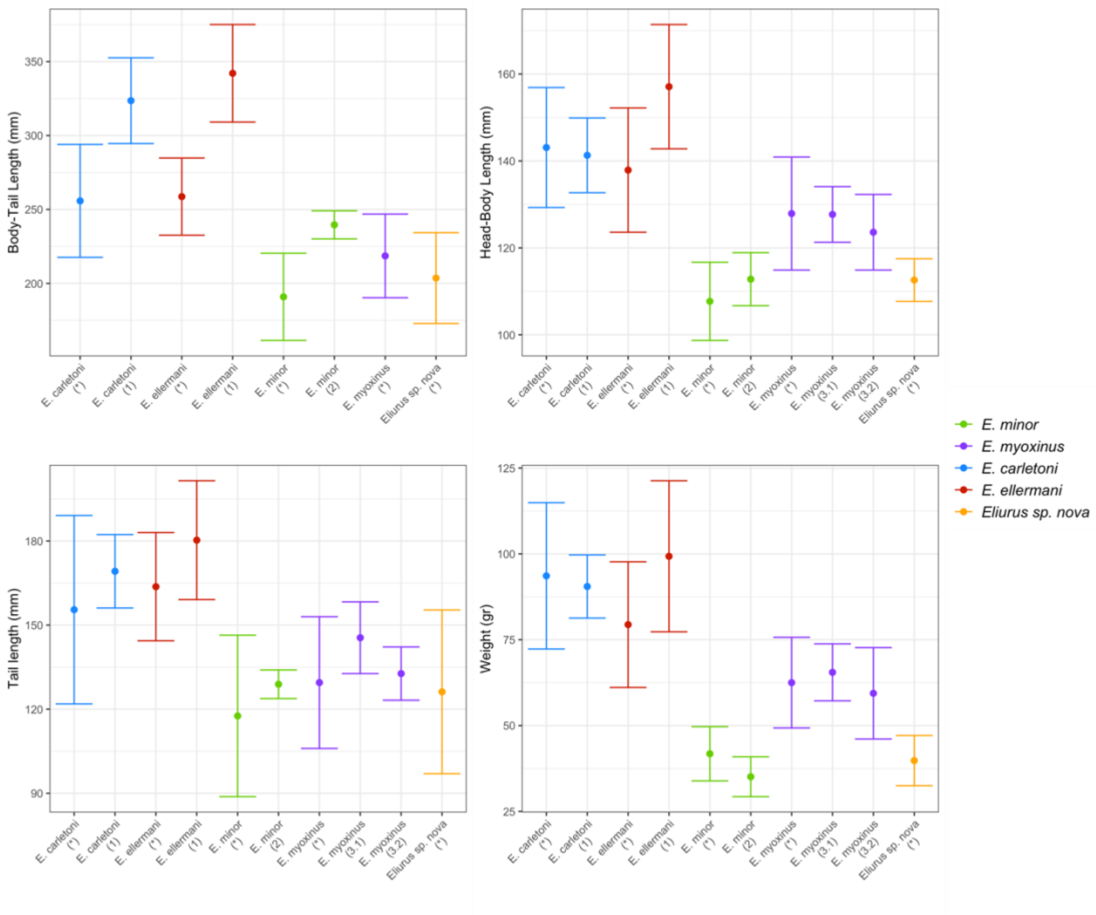


**Figure S15. Comparisons of morphological data from the present study with previously published data.** "*" indicates data from the present study; "1", "2" and "3" indicate data retrieved from Jansa et al., (2019), Goodman and Carleton, (1996) and Carleton et al., (2001), respectively. Code "3.1" and "3.2" are used to distinguish data for *E. myoxinus* collected in Analavelona and Ankarafanstika, respectively. The results reveal consistency between our and previously published data.

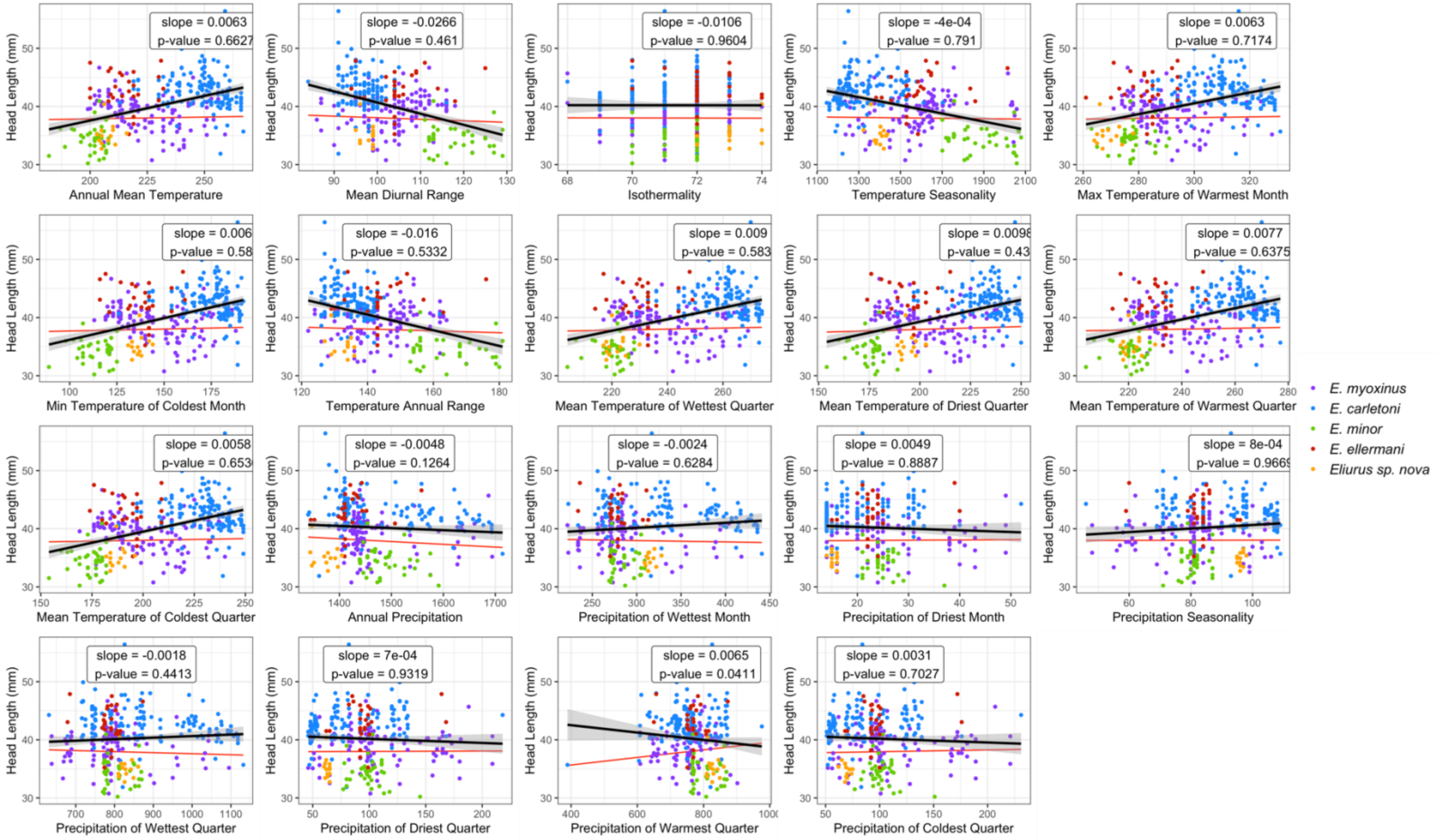


**Figure S16. PGLS analysis between head length and 19 bioclimatic variables.** Black line indicates linear regression without accounting for phylogenetic relatedness. Red line indicates linear regression based on PGLS analysis. The results show that after correcting for phylogenetic relatedness, any of the bioclimatic variables correlate with head length.

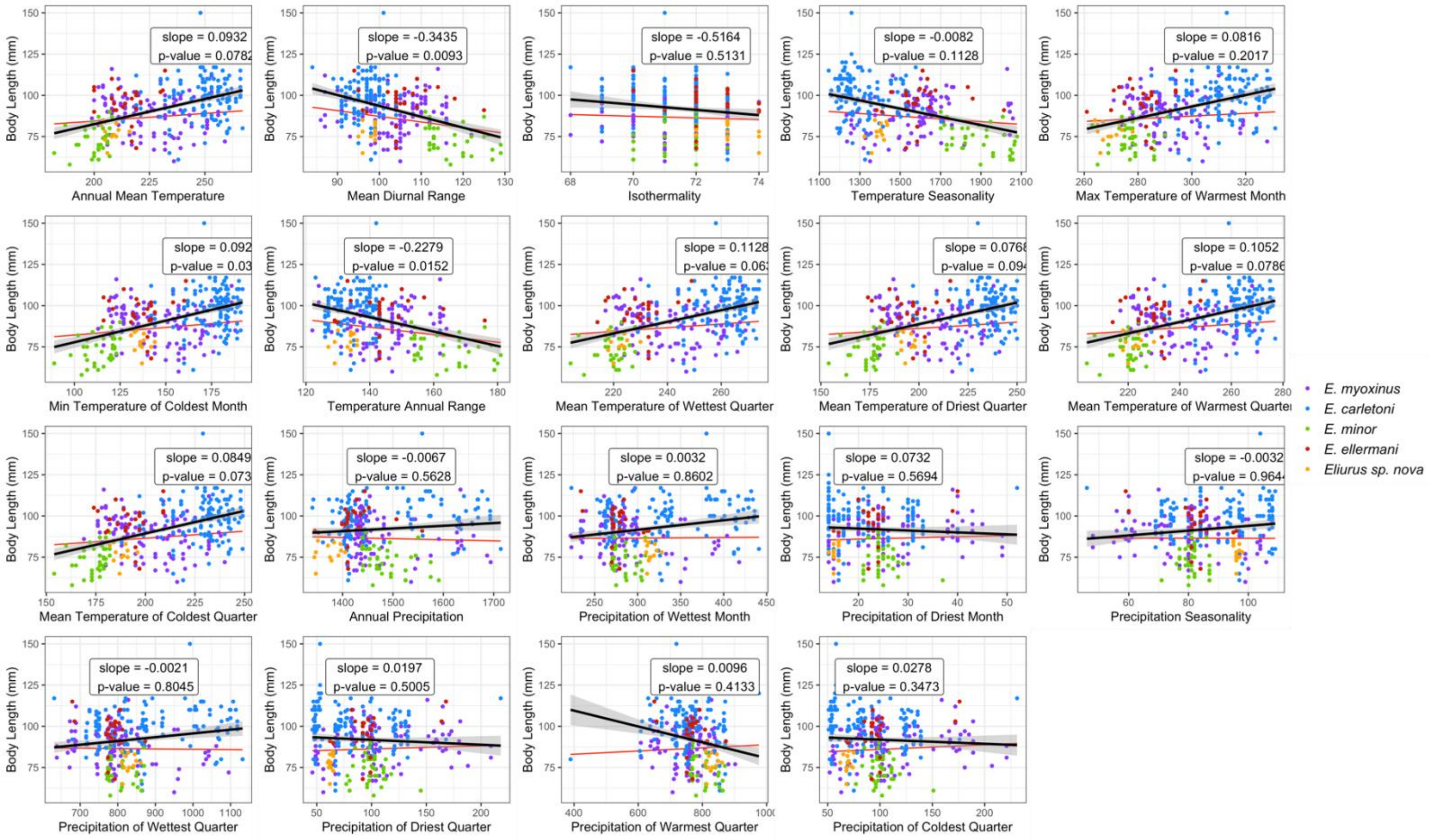


**Figure S17. PGLS analysis between body length and 19 bioclimatic variables.** Black line indicates linear regression without accounting for phylogenetic relatedness. Red line indicates linear regression based on PGLS analysis. The results show that after correcting for phylogenetic relatedness, 3 out of 19 bioclimatic variables show significant correlation with body length.

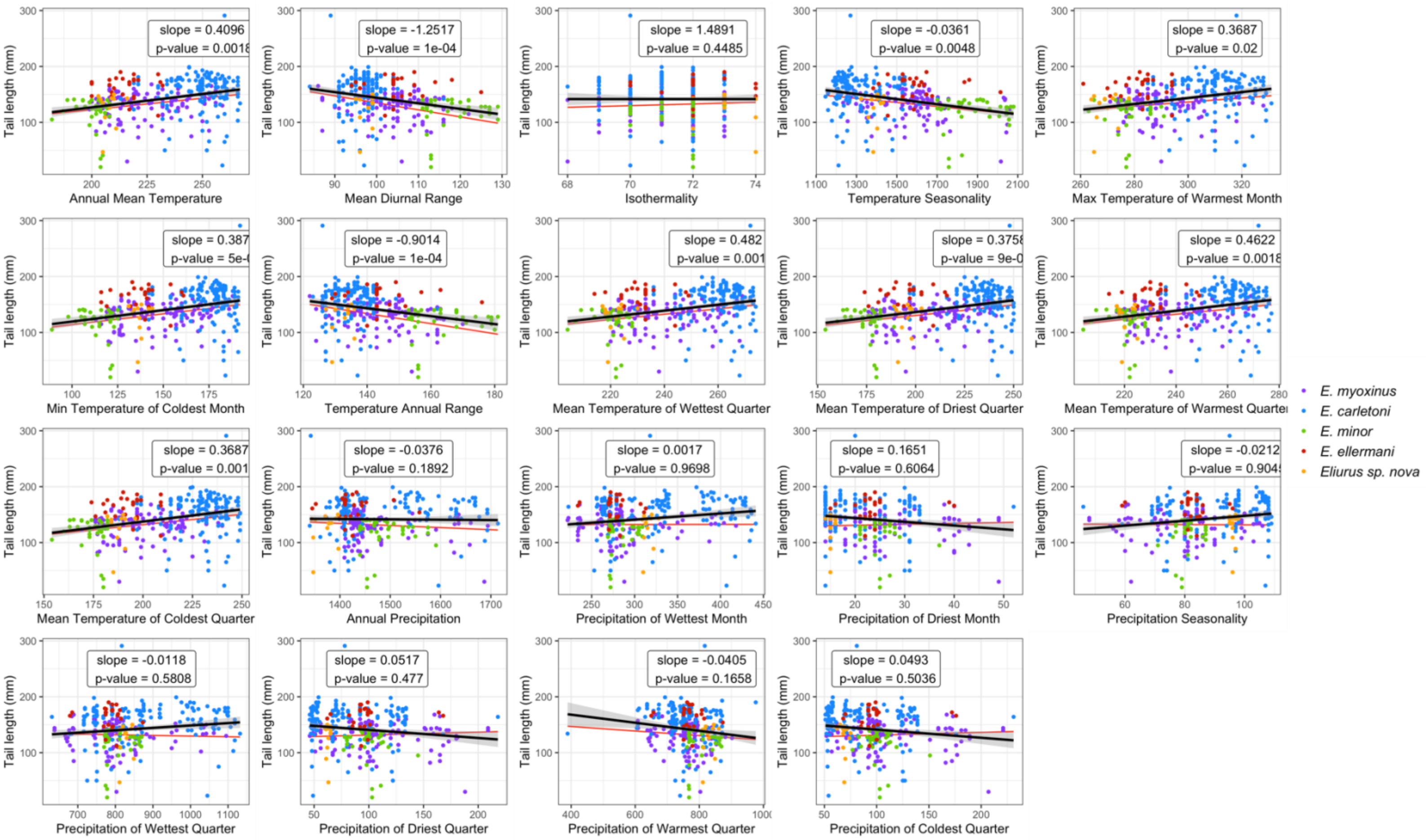


**Figure S18. PGLS analysis between tail length and 19 bioclimatic variables.** Black line indicates linear regression without accounting for phylogenetic relatedness. Red line indicates linear regression based on PGLS analysis. The results show that after correcting for phylogenetic relatedness, 10 out of 19 bioclimatic variables show significant correlation with tail length.

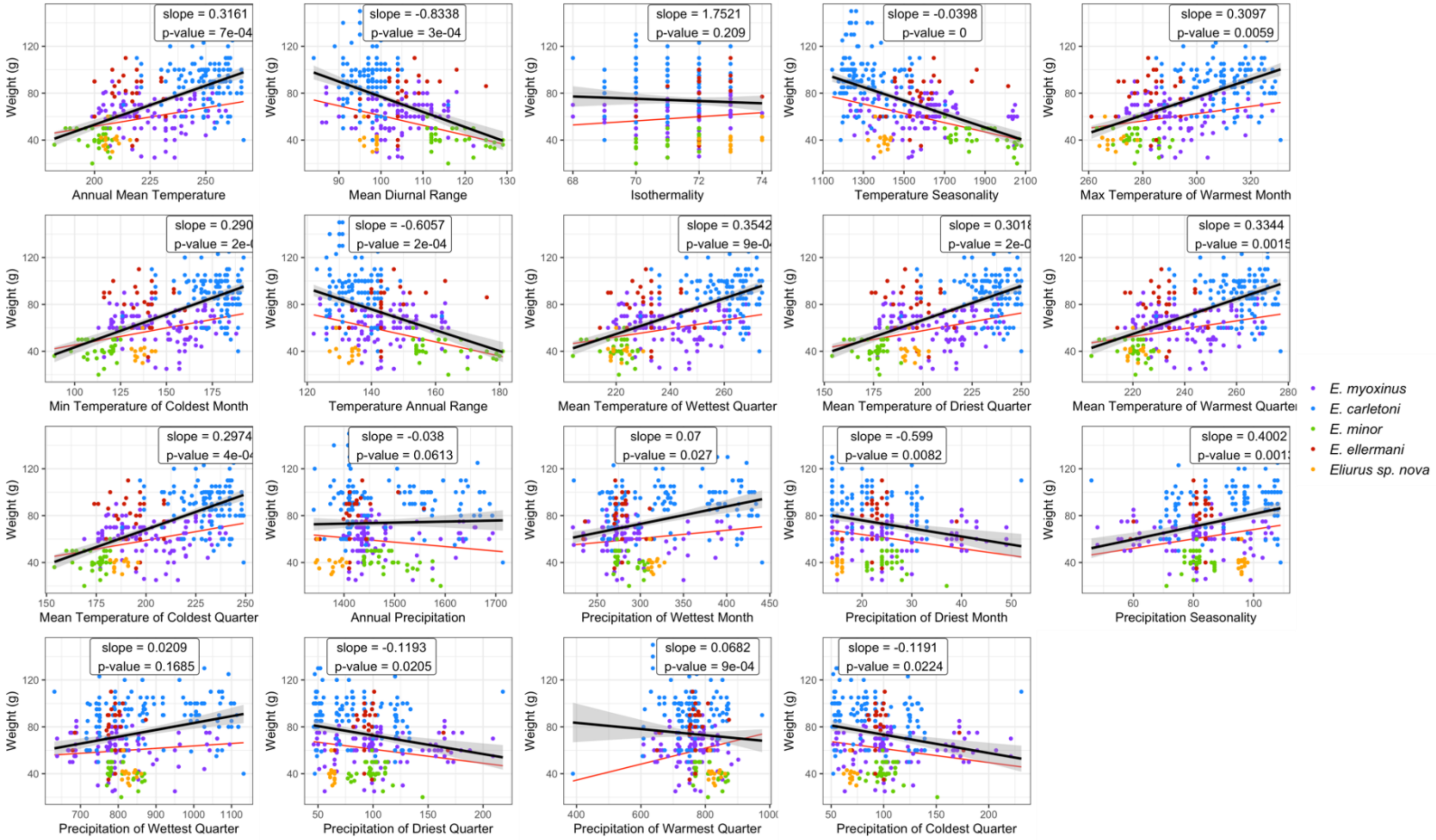


**Figure S19. PGLS analysis between weight and 19 bioclimatic variables.** Black line indicates linear regression without accounting for phylogenetic relatedness. Red line indicates linear regression based on PGLS analysis. The results show that after correcting for phylogenetic relatedness, 16 out of 19 bioclimatic variables show significant correlation with weight.

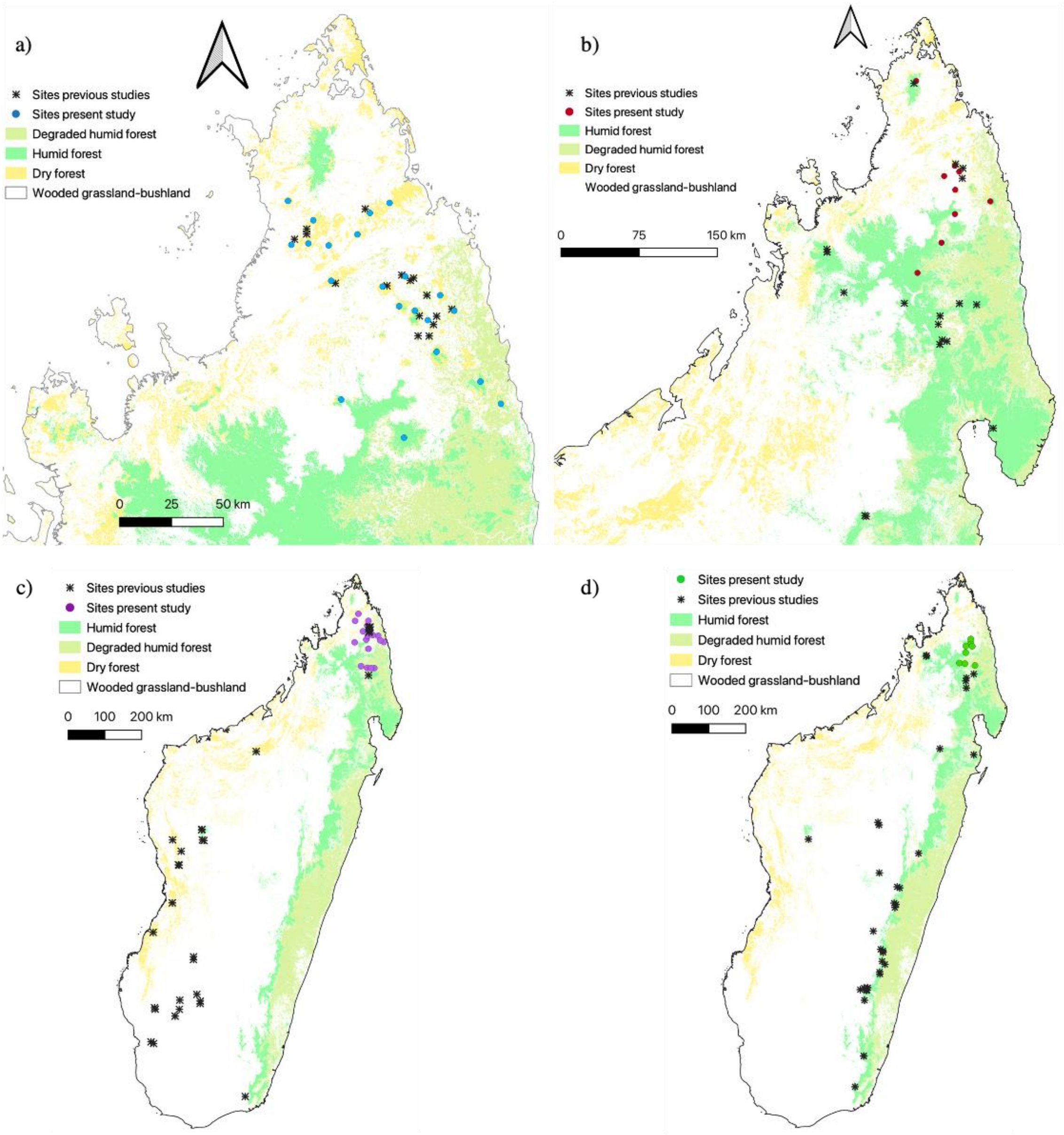


**Figure S20. Range distribution of the five *Eliurus* species identified in the present study.**

a) *E. carletoni*; b) *E. ellermani*; c) *E. myoxinus*; *E. minor*.

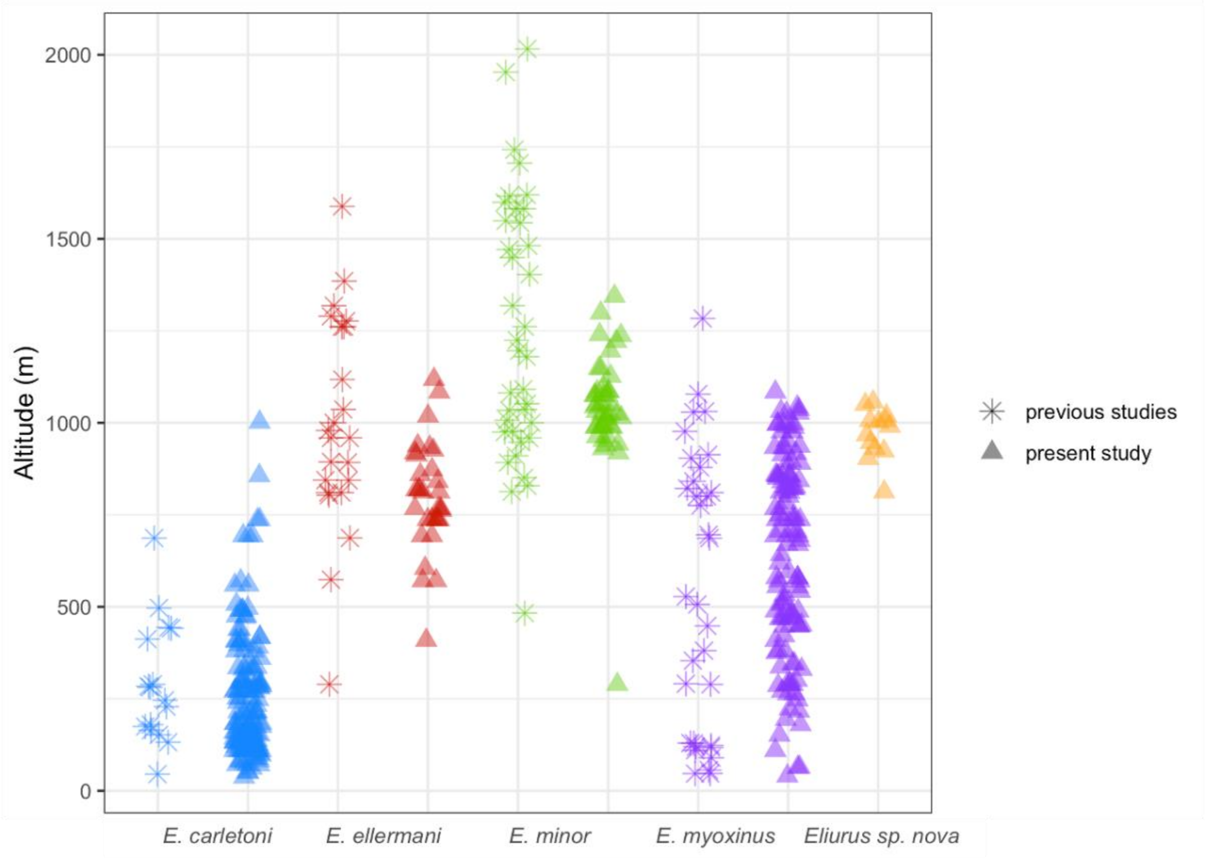


**Figure S21. Altitudinal range of *Eliurus* species sampled in northern Madagascar, across their entire distribution.** Significant differences in elevational range were identified for *E. carletoni*, *E. myoxinus*, *E. ellermani* and *E. minor* (p-value < 0.001), whereas no significant differences were detected between *E. ellermani* – *Eliurus sp. nova* (p-value = 0.78) and *E. minor* - *Eliurus sp. nova* (p-value = 0.14).
